# Comparative transcriptomics reveal a novel tardigrade specific DNA binding protein induced in response to ionizing radiation

**DOI:** 10.1101/2023.09.08.556854

**Authors:** M. Anoud, E. Delagoutte, Q. Helleu, A. Brion, E. Duvernois-Berthet, M. As, X. Marques, K. Lamribet, C. Senamaud, L. Jourdren, A. Adrait, S. Heinrich, G. Toutirais, S. Hamlaoui, G. Gropplero, I. Giovannini, L. Ponger, M. Gèze, C. Blugeon, Y. Coute, R. Guidetti, L Rebecchi, C. Giovannangeli, A. De Cian, J-P. Concordet

## Abstract

Tardigrades, microscopic animals found in virtually all ecosystems, are renowned for their remarkable ability to withstand extreme conditions. Recent studies have identified novel tardigrade specific protein families that aid in resistance to desiccation and ionizing radiation (IR). Notably, a tardigrade specific DNA binding protein called Dsup (for DNA damage suppressor) has been found to protect from X-ray damage in human cells and from hydroxyl radicals *in vitro*. However, Dsup has only been found in two species within the Hypsibioidea superfamily.

To better understand mechanisms underlying radio-resistance in the Tardigrada phylum, we first characterized DNA damage and repair in response to IR in the model species *Hypsibius exemplaris*. By analysis of phosphorylated H2AX, we demonstrated the induction and repair of DNA double-strand breaks after IR exposure. Importantly, the rate of single-strand breaks induced was roughly equivalent to that in human cells, suggesting that DNA repair plays a predominant role in the remarkable radio-resistance of tardigrades. In order to identify novel tardigrade specific genes involved, we next conducted a comparative transcriptomics across three species, *H. exemplaris*, *Acutuncus antarcticus* and *Paramacrobiotus fairbanksi*, the latter belonging to the Macrobiotoidea superfamily known to lack Dsup homologs. In all three species, many genes of DNA repair were among the most strongly overexpressed genes alongside a novel tardigrade specific gene, named *T*ardigrade *D*NA damage *R*esponse protein *1* (TDR1). We found that TDR1 protein interacts with DNA and forms aggregates at high concentration suggesting it may condensate DNA and act by preserving chromosome organization until DNA repair is accomplished. Remarkably, when expressed in human cells, TDR1 improved resistance to Bleomycin, a radiomimetic drug. Based on these findings, we propose that TDR1 is a novel tardigrade specific gene responsible for conferring resistance to IR. Our study sheds light on mechanisms of DNA repair helping to cope with high levels of DNA damage. Furthermore, it suggests that at least two tardigrade specific genes, respectively for Dsup and TDR1, have independently evolved DNA-binding functions that contribute to radio-resistance in the Tardigrada phylum.

## INTRODUCTION

Tardigrades are microscopic animals found in marine or freshwater environments, and in semi-terrestrial habitats such as moss, lichen and leaf litter. They are well known for their resistance to IR (Jönsson 2019) and extreme conditions like desiccation, freezing, and osmotic stress (Guidetti, Altiero, and Rebecchi 2011). With over 1400 species, they belong to the clade of ecdysozoans, which also includes nematodes and arthropods (Degma and Guidetti, 2023). Tardigrades share a highly conserved body plan, with a soft body protected by a cuticle, four pairs legs and a characteristic feeding apparatus. Tardigrades, however, can differ in their resistance to extreme conditions. For example, *Ramazzottius oberhaeuseri* withstands extremely rapid desiccation while *Hypsibius dujardini* only survives gradual dehydration (Wright 1989), and the freshwater *Thulinius ruffoi* is not resistant to desiccation (Kondo et al. 2020). Many species across the Tardigrada phylum can tolerate irradiation doses higher than 4000 Gy (Hashimoto and Kunieda 2017), but some are less tolerant, like *Echiniscoides sigismundi*, which has an LD50 at 48 hours of 1000 Gy (Jönsson et al. 2016). However, the doses compatible with maintenance of fertility seem much lower, e.g. the maximum is 100 Gy for *H. exemplaris* (Beltrán-Pardo et al. 2015). Due to challenges in rearing tardigrades in the laboratory (Altiero and Rebecchi 2001), the maintenance of fertility has seldom been investigated and some species might remain fertile at higher doses.

Understanding the genes involved in tardigrade resistance to IR is essential to unraveling the mechanisms of their exceptional resilience. Systematic comparison of whole genome sequences has suggested that tardigrades have one of the highest proportions of gene gain and gene loss among metazoan phyla (Guijarro-Clarke, Holland, and Paps 2020). Several novel, tardigrade specific genes have indeed been involved in resistance to desiccation including CAHS, MAHS, SAHS, and AMNP gene families (Hesgrove and Boothby 2020; Arakawa 2022; Yamaguchi et al. 2012; S. Tanaka et al. 2015; Yoshida et al. 2022). For resistance to IR, the tardigrade specific gene Dsup (for DNA damage suppressor) has been discovered in *Ramazzottius varieornatus*. Dsup encodes an abundant chromatin protein that increases resistance to X-rays when expressed in human cells (Hashimoto et al. 2016). *In vitro* experiments have shown that DNA damage induced by hydroxyl radicals was reduced when Dsup was added to nucleosomal DNA (Chavez et al. 2019), indicating DNA protection by Dsup. However, it is not yet possible to inactivate genes with CRISPR-Cas9 in tardigrades (Goldstein 2022) and direct evidence for the importance of Dsup in radio-resistance of tardigrades is still lacking. Interestingly, the presence of resistance genes differs across tardigrade genomes (Arakawa 2022). While AMNP genes are found in both classes of tardigrades, Heterotardigrada and Eutardigrada, CAHS, SAHS, and MAHS genes are only found in Eutardigrada, and Dsup appears restricted to the Hypsibioidea superfamily of Eutardigrada (Arakawa 2022). However, given the range of species demonstrated to be radio-resistant across the phylum (Hashimoto and Kunieda 2017), it seems likely that additional tardigrade specific genes are involved in tardigrades’ radio-resistance.

In addition to tardigrades, other animals display exceptional resistance to IR including rotifers, nematodes and larvae of *Polypedilum vanderplanki* midges, all surviving doses more than 100 times higher than humans. Recent studies have begun to shed light on the mechanisms involved (Ujaoney et al. 2024). In addition to DNA protection, DNA repair may also help maintain genome integrity upon irradiation. In rotifers, the rate of DNA double-strand breaks is equivalent to that in human cells (Gladyshev and Meselson 2008), showing that DNA repair, rather than DNA protection, plays a predominant role in their radio-resistance. Furthermore, it was recently found that genes of DNA repair are upregulated in response to IR in rotifers and *P. vanderplanki* larvae (Moris et al. 2023; Ryabova et al. 2017). In rotifers, upregulation of a DNA ligase gene acquired by horizontal gene transfer may be essential to radio-resistance (Nicolas et al. 2023). In prokaryotes, radio resistance has been investigated in the bacterium *Deinococcus radiodurans*, showing highly efficient DNA repair in response to the high levels of DNA damage induced by high doses of IR and the contribution of *D. radiodurans* DNA repair genes (Timmins and Moe 2016). Previous studies have suggested that upregulation of DNA repair genes may also play a role in radio-resistance of tardigrades: irradiation with IR increases expression of Rad51, the canonical recombinase of homologous recombination (HR), in *Milnesium inceptum* (Beltrán-Pardo et al. 2013), and regulation of DNA repair genes was observed in *R. varieornatus* (Yoshida et al. 2021).

To improve our understanding of resistance to IR in tardigrades, we sought to characterize DNA damage and repair after irradiation and to identify novel tardigrade specific genes involved in resistance to IR. For this purpose, we first examined the kinetics of DNA damage and repair after IR in the model species *H. exemplaris*. This species was chosen due to its ease of rearing in laboratory conditions and its known genome sequence. Additionally, to identify novel genes involved in resistance to IR, we analyzed gene expression in response to IR in *H. exemplaris* and two additional species, *Acutuncus antarcticus,* from the Hypsibioidea superfamily (Giovannini et al. 2018), and *Paramacrobiotus fairbanksi* of the Macrobiotoidea superfamily (Guidetti et al. 2019). Together with multiple DNA repair genes, a tardigrade specific gene, which we named *T*ardigrade *D*NA damage *R*esponse gene *1* (TDR1) was strongly upregulated in response to IR in all three species analyzed. Further analyses in *H. exemplaris*, including differential proteomics and Western blots, showed that TDR1 protein is present and upregulated. *In vitro* experiments demonstrated that recombinant TDR1 interacts with DNA and forms aggregates with DNA at high concentrations. Importantly, when expressed in human cells, TDR1 reduced the number of phospho-H2AX foci induced by Bleomycin, a DNA damaging drug used as a radiomimetic. These findings show the importance of DNA repair in radio-resistance of tardigrades and suggest that TDR1 is a novel tardigrade specific DNA binding protein involved in DNA repair after exposure of tardigrades to IR.

## RESULTS

### Double- and single-strand breaks are induced and repaired after exposure of *H. exemplaris* to IR

IR causes a variety of damages to DNA such as nucleobase lesions, single- and double-strand breaks (Téoule 1987). In eukaryotes, from yeast to humans, phosphorylation of H2AX is a universal response to double-strand breaks (DSBs) and an early step in the DNA repair process (Fernandez-Capetillo et al. 2004). To investigate DSBs caused by IR, we generated an antibody against phosphorylated H2AX of *H. exemplaris* (Supp Figure 1). *H. exemplaris* tardigrades were exposed to either 100 or 1000 Gy of ^137^Cs γ-rays, which are known to be well tolerated by this species (Beltrán-Pardo et al. 2015). We analyzed phospho-H2AX in protein extracts of *H. exemplaris* collected at 30 min, 4h, 8h30, 24h and 73 h after irradiation. For both 100 and 1000 Gy doses, phospho-H2AX was detected at 30 min after irradiation, reached its peak levels at 4h and 8h30 and then gradually decreased (Figure 1a). Irradiation was also performed with an accelerated electron beam, which delivered identical doses in much shorter times, 1000 Gy in 10 min instead of 1h for the ^137^Cs source, in order to better appreciate the early peak of phospho-H2AX. A peak of phospho-H2AX was detected at 4h and a similar, gradual decrease was observed (Supp Figure 2a). Next, we performed whole mount immunolabeling of tardigrades and observed intense, ubiquitous phospho-H2AX labeling in nuclei 4h after 100 Gy irradiation, which had significantly decreased 24h later (Figure 1b). This suggests irradiation impacts all adult cells and indicates efficient DNA repair by 24h after 100 Gy irradiation, consistent with the results of Western blot analysis. After 1000 Gy irradiation, the intense signal detected at 4h had decreased in most nuclei at 24h but it persisted at high intensity specifically in gonads (Supp Figure 2b-c). The finding of persistent DSBs in gonads at 72h after 1000 Gy likely explains why *H. exemplaris* no longer lay eggs and become sterile after exposure to 1000 Gy (Beltran-Pardo 2005). In order to investigate DNA synthesis taking place after irradiation, we incubated tardigrades with the thymidine nucleotide analog EdU (Gross et al. 2018). Using confocal microscopy, we could detect DNA synthesis in replicating intestinal cells of control animals, as previously shown by (Gross et al. 2018). In contrast, we could not detect any specific signal in irradiated tardigrades compared to controls, suggesting (i) that DNA synthesis induced during DNA repair remained at low, undetectable levels and (ii) that dividing intestinal cells detected in control animals were irreversibly damaged by the 1000 Gy irradiation (Supp Figure 2d). Together, these results demonstrate the dose-dependent induction and repair of DSBs in response to IR. Phospho-H2AX immunolabeling experiments also suggested that 1000 Gy induces irreversible damage in the gonads and dividing intestinal cells.

**Figure 1.**
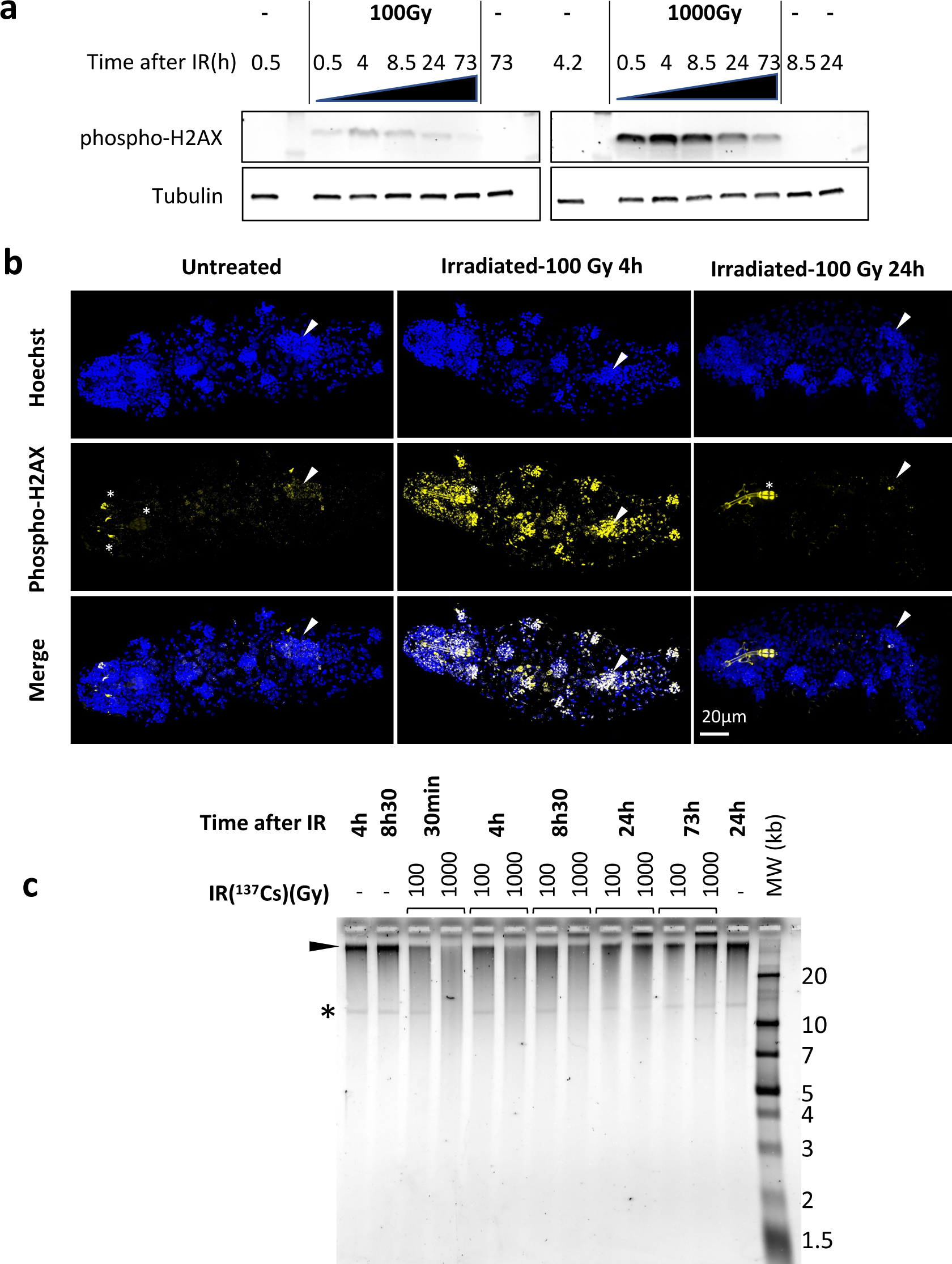
Analysis of DNA damage and repair in *Hypsibius exemplaris* after γ-ray irradiation. **(a)** Analysis of phospho-H2AX expression after exposure of *H. exemplaris* to IR. Western blot analysis with in-house antibody against phosphorylated *H. exemplaris* H2AX (anti-phospho-H2AX) at indicated time points after irradiation of tardigrades with indicated dose of γ-ray irradiation. Phospho-H2AX levels were normalized by total α−tubulin expression levels and quantification is provided in Supp Figure 2a. (-) lanes show extracts from control tardigrades processed in parallel to irradiated tardigrades at indicated time points post-irradiation. **(b)** Analysis of phospho-H2AX expression in whole mount *H. exemplaris* after exposure to 100 Gy. Tardigrades were exposed to 100 Gy, fixed with 4% PFA at 4 and 24h post irradiation, immunolabeled with anti-phosphoH2AX antibody and anti-rabbit IgG conjugated to Alexa488 and visualized by confocal microscopy using the Airyscan2 module. Maximum projection of confocal Z-stack are shown. Images at different time points were taken with identical settings so that signal intensity could be compared. Upper panel shows Hoechst staining of nuclei (in blue). Arrowhead indicates position of the gonad (revealed by intense Hoechst and larger nuclei signal). The gonad exhibits intense labeling phospho-H2AX at 4h which is no longer detected at 24h, showing efficient DNA repair consistent with preservation of the capacity to lay eggs and reproduce after 100 Gy IR (Beltran-Pardo et al, 2005). * indicates autofluorescence of bucco-pharyngeal apparatus. Scale bar 20µm **(c)** Analysis of single-strand breaks by denaturing agarose gel electrophoresis of DNA isolated from ∼8000 *H. exemplaris* at indicated time points post-irradiation (100Gy or 1000Gy γ-rays from ^137^Cs source). (-) indicates DNA from control, non-irradiated tardigrades collected and processed in parallel to treated samples from indicated time points. MW corresponds to the Molecular Weight ladder. * indicates a discrete band of single stranded DNA detected in *H. exemplaris* genomic DNA. Arrowhead indicates high molecular weight single-stranded DNA that is not resolved by agarose gel electrophoresis. (-) lanes show DNA from control tardigrades processed in parallel to irradiated tardigrades at 4h or 8h30 post-irradiation as indicated.

Next, we assessed the physical integrity of genomic DNA at several time points after irradiation. Samples from Figure 1a were run in native agarose gels and irradiated samples were found to be indistinguishable from non-irradiated controls (Supp Figure 2e), showing DSBs and the resulting DNA fragmentation could not be detected in this experimental setting. Single-strand breaks (SSBs) were evaluated by migrating DNA samples in denaturing agarose gels (Figure 1c). DNA from control, untreated tardigrades appeared as a predominant band running above the 20 kb marker with a smear. The smear, likely due to the harsh extraction conditions needed for tardigrade cuticle lysis, extended down between 20 and 10 kb markers where a discrete band, of unknown origin, could be detected (Figure 1c). At 30 min after 1000 Gy irradiation, intensity of the high molecular weight band was drastically reduced, and DNA detected in the smear between 10 and 20 kb was strongly increased. In addition, the discrete band could no longer be detected. This clearly indicates that 1000 Gy IR induces high rates of SSBs. Considering that the majority of DNA fragments detected had a size of 10 to 20 kb and that the discrete band of 10-20 kb could no longer be detected, we can roughly evaluate that there is approximately 1 SSB every 10 to 20 kb. This corresponds to induction of SSBs at a rate of 0.05-0.1 SSB/Mb/Gy. Between 4h and 24h, the DNA migration profile was progressively restored and between 24h and 73h, it was identical to controls. Similar results were observed with the 100 Gy dose (Figure 1c). However, compared to 1000 Gy, the changes observed were not as marked and the discrete 10-20 kb band could always be detected, indicating SSBs were induced at lower rates. These results indicate that SSBs are inflicted by IR in a dose-dependent manner, roughly estimated to 0.05-0.1 SSB/Mb/Gy, and progressively repaired within the next 24 to 73h (Figure 1c).

### *H. exemplaris* strongly overexpresses canonical DNA repair genes as well as RNF146 and TDR1, a novel tardigrade specific gene, in response to IR

To examine the gene expression changes associated with tardigrade response to IR, we performed RNA sequencing of *H. exemplaris* collected 4h after irradiation. The analysis revealed that 421 genes were overexpressed more than 4-fold (with an adjusted p-value < 0.05) including 120 overexpressed more than 16-fold (Figure 2a, Supp Table 1, Supp Figure 3a). The Gene Ontology analysis of overexpressed genes highlighted a strong enrichment of DNA repair genes (Figure 2a, Supp Figure 4). In particular, genes for both major pathways of DNA double strand break repair, HR and NHEJ, were among the most strongly stimulated genes. Examples are genes for RAD51 and MACROH2A1 in HR (Khurana et al. 2014; Baumann and West 1998) and XRCC5 and XRCC6 in NHEJ (Figure 2a). The gene for POLQ, the key player of the alternative end joining pathway of DNA double strand break repair (Mateos-Gomez et al. 2015), was also strongly upregulated (Figure 2a). Also notable among most strongly overexpressed genes were genes for XRCC1, PNKP and LIG1 in Base Excision repair (Whitehouse et al. 2001; Krokan and Bjørås 2013), along with genes for PARP2 and PARP3, which catalyze PARylation of many DNA repair proteins (Pascal 2018) and RNF146 (Figure 2a). Interestingly, RNF146 is a ubiquitin ligase that has been reported to be important for tolerance to IR in human cells by targeting PARylated XRCC5, XRCC6 and XRCC1 for degradation (Kang et al. 2011). Our results suggest that RNF146 upregulation could contribute to the remarkable resistance of tardigrades to IR.

**Figure 2.**
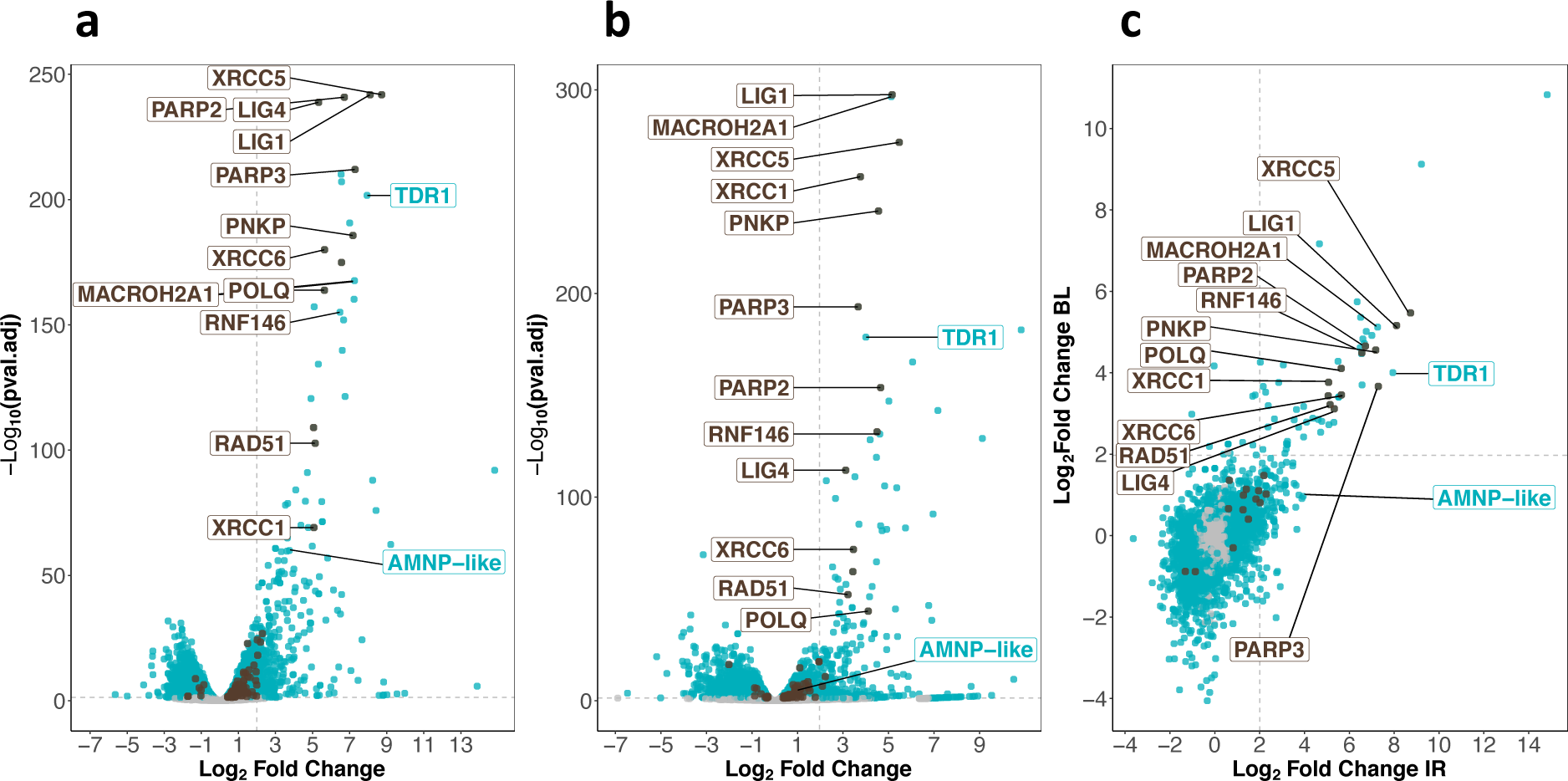
Transcriptomic response of *Hypsibius exemplaris* to IR and Bleomycin. **(a)** and **(b)** Volcano plots representing Log_2_ Fold Change and adjusted p-value (−log base 10) of RNA levels between *H. exemplaris* irradiated with 1000Gy γ-rays and untreated controls *(n=3)* **(a)** and between *H. exemplaris* treated with 100 µM Bleomycin for 4 days and untreated controls *(n=3)* **(b).** The vertical dotted lines indicate the Log_2_Fold Change value of 2 (Fold Change of 4). **(c)** Correlation between Log_2_ Fold Change after exposure to IR and after Bleomycin (BL) treatment for abundant transcripts (with baseMean>500 after DESeq2 analysis). The horizontal and vertical dotted lines indicate the Log_2_Fold Change value of 2 (Fold Change of 4). Blue dots represent transcripts with a Log_2_ Fold Change with an adjusted p-value p<0.05. Brown dots indicate transcripts of DNA repair genes (based on KEGG DNA repair and recombination gene group ko03400) that have a Log_2_ Fold Change with adjusted p-value p<0.05. Grey dots represent transcripts with a Log_2_ Fold Change with an adjusted p-value p>0.05. Brown labels indicate representative strongly upregulated genes of DNA repair. Blue labels indicate two tardigrade specific genes induced in response to IR: the TDR1 gene identified in this work, and the AMNP-like gene (BV898_10264), a member of the family of AMNP/g12777-like genes upregulated in response to desiccation and UVC (Yoshida et al 2022).

Among overexpressed genes, we also observed AMNP gene family members (Yoshida et al. 2022) (one representative was labeled AMNP-like, Figure 2a). AMNP genes encode recently discovered tardigrade specific Mn-Peroxidases which are overexpressed in response to desiccation and UVC in *R. varieornatus* (Yoshida et al. 2022). AMNP gene g12777 was shown to increase tolerance to oxidative stress when expressed in human cells (Yoshida et al. 2022). Based on our results, it is possible that AMNP genes such as the AMNP-like gene identified here could contribute to resistance to IR by increasing tolerance to the associated oxidative stress.

In parallel, we also determined the transcriptomic response of *H. exemplaris* to Bleomycin, a well-known radiomimetic drug (Bolzán and Bianchi 2018) (Figure 2b, Supp Table 2, Supp Figure 3b). In preliminary experiments, we found that *H. exemplaris* tardigrades survived for several days in the presence of 100 μM Bleomycin, suggesting that *H. exemplaris* could resist chronic genotoxic stress. We hypothesized that key genes of resistance to acute genotoxic stress induced by IR would also be induced by Bleomycin treatment. As expected, the correlation between highly expressed genes after IR and after Bleomycin treatment (with baseMean > 500, Supp Figure 3 and Supp Table 3) was strong for most upregulated DNA repair genes such as XRCC5, XRCC6, PARP2, PARP3, XRCC1, LIG4, LIG1 and RNF146 (Figure 2c). Importantly, in addition to DNA repair genes, several genes of unknown function were also strongly overexpressed in both conditions and considered as promising candidates for a potential role in resistance to IR. One such gene, which we named TDR1 (for Tardigrade DNA damage Response 1), was chosen for further investigation. ONT long read sequencing and cDNA cloning of TDR1 allowed us to determine the predicted TDR1 protein sequence which is 146 amino acids long (Supp Figure 5, Supp Table 4). We observed that the current genome assembly predicts a partially truncated TDR1 protein sequence, BV898_14257, due to an assembly error (Supp Figure 5). Our BLAST analysis against NCBI nucleotide non-redundant database suggested that TDR1 is a novel tardigrade specific gene as no homolog could be found in any other ecdysozoan (Supp Table 5).

### Analysis of proteomic response to IR in *H. exemplaris* confirms overexpression of TDR1

We next examined whether stimulation of gene expression at the RNA level led to increased protein levels and in particular, whether TDR1 protein was indeed overexpressed. For this purpose, we first generated specific antibodies to *H. exemplaris* TDR1, XRCC5, XRCC6, and Dsup proteins. Protein extracts from *H. exemplaris* treated with Bleomycin for 4 days or 1000Gy of γ-rays at 4h and 24h post-irradiation were compared to untreated controls. The apparent molecular weight of the TDR1 protein detected on Western blots was consistent with the expected 16 kD predicted from the 146 amino acid long sequence (Figure 3a). Remarkably, similar to phospho-H2AX, TDR1 was only detected after the induction of DNA damage (Figure 3a). XRCC5 and XRCC6 protein levels were also stimulated by both Bleomycin and IR treatments, although the fold stimulation was much lower than at the RNA level (Figure 3a, Supp Figure 6). Furthermore, we checked expression of *He*-Dsup homolog in *H. exemplaris* (Chavez et al. 2019), which remained constant at the RNA level (see BV898_01301, Supp Tables 1-2), and found that it also remained stable at the protein level after the induction of DNA damage (Figure 3a).

**Figure 3.**
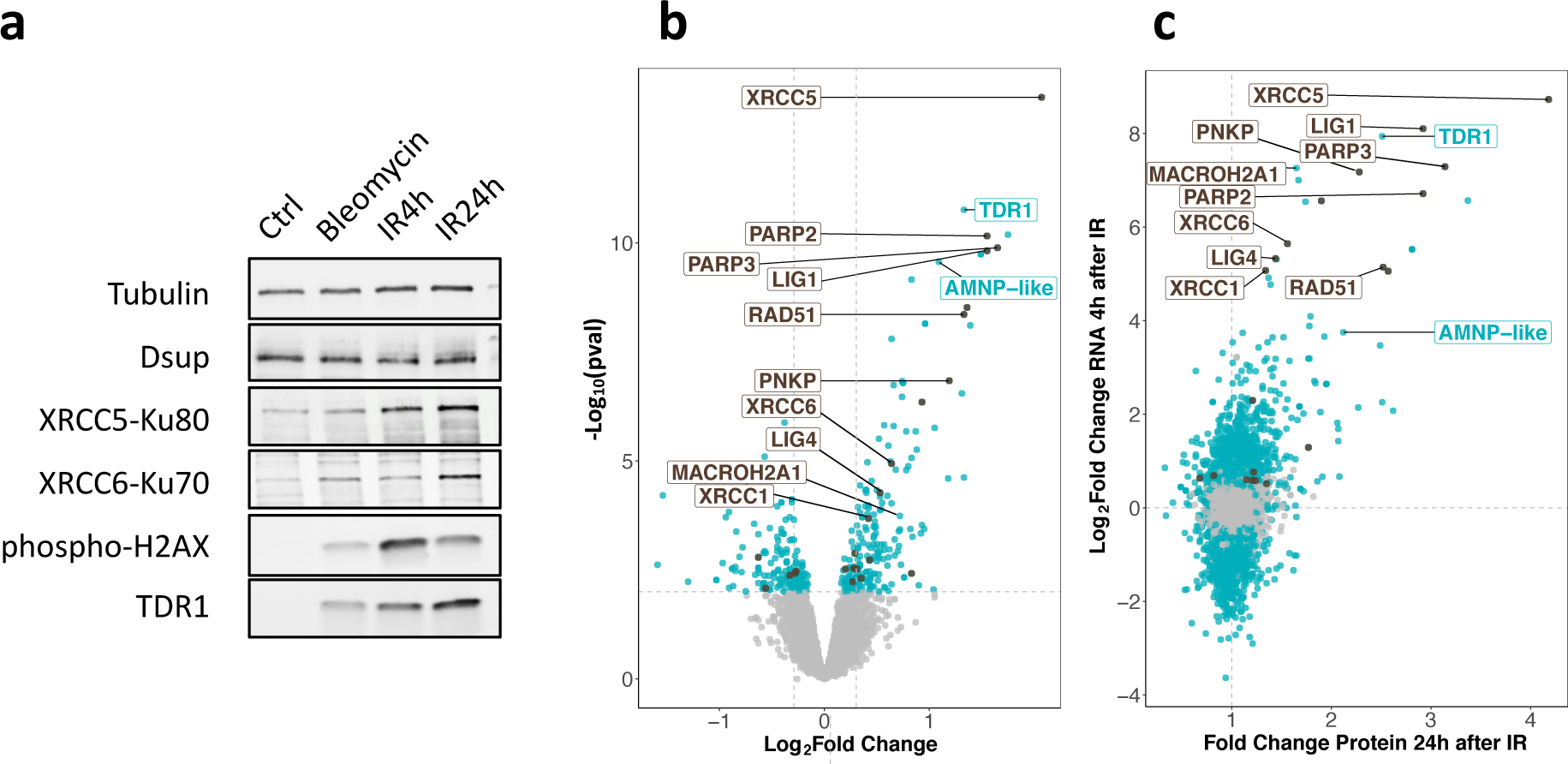
Changes in protein expression in *Hypsibius exemplaris* after exposure to IR. **(a)** Western blot analysis of *He-*TDR1, *He-*XRCC5, *He-*XRCC6 (among the most strongly stimulated genes at the RNA level) and *He-*Dsup (not stimulated at the RNA level) in irradiated *H. exemplaris* tardigrades Control, untreated *H. exemplaris* (Ctrl) or *H. exemplaris* treated with 100µM Bleomycin for 4 days, or with 1000Gy γ-rays and extracts prepared at indicated times post-irradiation (IR4h and IR24h). Alpha-tubulin was used for normalization and phospho-H2AX for showing induction of DNA double-strand breaks. Quantification of 4 independent experiments are shown in Supp Figure 6c. Molecular weight marker present in uncropped Western blots (Supp Figure 16b) is consistent with the expected 16kDa size of TDR1. **(b)** Volcano plot representing Log_2_ Fold Change and −Log_10_(limma p-value) of proteins between *H. exemplaris* 24h post-irradiation with 1000Gy γ-rays and untreated control animals *(n=4)*. Blue dots represent proteins with a Log_2_ Fold Change with a −Log_10_(limma p-value)>=2. Brown dots represent DNA repair proteins (based on KEGG DNA repair and recombination gene group ko03400) with −Log_10_(limma p-value)>=2. Grey points represent proteins with Log_2_ Fold Change with −Log10(limma p-value)<2 and the vertical grey lines delimit Log2(FC)>0.3 or <-0.3. Brown labels indicate representative strongly upregulated genes of DNA repair. Blue labels indicate two tardigrade specific genes induced in response to IR: the TDR1 gene identified in this work, and the AMNP-like gene (BV898_10264), a member of the family of AMNP/g12777-like genes upregulated in response to desiccation and UVC (Yoshida et al 2022). **(c)** Correlation between Fold Changes of protein levels 24h post-irradiation with 1000 Gy (as measured in (b)) and Log_2_FoldChange of RNA levels 4h post-irradiation (as measured in Figure 2a).

To ensure that the observed stimulation was due to new protein synthesis, we treated tardigrades with the translation inhibitor cycloheximide before irradiation (Supp Figure 7a). As expected, no increase in TDR1, XRCC5 or XRCC6 protein levels could be detected after irradiation in extracts from animals treated with cycloheximide (Supp Figure 7b). In particular, TDR1 protein could not be detected when animals were treated with cycloheximide, further confirming that TDR1 is strongly overexpressed in response to IR.

To further extend the analysis of the protein-level response to IR, we conducted an unbiased proteome analysis of *H. exemplaris* at 4h and 24h after irradiation and after Bleomycin treatment using mass spectrometry-based quantitative proteomics. More than 5600 proteins could be detected in all conditions (Supp Table 6). Among them, 58, 266 and 185 proteins were found to be differentially abundant at 4h post-irradiation, 24h post-irradiation and after Bleomycin treatment, respectively compared to control tardigrades (Log_2_Fold Change> 0.3 and limma p-value < 0.01, leading to a Benjamini-Hochberg FDR < 3%, Figure 3b, Table 1 and Supp Table 6). We observed a good correlation between stimulation at RNA and protein levels (Figure 3c). It is worth noting that the fold changes observed for proteins were smaller than those obtained for mRNAs, possibly due to the use of an isobaric multiplexed quantitative proteomic strategy known to compress ratios (Hogrebe et al. 2018). For strongly overexpressed canonical DNA repair genes discussed above, we confirmed significantly increased protein levels in response to IR (Figure 3b). RNF146, in contrast, could not be detected, likely due to limited sensitivity of our mass spectrometry-based quantitative proteomics. Importantly, despite the small size of the predicted TDR1 protein, we detected four different TDR1-related peptides, providing direct evidence of strong TDR1 over-expression in response to IR (Supp Table 6).

**Table 1:** Proteomic analysis metrics: numbers of differentially expressed (DE) proteins (with limma p-value < 0.01 and Log_2_FoldChange<-0.3 or >0.3) for each indicated condition in *Hypsibius exemplaris.* The numbers of tardigrade specific DE proteins are also indicated. 9 tardigrade specific DE proteins were common to the 3 conditions, the corresponding list is provided in Supp Table 10. Tardigrade specific proteins are defined as detailed in the Methods section.

| 1000 Gy $\gamma$ Irradiation vs Control | IR4h vs Ctrl | IR24h vs Ctrl | Bleomycin vs Ctrl |
| --- | --- | --- | --- |
| **Total number of identified proteins** |  | **5625** |  |
| **DE Proteins** | **58** | **266** | **185** |
| DE Proteins Up | 42 | 168 | 128 |
| DE Proteins Down | 16 | 98 | 57 |
| *DE proteins in the 3 conditions* |  | **36** |  |
| **Tardigrade specific DE proteins** | **13** | **61** | **70** |
| Tardigrade specific DE proteins Up | 11 | 52 | 47 |
| Tardigrade specific DE proteins Down | 2 | 9 | 23 |
| *Tardigrade specific DE proteins in the 3 conditions* |  | **9** |  |

### Conservation of TDR1 and transcriptional response to IR in other tardigrade species

To gain insight into the importance of the upregulation of TDR1 and DNA repair genes in resistance to IR, we chose to investigate its conservation in other tardigrade species. We successfully reared two other species in the lab: *Acutuncus antarcticus,* from the Hypsibioidea superfamily, known for its resistance to high doses of UV, likely related to its exposure to high levels of UV in its natural Antarctic habitat (Giovannini et al. 2018), and *Paramacrobiotus fairbanksi* (Guidetti et al. 2019), which was reared from a garden moss and was of high interest as a representative of Macrobiotoidea, a major tardigrade superfamily considered to have diverged from Hypsibioidea more than 250 My ago (Regier et al. 2005). It was in *Paramacrobiotus areolatus*, which also belongs to Macrobiotoidea, that the first demonstration of resistance to IR was carried out (with a LD50_24h_ of 5700Gy) (May 1964). Importantly, species of Macrobiotoidea examined so far lack Dsup homologs (Arakawa 2022). We found by visual inspection of animals after IR that *A. antarcticus* and *P. fairbanksi* readily survived exposure to 1000 Gy. As done above in *H. exemplaris*, we therefore examined genes differentially expressed 4hrs after 1000 Gy IR. In both species, we found numerous genes to be significantly overexpressed in response to IR, and similar to what we observed in *H. exemplaris*, upregulation was often remarkably strong (Figure 4a and 4b, Table 2, Supp Tables 7-9, Supp Figure 8). Crucially, we identified TDR1 homologs in transcriptomes of *A. antarcticus* and *P. fairbanksi* and just like in *H. exemplaris*, these TDR1 homologs were among the most overexpressed genes in both species after IR and in response to Bleomycin treatment of *A. antarcticus* (Table 2, Supp Table 7-9), strongly suggesting a conserved role of TDR1 in resistance to IR. In contrast, as expected from previous studies, we could identify a Dsup homolog in *A. antarcticus* (Aant_geneID_rb_14333, Supp Table 7-8), from the Hypsibioidea superfamily, but not in *P. fairbanksi* from Macrobiotoidea.

**Figure 4.**
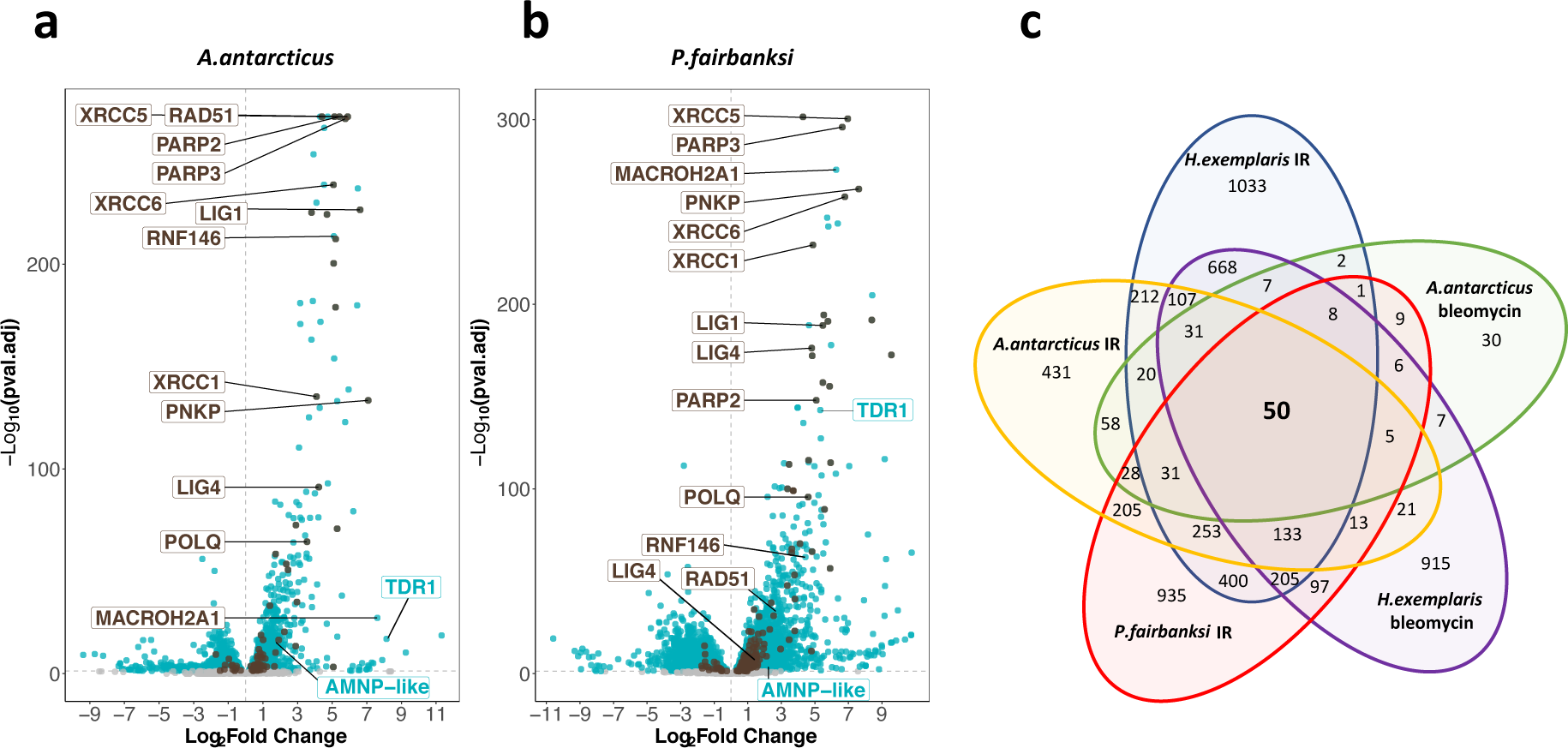
Transcriptomic response of *Acutuncus antarcticus* and *Paramacrobiotus fairbanksi* to IR. **(a)** and **(b)** Volcano plots representing Log_2_Fold Change and adjusted p-value (−log base 10) of RNA levels after irradiation with 1000 Gy γ-rays between irradiated *A. antarcticus* and untreated controls *(n=3)***(a)** and between irradiated *P. fairbanksi* and untreated controls *(n=3)* **(b).** Blue dots represent transcripts with an adjusted p-value p<0.05. Brown dots indicate transcripts of DNA repair genes (based on KEGG DNA repair and recombination gene group ko03400) with an adjusted p-value p<0.05. Brown labels indicate representative strongly upregulated genes of DNA repair. Blue labels indicate two tardigrade specific genes induced in response to IR: the TDR1 gene identified in this work, and the AMNP-like gene (BV898_10264), a member of the family of AMNP/g12777-like genes upregulated in response to desiccation and UVC (Yoshida et al 2020). **(c)** Venn diagram showing upregulated genes with an adjusted p-value p<0.05 common to the transcriptomic response to IR in the three species analysed and to Bleomycin in *H. exemplaris* and *A. antarcticus*.

**Table 2:** Number of differentially expressed genes (DEG with adjusted p-value < 0.05) after IR with 1000 Gy γ-rays *vs* untreated in 3 species (*H. exemplaris, A. antarcticus, P. fairbanksi*) and Bleomycin treatment for 4 or 5 days in *H. exemplaris* and *A. antarcticus.* A heatmap of the 50 putative orthologous upregulated genes common to all conditions is given in Supp Figure 10.

| **γ Irradiation vs Control** | *H. exemplaris* | *A. antarcticus* | *P. fairbanksi* | **Bleomycin vs Control** | *H. exemplaris* | *A. antarcticus* |
| --- | --- | --- | --- | --- | --- | --- |
| Total number of DEG | 6209 | 3708 | 7515 | Total number of DEG | 5116 | 1458 |
| DEG Up | 3178 | 1875 | 3687 | DEG Up | 2284 | 399 |
| DEG Down | 3031 | 1833 | 3828 | DEG Down | 1113 | 1059 |

Furthermore, similar to *H. exemplaris*, Gene Ontology analysis of overexpressed genes highlighted a robust enrichment of DNA repair genes in *A. antarcticus* and *P. fairbanksi* in response to IR (Supp Figure 9a-b). Notably, a high proportion of genes of the main repair pathways of DNA damages caused by IR (DSB and SSB repair, and Base Excision repair) were significantly overexpressed after IR in all three species (Supp Figure 9c-d) and as in *H. exemplaris*, among the genes with the strongest overexpression in *A. antarcticus* and *P. fairbanksi,* we observed the canonical DNA repair genes for XRCC5, XRCC6, XRCC1, PARP2, PARP3 as well as the gene for RNF146. Interestingly, a set of 50 putative orthologous genes was upregulated in response to IR in all three species, suggesting a conserved signaling and transcriptional program is involved in the response to IR between the distantly related Hypsibioidea and Macrobiotoidea superfamilies (Supp Figure 10).

### *He*-TDR1 interacts directly with DNA *in vitro* and co-localizes with DNA in transgenic tardigrades

In addition to the three species studied, BLAST searches against recent tardigrade transcriptomes enabled the identification of potential TDR1 homologs in other tardigrade species, which all belong to the Macrobiotoidea superfamily (Figure 5a, Supp Table 4). The TDR1 proteins are predicted to be 146 to 291 amino acids long, with the C-terminal part showing the highest similarity (Figure 5a). Interestingly, TDR1 proteins contain a relatively high proportion of basic amino acid residues (20.5% of K or R amino acids for TDR1 of *H. exemplaris, He-*TDR1), including at conserved positions in the C-terminal domain (Figure 5a). This led us to wonder if TDR1 might interact directly with DNA. To investigate this possibility, we purified recombinant *He*-TDR1 (Supp Figure 11) and tested its interaction with DNA using gel shift assays. As shown in Figure 5b-c, when circular or linear plasmid DNA was incubated with increasing concentrations of *He*-TDR1, a shift in plasmid mobility was detected in agarose gel electrophoresis, indicating the formation of a complex between *He*-TDR1 and DNA. The observed binding of *He*-TDR1 at a ratio of 1 He-TDR1 protein to every 3 bp of DNA is similar to the binding reported for non-sequence specific DNA binding proteins such as the Rad51 recombinase (Zaitseva, Zaitsev, and Kowalczykowski 1999). Upon adding the highest amounts of *He*-TDR1, we noted that the amount of plasmid DNA detected by ethidium bromide staining appeared to decrease. We ruled out that plasmid DNA was degraded during incubation by performing Proteinase K treatment which revealed that the amounts of intact plasmid DNA had not changed after incubation with *He*-TDR1. As an alternative explanation, we considered that at high *He*-TDR1 concentrations, *He*-TDR1 and DNA might form aggregates that could not enter the gel. To explore this possibility, we examined mixes of *He*-TDR1-GFP and plasmid DNA by fluorescence microscopy. At ratios at which complex formation was detected by agarose gel electrophoresis (Figure 5b and 5c), we observed fluorescent spots in the samples, suggesting the presence of large protein-DNA aggregates (of 2-5 µm) likely unable to enter the agarose gels (Supp Figure 12).

**Figure 5.**
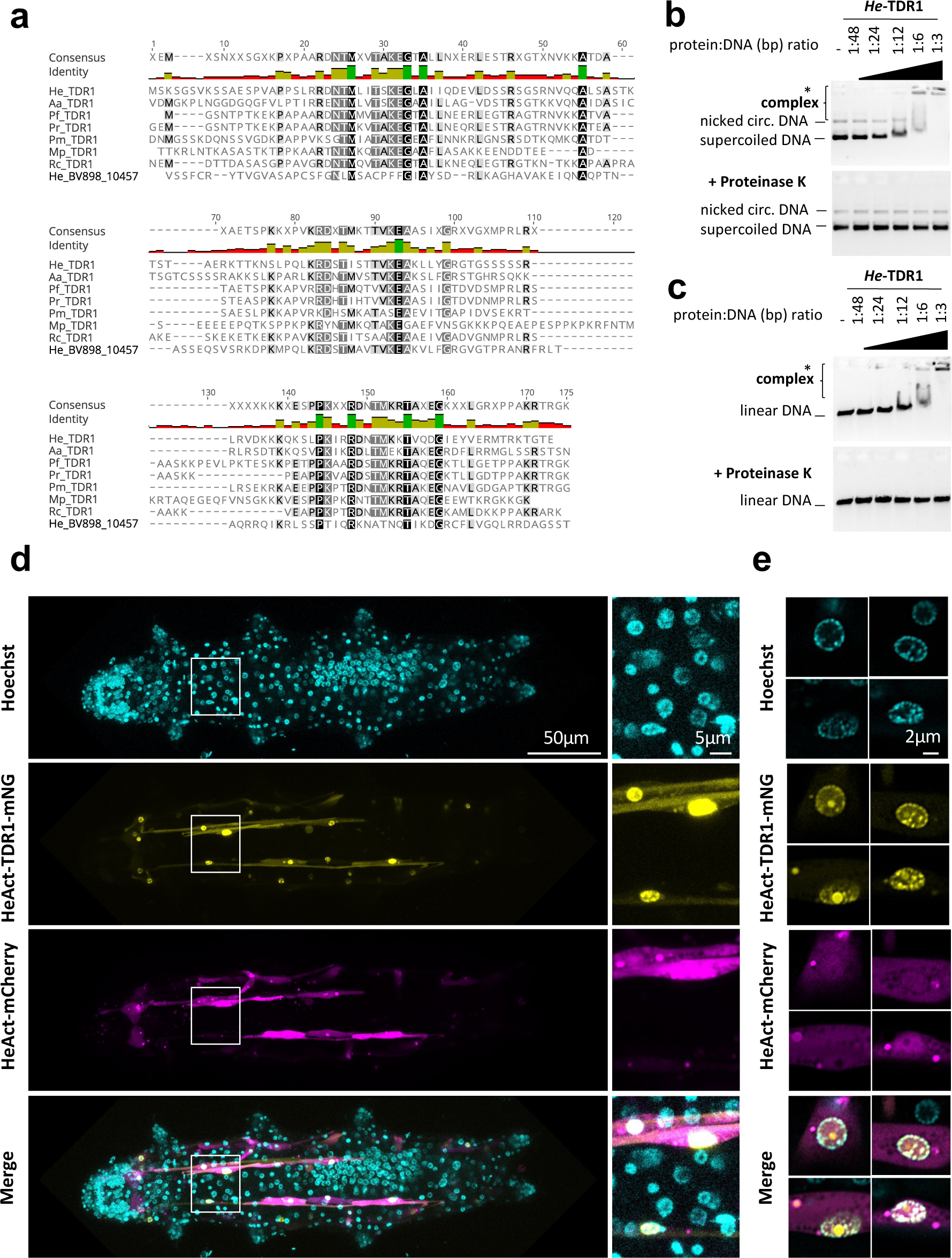
*He-*TDR1 interacts directly with DNA. **(a)** Sequence alignment of the conserved C-terminal domain of TDR1 proteins from *H. exemplaris* (He), *A. antarcticus* (Aa), *P. fairbanksi (*Pf) (identified in this work), and from *P. richtersi* (Pr) (NCBI transcriptome assembly GFGY00000000.1, now known as *P.spatialis*), *P. metropolitanus* (Pm), *R.coronifer* (Rc) (Kamilari et al 2019), *M. philippinicus* (Mp) (Mapalo et al 2020). He_BV898_10457 corresponds to a paralog of *He*TDR1 in *H. exemplaris* with weaker sequence identity to *He*TDR1 than TDR1 homologs from other species. **(b-c)** Gel shift assay of recombinant *He*-TDR1 with circular plasmid (b) or linear plasmid (c). Mixes of plasmid DNA and recombinant *He*-TDR1 at indicated protein to DNA (bp) ratios were incubated at 25°C for 20 minutes and migrated, either directly or after proteinase K digestion, at room temperature on 0.75% agarose with ethidium bromide. Fluorescence was revealed with a ChemiDoc MP imager. Complexes of plasmid DNA and recombinant *He*-TDR1 are indicated by a bracket. High-molecular weight complexes that remained in the loading wells and did not migrate into the gel are indicated by an asterisk. **(d)** Expression of *He*-TDR1-mNeonGreen in transient transgenic *H. exemplaris* tardigrades Expression plasmids of *He*-TDR1-mNeonGreen (mNG) and mCherry (both under control of the *He-*Actin promoter) were microinjected into the body fluid of *H. exemplaris* adults and electroporation was performed to induce delivery into cells following the protocol of Tanaka et al. 2023. Confocal microscopy was carried out on live animals immobilized in carbonated water at day 8 post-microinjection after 2 days of treatment with 20µM Hoechst 33342 to stain nuclei. Maximum projections of confocal Z-stack are shown. **(e)** High resolution imaging of nuclei expressing He-TDR1-mNG and Hoechst staining of the nucleus using the Airyscan2 module (one Z-slice is shown). Nuclear He-TDR1-mNG is co-localized with Hoechst staining except for 1 big foci which was observed in some high-resolution images (e-yellow channel), likely corresponding to nucleolar accumulation of overexpressed He-TDR1-mNG.

To further examine the potential interaction of *He*-TDR1 with DNA *in vivo*, we generated a tardigrade expression plasmid with *He*-TDR1-mNeonGreen cDNA downstream of *He-Actin* promoter sequences and introduced it into tardigrade cells using a recently reported protocol (S. Tanaka, Aoki, and Arakawa 2023). *He*-TDR1-mNeonGreen was easily detected in muscle cells, likely due to high muscle-specific activity of the *He-Actin* promoter, and predominantly localized to nuclei, as observed by confocal microscopy (Figure 5d-e). Importantly, *He*-TDR1 co-localized with Hoechst staining, suggesting *He-*TDR1 is able to interact with DNA *in vivo*. In summary, these experiments clearly documented interaction of *He-*TDR1 with DNA but also revealed its unexpected ability to compact DNA into aggregates.

### Expression of TDR1 proteins diminishes the number of phospho-H2AX foci in human U2OS cells treated with Bleomycin

Next, we aimed to investigate whether the expression of TDR1 could impact the number of phospho-H2AX foci detected upon treatment of human U2OS cells with the radiomimetic drug Bleomycin. When DSBs occur, H2AX is phosphorylated along extended DNA regions near the break and phospho-H2AX foci can be easily detected by immunolabeling, providing a means to indirectly visualize and quantify DSBs in nuclei (Lowndes and Toh 2005). We designed plasmids for expression of TDR1 proteins from different tardigrade species fused to GFP and transfected them into human U2OS cells. After 48h, we treated cells with 10µg/ml Bleomycin to induce DSBs. This allowed us to quantify phospho-H2AX foci in response to Bleomycin by immunolabeling with anti-human phospho-H2AX antibody. As controls, we transfected plasmids expressing either GFP, *Rv*Dsup-GFP or *He*RNF146-GFP. The quantification of phospho-H2AX was carried out in transfected cells (Figure 6a and Supp Figure 13a). As previously demonstrated for *Rv*Dsup (Hashimoto et al. 2016) and as expected from the characterization of human RNF146 (Kang et al. 2011), expression of *Rv*Dsup-GFP and *He*RNF146-GFP respectively reduced the number of phospho-H2AX foci. This result strongly suggests that *He*RNF146 is a homolog of human RNF146. Moreover, expression of TDR1-GFP fusion proteins from all species tested also significantly reduced the number of phospho-H2AX foci in human cells treated with Bleomycin, supporting the potential role of TDR1 proteins in tardigrade resistance to IR. Figure 6b shows that He-TDR1-GFP protein was localized in the nucleus of transfected cells, which is consistent with its ability to directly interact with DNA and its nuclear localization after transgenic expression in *H. exemplaris*.

**Figure 6.**
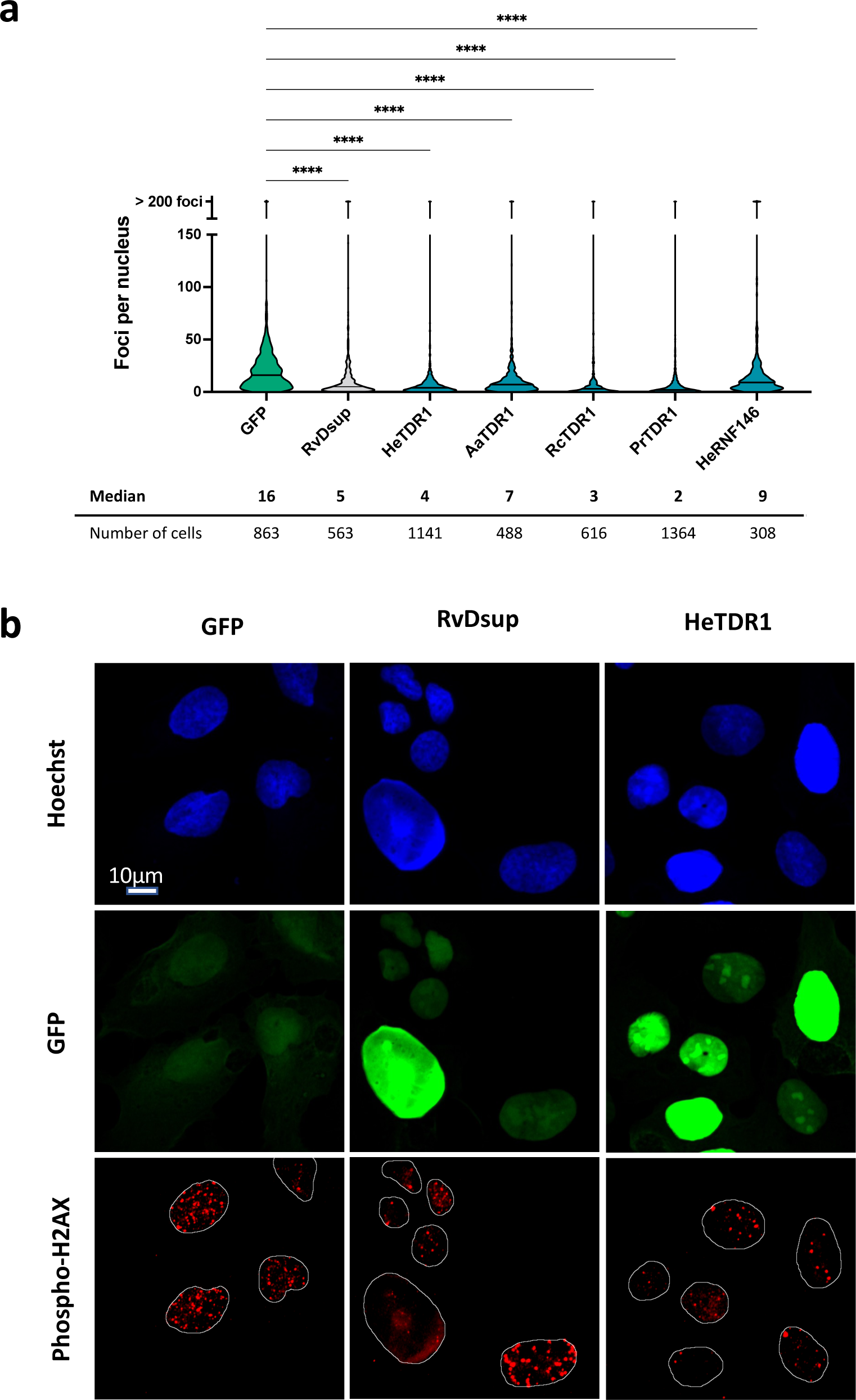
Reduced numbers of phospho-H2AX foci after Bleomycin treatment in human U20S cells expressing TDR1-GFP from multiple tardigrade species. **(a)** Violin plot of the number of phospho-H2AX foci per nucleus of cells expressing the indicated protein. Phospho-H2AX foci were counted after 1µg.ml^-1^ Bleomycin 1 hour-treatment of U2OS cells electroporated with a plasmid expressing either eGFP (control), *Rv*Dsup-GFP, TDR1-GFP from *H. exemplaris* (He), *A. Antarcticus* (Aa), *R. coronifer* (Rc) and *P. richtersi* (Pr), *He*-RNF146-GFP. Cells were fixed with a 4% PFA PBS solution for 1h, immunolabeled with chicken anti-GFP and mouse anti-phospho-H2AX antibodies and imaged by confocal microscopy. **** indicates P<0.0001 (Kruskal-Wallis test). A minimum of 308 nuclei were counted in each experimental condition (n=3). A representative experiment is shown here. Data from independent replicates are given in Supp Figure 13. **(b)** Representative confocal fluorescence imaging of experiment analyzed in (a). Images were taken with identical settings and printed with same thresholding so that signal intensity could be compared. Scale bar corresponds to 10 µM.

## DISCUSSION

Our study aimed to understand the role of DNA repair in the remarkable radio-resistance of tardigrades. We examined the DNA damage and repair mechanisms in the tardigrade species *H. exemplaris* after exposure to ionizing radiation (IR) and performed comparative transcriptomics in three species of the Tardigrada phylum. Our results indicate that DNA repair plays a major role in the radio-resistance of tardigrades compared to human cells and identified the gene for TDR1, a novel DNA-binding protein highly upregulated in response to IR and likely to play an original function in DNA repair.

### DNA repair plays a major role in resistance of tardigrades to IR

Using an antibody raised against phosphorylated *He*-H2AX, we could detect DSBs by Western blot and by immunolabeling (Figure 1). Our analysis documented dose-dependent DNA damage and repair taking place after exposure to IR. DNA damage could be detected in virtually all nuclei by immunolabeling. However, at 1000 Gy, phosho-H2AX labeling persisted longer than at 100 Gy in the gonad. Additionally, at 1000 Gy, cell divisions could no longer be detected in the midgut of the digestive system. These two consequences of exposure to higher doses of IR may be due to higher sensitivity of replicating cells to IR and explain why *H. exemplaris* tardigrades no longer lay eggs and become sterile after irradiation with 1000 Gy (Beltrán-Pardo et al. 2015).

Using standard agarose gel electrophoresis, we were able to observe that SSBs were induced every 10 to 20 kb in *H. exemplaris* after exposure to 1000 Gy of γ-rays, indicating a rate of 0.05 to 0.1 SSB/Gy/Mb (Figure 1c). Remarkably, this rate is roughly similar to that reported for cultured human cells which is 0.17 SSB/Gy/Mb (Mohsin Ali et al. 2004), showing that high levels of DNA damage are induced after high doses of IR and thus supporting the importance of DNA repair in the radio-resistance of *H. exemplaris* compared to human cells. In radio-resistant rotifers, the rate of DSBs was comparable to non-radioresistant organisms (Gladyshev and Meselson 2008), also suggesting the importance of DNA repair in radio-resistance. Concerning the role of DNA protection in radio-resistance, further studies would be necessary; in particular, determining the rate of DNA double-strand breaks and testing the importance of Dsup in live *H. exemplaris*. Quantification of phospho-H2AX foci is frequently used as a proxy but given the small size of tardigrade nuclei, standard imaging by confocal microscopy was not sufficient to clearly identify and quantify independent phospho-H2AX foci. Recent developments in super-resolution microscopy could make it possible to perform such quantification in the future (Tillberg and Chen 2019). Pulse field electrophoresis is a method that would allow to directly examine DNA damage, but it would require to disrupt the cuticle and release DNA without causing damage which would confound the analysis of DNA integrity.

### Fine regulation of scaffolding proteins to cope with high rates of DNA damage

We performed comparative transcriptomics in three different species and uncovered the conserved upregulation of a wide number of DNA repair genes in response to IR (Figure 4 and Supp Figure 9). Remarkably, the strongest upregulations, both at RNA and protein levels, were detected for proteins acting early in DNA repair in the different pathways involved: XRCC5/XRCC6 in NHEJ, POLQ in MMEJ, XRCC1 in SSB, and PARP2/PARP3 which act as DNA damage sensors common to all double-strand break repair pathways (Pandey and Black 2021). These early acting proteins either stabilize DNA ends or provide essential scaffolding for subsequent steps of DNA repair. It is possible that producing higher amounts of such proteins is essential to maintain DNA ends long enough for more limiting components of DNA repair to cope with an exceptionally high number of damages. For XRCC5 and XRCC6, our study established, by two independent methods, proteomics and Western blot analyses, that the stimulation at the protein level was much more modest (6 and 20-fold at most, Supp Figure 6) than at the RNA level (420 and 90 fold respectively). This finding suggests that the abundance of DNA repair proteins does not simply increase massively to quantitatively match high numbers of DNA damages. Interestingly, in response to IR, the RNF146 ubiquitin ligase was also found to be strongly upregulated. RNF146 was previously shown to interact with PARylated XRCC5 and XRCC6 proteins and to target them for degradation by the ubiquitin-proteasome system (Kang et al. 2011). To explain the more modest fold stimulation of XRCC5 and XRCC6 at protein levels compared to its massive increase at mRNA levels, it is therefore tempting to speculate that, XRCC5 and XRCC6 protein levels (and perhaps that of other scaffolding complexes of DNA repair) are regulated by a dynamic balance of synthesis, promoted by increased gene transcription, and degradation, made possible by RNF146 upregulation. Consistent with this hypothesis, we found that, similar to human RNF146 (Kang et al. 2011), *He*-RNF146 expression in human cells reduced the number of phospho-H2AX foci detected in response to Bleomycin (Figure 6).

Further studies should investigate the molecular mechanisms leading to such marked upregulation of RNA levels of these genes. Functional analysis of promoter sequences in transgenic tardigrades is now possible (S. Tanaka, Aoki, and Arakawa 2023) and could help to identify a conserved set of transcription factors and/or co-regulators common to Macrobiotodea and Hypsibioidea tardigrades. Such information would provide original insight into the acquisition of resistance to IR and help analyze its relation to the resistance to desiccation. Another outstanding issue, given the high rates of DNA damage taking place, is whether DNA repair is accurate. This is particularly relevant for germ line cells where mutations will be transmitted to the progeny and could impact evolution of the species.

### A novel tardigrade specific DNA binding protein involved in resistance to IR

Among the genes overexpressed in response to IR in the three species studied, we identified TDR1 as a promising tardigrade specific candidate (Figures 3 and 4). At the functional level, we found that TDR1 protein interacts with DNA (Figure 5) and that when expressed in human cells, TDR1 protein can reduce the number of phospho-H2AX foci induced by Bleomycin (Figure 6). TDR1 is strongly overexpressed in response to IR (Figure 3), suggesting that it favors DNA repair. Proteins directly involved in DNA repair, however, usually accumulate at sites of DNA damage (Rothkamm et al. 2015), which did not appear to be the case for TDR1 overexpressed in human cells. Given that TDR1 can form aggregates with DNA *in vitro*, we speculate that it may favor DNA repair by regulating chromosomal organization. Intriguingly, the DNA binding activities of TDR1 are reminiscent of DdrC from *D. radiodurans*. DrdC is a small DNA binding protein which is among the most strongly overexpressed proteins after irradiation of *D. radiodurans* with γ-rays (M. Tanaka et al. 2004) and DdrC forms aggregates with DNA *in vitro* (Banneville et al. 2022; Bouthier de la Tour et al. 2017). Further investigations of TDR1 may thus reveal unexpected parallels between mechanisms of DNA repair conferring radio-resistance in tardigrades and bacteria.

Recent progress in tardigrade transgenesis (S. Tanaka, Aoki, and Arakawa 2023) and promising findings of somatic mutagenesis by CRISPR-Cas9 (Kumagai, Kondo, and Kunieda 2022) are paving the way towards germ line gene editing in *H. exemplaris*. Knocking out Dsup and TDR1 genes should help to better appreciate their importance in radio-resistance and the underlying mechanisms. The C-terminal portion of TDR1 is conserved in species of Macrobiotoidea and Hypsibioidea superfamilies of the Parachela order of Eutardigrada but absent from *Milnesium inceptum*, the only representative of the Apochela order of Eutardigrada for which transcriptomic data is currently available (Supp Figure 14), and from Heterotardigrada. Compared to Dsup, which has only been found in Hypsibioidea, TDR1 appears more widely present and could be a more ancient tardigrade gene. As additional tardigrade species, more fully representing the phylogenetic diversity of the phylum, are reared in laboratory conditions and become amenable to experimental analysis, more novel genes and mechanisms of radio-resistance may become apparent. Generally, evolution of tardigrade specific gene sequences appears highly dynamic in the phylum (Arakawa 2022), and further sequencing of tardigrade genomes will help get a better picture of gain and loss of tardigrade specific genes and their relation to resistance to extreme conditions.

In conclusion, our findings suggest that DNA repair is a major contributor to tardigrade radio-resistance. Functional investigations of TDR1, as well as the study of transcriptional regulation in response to IR, will contribute to a deeper understanding of the mechanisms underlying radio-resistance. Additionally, we believe that, as done here, further exploration of tardigrade-specific genes and comparative studies among tardigrade species will shed light on the evolution and diversity of radio-resistance mechanisms in these fascinating organisms.

## MATERIAL AND METHODS

### Tardigrade culture

*Hypsibius exemplaris* (strain Z151, Sciento, UK) and *Acutuncus antarcticus* (Giovannini et al. 2018) were cultivated from 1 individual in mineral water (Volvic, France) at 16°C with a 14h day and 10h night cycle and fed with *Chlorella vulgaris* microalgae (from Algothèque MNHN, France or ordered from ldc-algae.com (Greenbloom pure fresh Chlorella). Microalgae were grown in 1X Cyanobacteria BG-11 Freshwater Solution (C3061, Sigma-Aldrich) at 23°C until saturation, collected by centrifugation at 2000g for 5 min and resuspended in Volvic mineral water 10-fold concentrated before feeding. For concentrated large amounts of *H. exemplaris* 0.2% of linseed oil emulsion (containing 95% linseed oil (organic), 1% Tween 80 (P4780, Sigma-Aldrich) and 4% (+/-)-α-Tocopherol (T3251, Sigma-Aldrich) could be spread at the bottom of the untreated petri dishes before adding algae and water. *Paramacrobiotus fairbanksi* was isolated from a suburban garden moss by A.D.C. and cultivated in the same conditions adding rotifers isolated from the same moss sample (grown separately in Volvic mineral water and fed with *C. vulgaris*) of as supplementary food for adults. Identification as *P. fairbanksi* was achieved by morphological and DNA markers (Supp Figure 15).

### IR and Bleomycin treatments of tardigrades

Prior to treatments, tardigrades were separated from *Chlorella* by filtering with a cell strainer 70 µm mesh (141379C, Clearline) for *H. exemplaris* or 100 µm mesh (141380C, Clearline) for *A. antarcticus* and *P. fairbanksi.* The cell strainer containing the washed tardigrades was put in a Petri dish containing Volvic mineral water for 3 to 7 days in order to allow live tardigrades to go out of the cell strainer and obtain tardigrades without any remaining *Chlorella*. Tardigrades were then collected on 40 µm mesh (141378C, Clearline) and washed with Volvic mineral water before proceeding to treatments. For each treatment, tardigrades were collected and split into treated and control samples. Control samples were subjected to the same conditions (temperature, travel (for irradiation), solvent (for Bleomycin)) as the treated tardigrades. For ionizing irradiations, tardigrades were exposed to a ^137^Cs γ-ray source (GSR-D1 irradiator, RPS Services Limited) at a dose rate of 12.74 or 16 Gy/min for a total dose of 100 or 1000 Gy. Samples were collected at different time points after the end of irradiation (for differential transcriptome analyses, at 4h post-irradiation; for differential proteomics, at 24h post-irradiation; for DNA damage and Western blot analyses, at different time points from 0h to 7 days as indicated in the text). For investigation of early first time point just after ionizing radiation, an electronic beam (KINETRON, GCR-MeV) with 4.5MeV energy and maximum dose rate of 4.103 Gy/sec was also used (Lansonneur et al. 2019). For Bleomycin treatment, after separating tardigrades from *Chlorella* by filtration, Bleomycin sulfate (#B5507, SIGMA) was added to the water at a concentration of 100 µM for 4 or 5 days. The response to Bleomycin treatment was examined in *H. exemplaris* and *A.antarcticus* but not in *P. fairbanksi* tardigrades (which grow more slowly than *H. exemplaris* and *A.antarcticus*).

### Production of antibodies against *H. exemplaris* proteins

Antibodies were raised against *H. exemplaris* proteins in rabbits by injecting selected peptide sequences by Covalab (Bron, France). For *He*-Ku80 (XRCC5), *He*-Ku70 (XRCC6 C-term), *He*-Dsup, *He*-TDR1 two peptides were injected in two rabbits and serums tested by Western blot on *H. exemplaris* extracts from animals treated with 100 µM Bleomycin for 4 days. Serum showing the best response on day 88 after injections was purified on Sepharose beads coupled to immunogenic peptides. Peptides used were the following, *He*-TDR1: Peptide 1 (aa 37-52, C-IQDEVLDSSRSGSRNVcoNH2), Peptide 2 (aa 109-123, C-DKKKQKSLPKIRRDN-coNH2); *He*-Ku80 (XRCC5): Peptide 1 (aa 120-135, C-IQFDEESSKKKRFAKR-coNH2), Peptide 2 (aa 444-457, C-LDGKAKDTYQPNDE-coNH2); *He*-Ku70 (XRCC6 Cterm): Peptide 1 (aa 182-197, C-IRPAQFLYPNEGDIRG-coNH2), Peptide 2 (aa 365-37, C-YDPEGAHTKKRVYEK-coNH2); *He*-Dsup: Peptide 1 (aa 63-77, C-KTAEVKEKSKSPAKE-coNH2), Peptide 2 (aa 166-181, C-KEDASATGTNGDDKKE-coNH2). Production of antibody to *He*-phospho-H2AX is detailed in Supp Figure 1.

### Western blot analysis

For each experiment, more than 10 000 *H. exemplaris* tardigrades were irradiated or untreated, and 1000-2000 tardigrades were collected at different time points after irradiation (30min, 4h, 8h30, 24h and 73h), centrifuged at 8000 rpm in 1.5 mL tubes for 5 min and the pellet frozen at -80°C until analysis. Lysis was carried out by sonication for 15 min (15s ON/15s OFF, medium intensity, Bioruptor, Diagenode) at 4°C in 100 µL/5000 tardigrades pellet of the following solution 12mM sodium deoxycholate, 12mM N-Lauryl sarcosine sodium salt, 80mM Tris-HCl pH8,5, 400mM NaCl, 1X cOmplete protease inhibitor (4693116001, Roche) and 1X PhosSTOP (PHOSS-RO, Roche). 0.4 vol of protein gel loading buffer (LDS 4X, Bio-Rad) and 0.1 vol of DTT 1M was added and the mixture heated at 95°C for 5 min before loading onto Any kD Mini-PROTEAN TGX Stain-Free protein gel (4568124, Bio-Rad) and migration in 1x Tris-Glycine-SDS buffer at 200V. Semi-dry transfer of proteins was performed with Transblot Turbo (Bio-Rad) onto nitrocellulose membrane and the membrane was cut half to separately detect proteins > 50 kDa and <50 kDa. Protein detection was done with rabbit primary antibodies diluted 1:1000 or 1:2000 (20-200ng/mL depending on antibody) in TBS-0.1% Tween 20, 5 % BSA, supplemented with 1:10000 dilution of anti-mouse alpha-Tubulin (Clone B-5-1-2; SIGMA) for 1 to 3 h at room temperature. Membranes were washed 3 times for 5 min with 1X TBS 0.1% Tween 20. Secondary antibodies diluted 1:5000 (anti-rabbit Starbright 700 (12004161, Bio-Rad) for specific tardigrade proteins and anti-mouse Starbright 520 (12005867, Bio-Rad) for alpha-tubulin detection) in TBS-1% milk were incubated on membrane for 1h at room temperature. Membranes were washed 3 times for 5 min with 1X TBS 0.1% Tween 20 and unsaturated fluorescent signal was acquired using Chemidoc MP imager (Bio-Rad) with identical settings for samples to be compared within an experiment.

### Genomic DNA extraction and analysis

For each experiment, more than 60 000 *H. exemplaris* tardigrades were irradiated or untreated, and 8000-12000 tardigrades were collected at different times after irradiation (30min, 4h, 8h30, 24h and 73h), centrifuged at 8000 rpm in 1.5 mL tubes for 5 min and the pellet frozen at -80°C until analysis. Genomic DNAs (gDNAs) were extracted using the Monarch HMW DNA extraction kit for Tissue from New England Biolabs (NEB) with the following modifications: Lysis buffer was supplemented with proteinase K before proceeding to lysis. Pellets were resuspended in 35 µL of lysis buffer and grinded on ice for 1 minute. This step was repeating twice leading to a final volume of ≈ 125 µL. After grinding, lysis proceeded in three steps: 1) incubation of 15 min at 56°C under gentle agitation (300 rpm), 2) incubation of 30 min at 56°C and 3) incubation of 10 min 56°C after addition of RNAse A. Proteinase K and RNAse A were added at the concentration recommended by NEB. Proteins were next separated from the gDNA by adding 40 µL of protein separation solution. Samples were next centrifuged (20 minutes, 16000 g, 20°C). gDNA was precipitated with 2 beads and next eluted from the beads with 100 µL of elution buffer. Extracted gDNAs were analysed by electrophoresis on native (0.9% agarose/1X TAE) or denaturing (0.9% agarose/ 30 mM NaOH/ 1 mM EDTA) gels. Electrophoresis conditions were: 2h30min/ 60 V/ 20°C for native gels, and 15 h/18 V/ 20°C for denaturing gel. Native gels were stained with ethidium bromide and denaturing gels with SyBR Green I.

### Immunohistochemistry of tardigrades

Immunohistochemistry protocol was derived from (Vladimir Gross and Mayer 2019; V. Gross, Bährle, and Mayer 2018). 10 000 tardigrades irradiated or untreated were sampled (by batches of 1000 tardigrades) at different time points after irradiation (5min, 4h, 24 or 72h), heated in Volvic mineral water 5 min at 70°C to extend the tardigrade body and directly fixed with 4% formalin (15686, EMS) in 1X PBS-1% Triton-X100 (PBS-Tx) by adding 5X solution. Fixation was carried out for 1 to 3 h at room temperature. After 1h of fixation tardigrade were pellet by centrifugation 5min at 8000rpm and kept in 200µL of fixative solution. The cuticule was punctured by sonication using Bioruptor (Diagenode) in 6x1.5mL tubes at a time (5 pulses of 5s ON/ 5s OFF in medium position). After fixation samples were pelleted by centrifugation and washed with 1mL of 1X PBS-1% Triton-X100 3 times (>3h wash) and tardigrades were transferred to a round well 96 plate for transferring tardigrades under the stereomicroscope. Blocking was done in 200µL of 5% BSA in 1X PBS-1% Triton-X100 for at least 1h and in-house primary antibody against phosho-H2AX (rabbit) (dilution 1:10) in blocking buffer was applied on tardigrades overnight or for 3 days at 4°C. Washes with PBS-Tx were done 4 times for several hours the next day transferring tardigrades to a new well filled with 200µL of PBS-Tx. Secondary antibody anti-rabbit Alexa Fluor 488 F(ab’)2 (A11070, Invitrogen, dilution 1:500) in PBS-Tx supplemented with 1% BSA was incubated overnight at room temperature. Washes with PBS-Tx were done 4 times for several hours the next day transferring tardigrades to a new well filled with 200µL. The last wash was done without Triton X100. Hoechst 33342 4µM in PBS 1X was incubated for 30min and tardigrades were quickly transferred to water and to slide with minimum amount of liquid and finally mounted with ProLong™ Glass Antifade Mountant (P36982, Invitrogen).

For analysis of EdU staining, EdU in 20mM Hepes-NaOH pH 7.5 was added to Volvic mineral water with 1000 filtered tardigrades at 50µM 2h before irradiation or with control, untreated tardigrades and kept until 7 days post irradiation. Samples were then processed as in (V. Gross, Bährle, and Mayer 2018) except for the permeabilization of tardigrades which was carried out by sonication as for phospho-H2AX labelling.

Imaging was done by confocal microscopy (Zeiss (LSM 880 and AiryScan module) with x63 lens) using Zenblack software or Leica DMIRE2 inverted microscope with 10X lens and Metamorph software. Confocal Z-stacks and Maxprojections were processed and adjusted in Fiji ImageJ (v2.9.0). Images were treated with Image J software. Image panels were assembled and labelled in Microsoft powerpoint for Mac (v16.66.1)

### RNA sequencing

15000-20000 *H. exemplaris* (n=3), 1000-1500 *A. antarcticus* (n=3) and 500-1000 *P. fairbanksi* (n=4) for each independent biological sample were collected and subjected to IR treatments: (i) control animals non irradiated and extracted 4h post-irradiation and (ii) irradiated animals (with ^137^Cs γ-ray source (GSR-D1 irradiator, RPS Services Limited) at a dose rate of 16Gy/min) extracted 4h post IR. 15000 -20000 *H. exemplaris* and 1000-1500 *A. antarcticus* (3 independent biological samples for each) were also subjected to Bleomycin treatment: (iii) control animals kept for 5 days in Volvic water and (iv) treated with 100µM Bleomycin in Volvic mineral water for 5 days. After treatments, tardigrades were collected and washed by filtration on 40µm nylon mesh and transferred to 1.5mL tubes to pellet by centrifugation at 5 min at 8000 rpm. RNA was extracted using Trizol (15596-026, Invitrogen) and by three freeze-thaw cycles in an ethanol-dry ice bath and mechanical disruption with glass beads and a plastic micro-tube homogenizer at each cycle. Yield was approximately 1µg RNA/1000 tardigrades. Integrity of RNAs was checked on an Agilent 2100 Bioanalyzer with the Eukaryote Total RNA Nano kit and only samples with RNA Integrity Number >6 were sequenced. For *H. exemplaris* RNA samples, single-end (1x75) sequencing (TruSeq Stranded) was done on Illumina NextSeq 500 System. For *A. antarcticus* and *P. fairbanksi* (whose genomes are not available), paired-end (2x150) sequencing (TruSeq Stranded) was performed. In addition to short read RNA sequencing in the different experimental conditions, long read sequencing of a mixture of RNA samples of *A. antarcticus* and of RNA samples of *P. fairbanksi* species were performed with Oxford Nanopore Technology (ONT) to help improve transcriptome assembly. 1D libraries were prepared according to ONT protocol with 1D PCR Barcoding kit and full length non-directional sequencing was performed on PromethION instrument (using Clontech- SMART-Seq v4 Ultra Low Input kit). Basecalling was conducted using guppy version (v6.4.2; parameters: --min_qscore 7 --flowcell FLO-MIN106 --kit SQK-PBK004 --use_quantile_scaling --trim_adapters --detect_primer --trim_primers).

### *De novo* transcriptome assembly

*De novo* transcriptome assembly was performed using full length cDNA sequences for *A. antarcticus* and *P. fairbanksi.* We used RNA-Bloom (v2.0.0; Ka Ming Nipal. 2023), to assemble the long reads, also using a subset of the produced short reads to correct the contigs. Then we used MMSeqs2 easy-cluster (v; parameters: --min-seq-id 0.85 -c 0.25 --cov-mode 1) to cluster together transcript isoforms to a gene (Mirdita, Steinegger, and Söding 2019). We set the minimum sequence identity to 0.85 and minimum coverage to 0.25 for both transcriptomes. Because *P. fairbanksi* is triploid with a high level of heterozygosity, we manually clustered differentially expressed genes that were annotated for the same function by EggNOG (see below). We aligned the isoforms from two or more clusters with the same EggNOG annotation using mafft (version 1.5.0; Katoh and Standley 2013) and we visually inspected the alignments on Geneious Prime (v2023.1). When isoforms from two or more clusters were properly aligning together, they were merged. For *A. antarcticus and P. fairbanksi*, we conducted the gene expression analysis using the softwares embedded in the Trinity suite (v2.15.0; Haas et al. 2013). We first mapped RNA-seq reads on the transcriptomes using Salmon (statistics of the mapping of RNAseq reads are given in Supp Table 11; Patro et al. 2017), then we measured differential gene expression using DESeq2 (Love, Huber, and Anders 2014). For *H. exemplaris,* as the genome was available, the gene expression analysis was conducted using Eoulsan workflow version 2.2 (v2.2-0-g4d7359e, build0 build on 764feac4fbd6, 2018-04-17 15:03:09 UTC) (Lehmann et al. 2021). We first mapped RNA-seq reads on the *de novo* transcriptomes using STAR (Dobin et al. 2013), then we measured differential gene expression using DESeq2. The results were plotted using R (v4.2.2) with the ggplot2, ggrepel, and VennDiagram packages. Heatmap was plotted using GraphPad Prism (v9.3.1).

To annotate expressed genes from the three species, we ran EggNOG mapper (v2.1.9) on the assemblies using the “genome” mode (Cantalapiedra et al. 2021). We also annotated all expressed genes through a sequence homology search against *Drosophila melanogaster* (GCF_000001215.4), *Caenorhabditis elegans* (GCF_000002985.6) *Homo sapiens* (GCF_000001405.40), *Hypsibisus exemplaris* (GCA_002082055.1), *Paramacrobiotus metropolitanus* (GCF_019649055.1) and *Ramazzottius varieornatus* (GCA_001949185.1). Since *H. exemplaris* genome is annotated, we ran the homology search against the target proteomes using blastp (v. 2.14.0). For *A. antarcticus* and *P. fairbanksi,* we conducted the homology search using the transcript as query (blastx) and as target (tblastn). Only blast hits with an e-value <0.05 were kept as potential homologs.

To identify tardigrade-specific genes, we ran a homology search using Diamond (v2.1.6.160) (Buchfink, Reuter, and Drost 2021) on the complete nr database (Downloaded Apr 12 11:17:28 2023) for each transcript from *A. antarcticus* and *P. fairbanksi* (by blastx) or for each protein sequence for *H. exemplaris* (by blastp). Sequences with no-hit on the nr database (diamond blastx or blastp –e 0.001 --taxon-exclude 42241 --ultra-sensitive) and no hit in the previous annotation using proteomes of C. elegans, D. melanogaster and H. sapiens (reciprocal hit – blastx and tblastn) were considered as tardigrade specific and noted “TardiSpe” in Supp Tables and data (Mapalo et al. 2020; Hara et al. 2021; Kamilari et al. 2019). Additionnally, for TDR1 we conducted a blast search on the nr database using NCBI blastp, which is more sensitive but slower, than diamond blastp. In contrary to diamond, NCBI blastp produced multiple hits but on non-ecdysozoans organisms only (see Supp Table 5). Similar results were obtained using HMMER (hmmsearch on EMBL-EBI website) on the reference proteome database (see Supp Table 5).

### Proteome analysis

For each replicate (n=4 independent biological samples), 18000 tardigrades for each of the four experimental conditions: i) untreated; ii) treated with Bleomycin at 100µM for 4 days; iii) Irradiated (with ^137^Cs γ-ray source (GSR-D1 irradiator, RPS Services Limited) at a dose rate of 12.74Gy/min) and collected after 4h; iv) irradiated and collected after 24h. The tardigrades were split in two samples, with 13 000 tardigrades for differential proteomic analysis and 5000 tardigrades for western blotting experiments, that were pelleted by centrifugation in 1.5mL tubes (8000 rpm for 5 min). The pellets were frozen at -80°C until all samples were available. All samples were lysed the same day 2 weeks before proteomics analysis in 100µL iST-NHS-Lysis buffer (PreOmics GmbH) by sonication (Bioruptor Diagenode, 15s ON/15s OFF for 15 min), and heating at 95°C for 10min. Soluble fractions were collected by centrifugation at 13000g for 15min at 4°C and frozen at -80°C until analysis. Protein concentration in each sample was measured using BCA assay (Sigma-Aldrich). 30µg of each sample were then prepared using the iST-NHS kit (Preomics). Peptides resulting from LysC/trypsin digestion were labelled using TMTpro™ 16plex Label Reagent Set (ThermoFisher Scientific) before mixing equivalent amounts for further processing. The peptide mix was then fractionated using the Pierce High pH Reversed-Phase Peptide Fractionation Kit (ThermoFisher Scientific). The 8 obtained fractions were analyzed by online nanoliquid chromatography coupled to MS/MS (Ultimate 3000 RSLCnano and Q-Exactive HF, Thermo Fisher Scientific) using a 180 min gradient. For this purpose, the peptides were sampled on a precolumn (300 μm x 5 mm PepMap C18, Thermo Scientific) and separated in a 200 cm µPAC column (PharmaFluidics). The MS and MS/MS data were acquired by Xcalibur (version 2.9, Thermo Fisher Scientific). The mass spectrometry proteomics data have been deposited to the ProteomeXchange Consortium via the PRIDE (Perez-Riverol 2022) partner repository with the dataset identifier PXD043897.

Peptides and proteins were identified and quantified using MaxQuant (version 1.6.17.0, Cox and Mann 2008) and the NCBI database (*Hypsibius dujardini* taxonomy, 2021-07-20 download, 20957 sequences), the UniProt database (*Chlorella* taxonomy, 2021-12-10 download, 21 219 sequences) and the frequently observed contaminant database embedded in MaxQuant (246 sequences). Trypsin was chosen as the enzyme and 2 missed cleavages were allowed. Peptide modifications allowed during the search were: C6H11NO (C, fixed), acetyl (Protein N-ter, variable) and oxidation (M, variable). The minimum peptide length and minimum number of unique peptides were respectively set to seven amino acids and one. Maximum false discovery rates - calculated by employing a reverse database strategy - were set to 0.01 at peptide and protein levels. Statistical analysis of MS-based quantitative proteomic data was performed using the ProStaR software (Wieczorek et al. 2017). Proteins identified in the reverse and contaminant databases, proteins identified only in the *Chlorella* database, proteins only identified by site, and proteins quantified in less than three replicates of one condition were discarded. After log2 transformation, extracted corrected reporter abundance values were normalized by Variance Stabilizing Normalization (vsn) method. Statistical testing for comparison of two conditions was conducted with limma, whereby differentially expressed proteins were sorted out using a Log_2_(Fold Change) cut-off of 0.3 and a limma p-value cut-off of 0.01, leading to a FDR inferior to 3 % according to the Benjamini-Hochberg estimator.

### Production of recombinant *He*-TDR1 and *He*-TDR1-GFP

*He*-TDR1 and *He*-TDR1-GFP (see plasmid sequence in Supp Table 4) were transformed in *E. coli* Rosetta^TM^ 2(DE3). Singles competent cells (Novagen, MerckMillipore). Protein expression was induced with 1 mM IPTG at OD_600_=0.6-0.7 in 2xYT medium (containing 50 µg/mL carbenicillin, 35 µg/mL chloramphenicol and 1% glucose) at 25°C during 20h. Cells were resuspended in lysis buffer 25 mM Tris-HCl pH 8, 500 mM NaCl, 20 mM imidazole, 1 mM TCEP (supplemented with protease inhibitor cocktail (Roche)), and lysed by sonication (Vibracell 75186 -7 sec ON / 7 sec OFF, 50% amplitude, 10 min). The first step of purification was binding on Ni Sepharose 6 Fast Flow resin in batch (overnight at 4°C). After binding, the resin was washed with lysis buffer and the protein was eluted with 25 mM Tris-HCl pH 8, 500 mM NaCl, 250 mM imidazole, 10% glycerol, 1 mM TCEP. Eluted protein is concentrated (Amicon Ultra 10K) and diluted in buffer 25 mM Tris-HCl pH8, 150 mM NaCl, 10% glycerol, 1 mM TCEP. The second step of purification was a gel filtration Superdex 200 increase 10/300 GL (Cytiva) equilibrated with 25 mM Tris-HCl pH 8, 150 mM NaCl, 10% glycerol, 1 mM TCEP using AKTA Pure instrument (Cytiva). Molecular weight calibration was obtained using Gel Filtration Standard (Bio-Rad).

### Protein-DNA interaction assays

For *He*-TDR1 interaction with plasmid DNA, a 5900 bp plasmid (a kind gift of Xie, Kwok, and Scully 2009) circular or linearized at 20 ng/µL (ie: 30 µM in bp was incubated with increasing amounts (0.625 to 10 µM with 2-fold serial dilutions) of recombinant *He*TDR1 or in buffer containing 15 mM Tris-OAc pH8, NaCl 180 mM, Glycerol 2%, DTT 5 mM, BSA 0.1 mg/mL.

After 20 min binding at room temperature, samples were diluted 2-fold with sucrose 50% (or sucrose 50% with proteinase K 80 U/µL and loaded onto a 0.75% agarose gel containing ethidium bromide. Migration was carried out for 35 min at 100 V (room temperature) and gel was imaged using GBox camera (Syngene).

For imaging of protein-DNA complexes, 1 µL of 5900pb plasmid at 200 ng/µL was added to 10 µL of 10 µM of *He*TDR1-GFP in 10 mM Tris-HCl PH8, 150 mM NaCl, 10% Glycerol and 1 mM TCEP (protein storage buffer) to allow 30 µM in bp (i.e.: 5 nM in plasmid molecule) final concentration. After 10 min incubation at room temperature the reaction was observed in a Kova counting chamber using Leica DMIRE2 40X lens. Images were acquired using Coolsnap HQ camera run by Metamorph software and treated with ImageJ software.

### Expression of *He*-TDR1-mNeongreen in *H. exemplaris* tardigrades

Act-He-TDR1-mNeongreen (NG) and Act-mCherry expression plasmids were constructed by Gibson assembly with plasmid backbone from (Loulier et al. 2014) (see sequence in Supp Table 4). Actin promoter sequences were amplified from *H. exemplaris* genomic DNA, *He*TDR1 cDNA from RNA of *H. exemplaris* adult tardigrades and mCherry from a mCherry containing plasmid. *He-*Act-*He*TDR1-GFP and Act5C-mCherry plasmids (2 µg/µL in milliQ water each) were co-injected in 20 starved *H. exemplaris* adults maintained in an in-house made PDMS injection chamber using Quartz micropipets. After 1h of microinjection, animals are let to recover in Volvic mineral water for 15min to 1h15. In order to get the plasmid into cells, tardigrades are next electroporated using NEPA21 Super Electroporator (Nepa Gene). Electric shock was carried out in 0.7X Optimem (Gibco, Thermofisher Life Sciences) with settings from (S. Tanaka, Aoki, and Arakawa 2023). Hoechst 33342 20 µM for 2 days or 40 µM for 1 day was also added to mineral water (Volvic, France) for live staining of the nucleus. Animals were immobilized using carbonated water and imaged by confocal microscopy (Zeiss (LSM 880 and AiryScan module) with x40 and x63 lens) using Zenblack software.

### Expression of TDR1-GFP fusion proteins in human U2OS cells

Expression plasmids for fusion proteins of GFP and tardigrade proteins were constructed by Gibson assembly into pEGFP-N1 (Clontech) of the tardigrade cDNA (obtained by gene synthesis from Integrated DNA) or ordered from TwistBiosciences. Full nucleotide sequences of fusion proteins are provided as supplementary information (Supp Table 4). Plasmids were transfected into human U2OS cells (ATCC HTB-96) by Amaxa electroporation with Nucleofector™ Kit V (Lonza) and plated in 6 well plates containing glass slides.

### Immunolabeling of phospho-H2AX foci in response to Bleomycin treatment and image analysis

Two days after transfection, Bleomycin sulfate-treated (treatment was for 1h with 1 µg/mL Bleomycin sulfate) or control cells were rinsed three times with PBS and fixed with 3.7% formaldehyde in PBS for 15 min at room temperature, rinsed three times with PBS, permeabilized with PBS, 0.5% Triton for 15 min, blocked with PBS, 0.1% Tween, 5% Fetal Calf Serum and incubated for 1h30 with specific anti-GFP (1 in 200 dilution of GFP Chicken polyclonal. #ab13970, Abcam) and anti-phospho H2AX (1 in 800 dilution of BW301, Merck) antibodies. After three PBS, 0.1% Tween washes, cells were incubated with secondary anti-chicken (Alexa Fluor® 488 Donkey Anti-Chicken. Reference: 703-546-155, Jackson Immunoresearch) and anti-mouse (Cy™3 Goat Anti-Mouse. Reference: 115-167-003, Jackson Immunoresearch) antibodies. After three PBS, 0.1% Tween washes, cells were incubated with Hoechst solution (11534886, Invitrogen) diluted 1/5000 in PBS, 0.1% Tween and mounted with ProLong™ Glass Antifade Mountant (P36982, Invitrogen). Cells were next imaged by confocal microscopy (Zeiss LSM 880) using Zenblack software and x40 lens in AiryScan mode acquisition of 7x7 contiguous XY fields and a Z-stack of 30 images at 0.1 µm intervals. Z-stacks were maximum projected and analyzed with Zen Blue software (v2.3) to automatically segment nuclei (using Hoechst staining), identify GFP-positive nuclei and count phospho-H2AX foci within each nucleus. When phospho-H2AX staining occupied more than a 1/3 of the nucleus surface, the number of foci was arbitrarily fixed as >400. Statistical significance of the difference in numbers of phospho-H2AX foci was measured with the non-parametric, rank-based Kruskal-Wallis test using GraphPad Prism (v9.3.1).

### SEM of *P. fairbanksi* adults and eggs

Adults and eggs specimens were fixed with 2.5% glutaraldehyde in Volvic mineral water for 1h and washed three times with distilled water. The adults were put in microporous capsules and the eggs were filtered on Isopore membrane filters. The samples were dehydrated in ethanol series (50%, 70%, 90% and 100%). Then critical point (Leica CPD300, PTME MNHN) was used to dry them. Adults and membranes with eggs were deposited on carbon adhesive on the SEM stubs, coated with platinum (Leica EM ACE600 coater PTME MNHN) and examined using a scanning electron microscope (Hitachi SU3500, PTME MNHN).

## ACKNOWLEDGEMENTS

We thank Dr V. Gross (University of Kassel) for advice on whole mount immunolabeling experiments, Gabriel Ramasamy (RADEXP facility, Institut Curie) for help with irradiation experiments and Nawel Cherkaoui (joint service unit CNRS UAR 3750 at Institute Pierre Gilles de Gennes) for manufacturing the brass mold for the PDMS injection chamber. Funding for the project was from Sorbonne Université (Projet Emergence, projet TardiGRaDe), INSERM, CNRS, MNHN and ANR TEFOR (ANRII-INSB-0014). M.A. was funded by a doctoral fellowship from Ministère de de l’Enseignement Supérieur et de la Recherche (France) and from Fondation pour la Recherche Médicale. The work at GenomiqueENS core facility was supported by the France Génomique national infrastructure, funded as part of the "Investissements d’Avenir" program managed by the Agence Nationale de la Recherche (contract ANR-10-INBS-0009). The proteomic experiments were partially supported by Agence Nationale de la Recherche under projects ProFI (Proteomics French Infrastructure, ANR-10-INBS-08) and GRAL, a program from the Chemistry Biology Health (CBH) Graduate School of University Grenoble Alpes (ANR-17-EURE-0003). Computing was supported by the Plateforme de Calcul Intensif et Algorithmique PCIA, Muséum national d’histoire naturelle, Centre national de la recherche scientifique, UAR 2700 2AD, Paris, France.

## Data availability

All DNA sequencing data reported in the manuscript have been deposited in the NCBI SRA database and are accessible with Bioproject ID PRJNA997229. The mass spectrometry proteomics data have been deposited to the ProteomeXchange Consortium via the PRIDE (Perez-Riverol 2022) partner repository with the dataset identifier PXD043897.

