## Supplemental Figures and supp Table 10 for "Comparative transcriptomics reveal a novel tardigrade specific DNA binding protein induced in response to ionizing radiation"

**SUPPLEMENTARY FIGURES**

**List of Supplementary Figures:**


**Supp Figure 1** Characterization of anti-phospho-H2AX antibody.
**Supp Figure 2** Analysis of DNA damage after γ-ray irradiation.
**Supp Figure 3** Abundance of *H. exemplaris* differentially expressed genes after IR and Bleomycin treatment
**Supp Figure 4** g:Profiler analysis of differentially expressed genes in *H. exemplaris* after IR.
**Supp Figure 5** Manual annotation of TDR1 gene correcting the *H. exemplaris* reference genome annotation.
**Supp Figure 6** Expression of selected proteins by western blot and quantifications.
**Supp Figure 7** Impact of cycloheximide on protein levels in *H. exemplaris* after exposure to IR
**Supp Figure 8** Abundance of differentially expressed genes of *A. antarcticus and P.fairbanksi* after IR and of *A. Antarcticus* after Bleomycin treatment.
**Supp Figure 9** DNA repair genes of major repair pathways of DNA damages caused by IR are up-regulated in all three species studied.
**Supp Figure 10** Heatmap of Log2Foldchange (ordered by adjusted p-value) of 50 genes upregulated in response to IR in all 3 species analyzed, *H. exemplaris*, *A. antarcticus* and *P. fairbanksi* and in response to Bleomycin in both *H. exemplaris* and *A. antarcticus.*
**Supp Figure 11** Production of recombinant *He*-TDR1 and *He*-TDR1-GFP.
**Supp Figure 12** Formation of aggregates of *He*-TDR1 and plasmid DNA.
**Supp Figure 13** Independent replicates of experiments of Figure 6.
**Supp Figure 14** Phylogenomics of tardigrade specific genes involved in resistance to desiccation and DNA damages (adapted from Arakawa, 2022).
**Supp Figure 15** Identification of *P. fairbanksi* tardigrades isolated and reared from moss garden.
**Supp Figure 16** Uncropped images of Western blots from Figures 1 and 3


**List of Supplementary tables:**


**Supp Table 1:** Table of differentially expressed genes after IR in *H. exemplaris (see SuppTable1_DEG_Hexe_IR.xlsx).*
**Supp Table 2**: Table of differentially expressed genes after Bleomycin treatment in *H. exemplaris (SuppTable2_DEG_Hexe_BL.xlsx).*
**Supp Table 3 :** Table of most abundant (BaseMean > 500) differentially expressed genes after IR and Bleomycin treatment in *H. exemplaris*
**Supp Table 4:** Sequences of plasmids and proteins of this study (see *SuppTable4_sequences.xlsx*).

**Supp Table 5 : BLAST and HMMER hit tables for He-TDR1 homologs** *(SuppTable5_TDR1_BLAST_HMMER.xlsx)*
**Supp Table 6**: Table of differentially expressed proteins after IR 4h or 24h post irradiation and after bleomycin treatment 5 days in *H. exemplaris (SuppTable6_DEP_Hexe_IR_BL.xlsx).*

**Supp Table 7:** Table of differentially expressed genes after IR in *A. antarcticus (SuppTable7_DEG_Aant_IR.xlsx).*
**Supp Table 8**: Table of differentially expressed genes after Bleomycin treatment in *A. antarcticus (SuppTable8_DEG_Aant_BL.xlsx).*
**Supp Table 9**: Table of differentially expressed genes after IR in *P. fairbanksi (SuppTable9_DEG_Pfai_IR.xlsx).*
**Supp Table 10**: List of Tardigrade specific proteins differentially expressed in all 3 conditions analyzed by mass spectrometry-based quantitative proteomics (4h after irradiation, 24h after irradiation and after Bleomycin treatment).
**Supp Table 11:** Mapping of RNA seq reads statistics (see *SuppTable11_RNAseq_mapping.xlsx*).
Supp Tables 1-9 and 11 are available from Zenodo repository (zenodo.org).

Supp Table 10 is at the end of this pdf file.

**a**

```
OQV18030.1      KTSNGPSQSTEHYDSQEQ---------KNPMAVKRPLAEAHENNANTQEV      161
OQV18029.1      KTSNGPSQSTEHYDSQEQ---------KNPMAVKRPLAEAHENNANTQEV      149
OQV19407.1      KSSKSDIGHSQEDRDDQDHSQSKTEHETKKKQPK--KQILQEAHVNTQEV      165
OWA54131.1      KSAGDAVEAKPAAVKTT----------K----PK--KTEAA-----SQEV      127
OQV24556.1      KTAGGPEAGEEKASASV----------KKPKAEK--KAAAA-----SQDV      151
OQV24685.1      KSAGDAVEAKP-AAVKT----------TK---PK--KTEAA-----SQEV      148
OQV20924.1      KSAGDAVEAKP-AVAKT----------AK---AK--KTEAA-----SQEV      148
OQV17567.1      KSVEGAEVAAAKVEKKP----------KS---PS--KKVEA-----SQDV      149
                *:  .                           .            :*:*
```

**b**

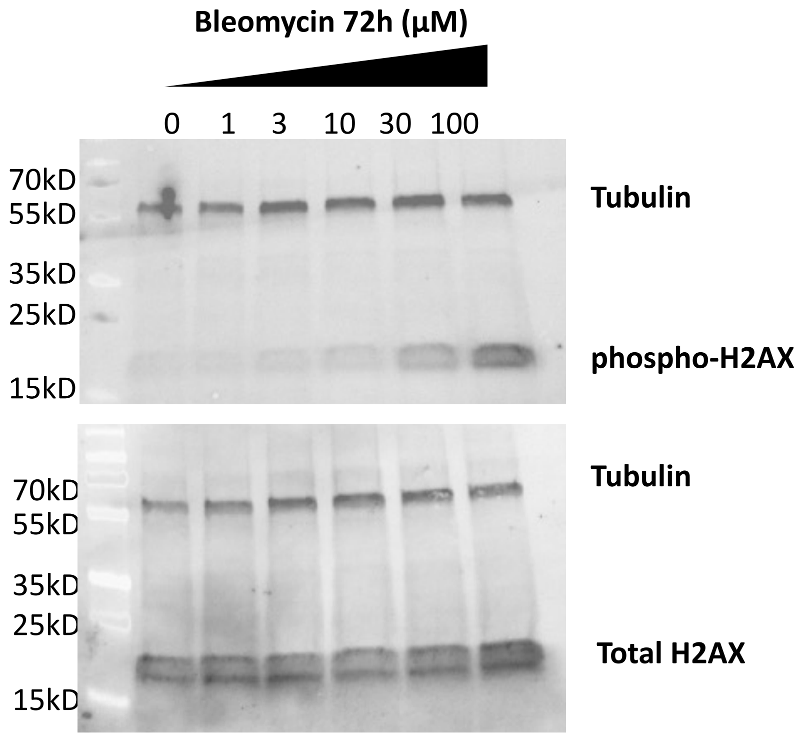


**Supp Figure 1** Characterization of anti-phospho-H2AX antibody.

**(a)** Alignment of C-terminal end of candidate H2AX homologs in *H. exemplaris*
**(b)** Western blot analysis of protein extracts from *H. exemplaris* treated with indicated concentrations of Bleomycin for 4 days using anti-phosphoH2AX raised against H2AX C-terminal peptide derived from homolog highlighted in green in (a) with phosphorylated Ser 145 and purified by negative/positive affinity chromatography to unphosphorylated/phosphorylated peptide respectively (upper panel). Unpurified antibody was used to detect total H2AX (lower panel).

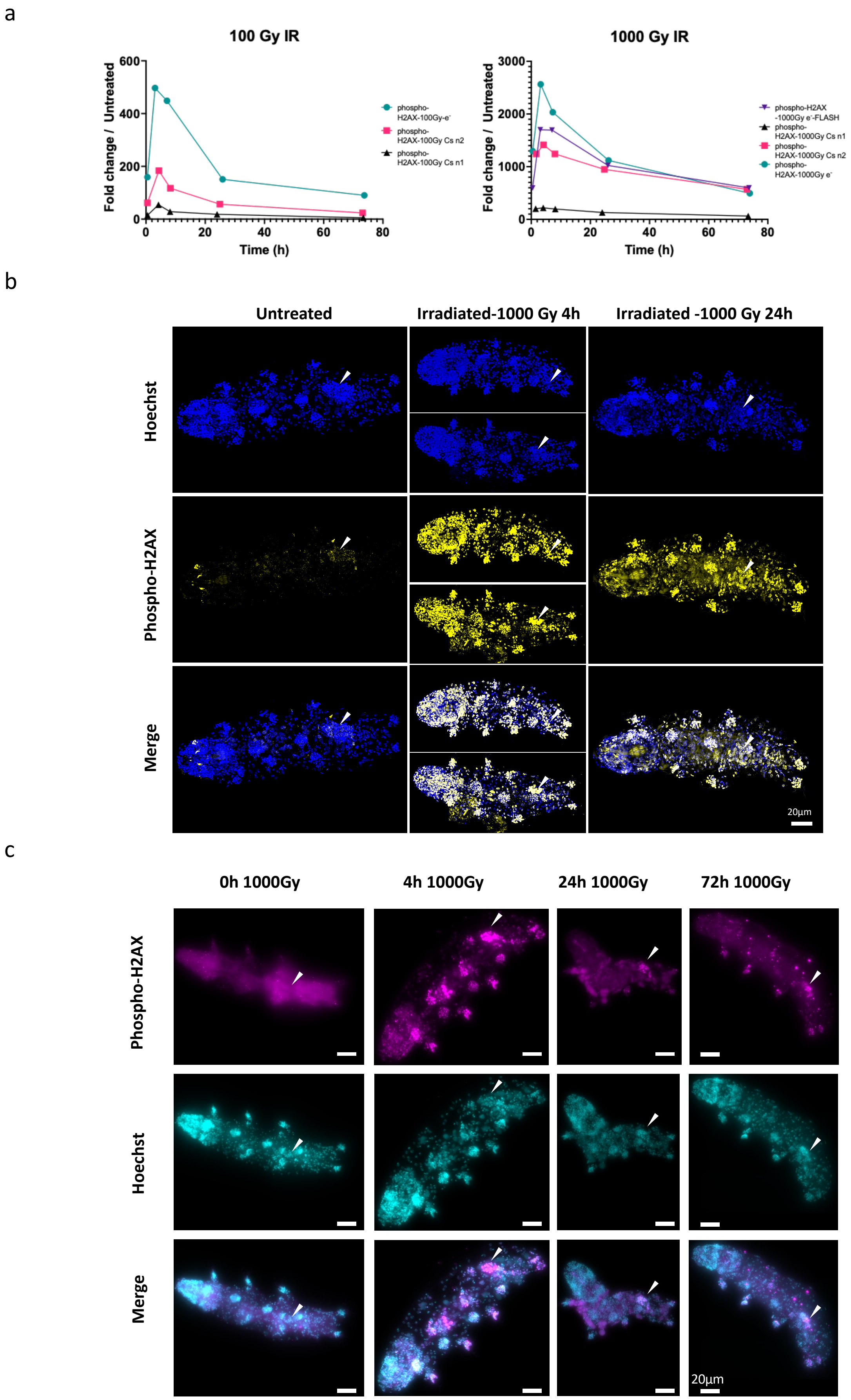


(**Supp Figure 2** Analysis of DNA damage after γ-ray irradiation)

d

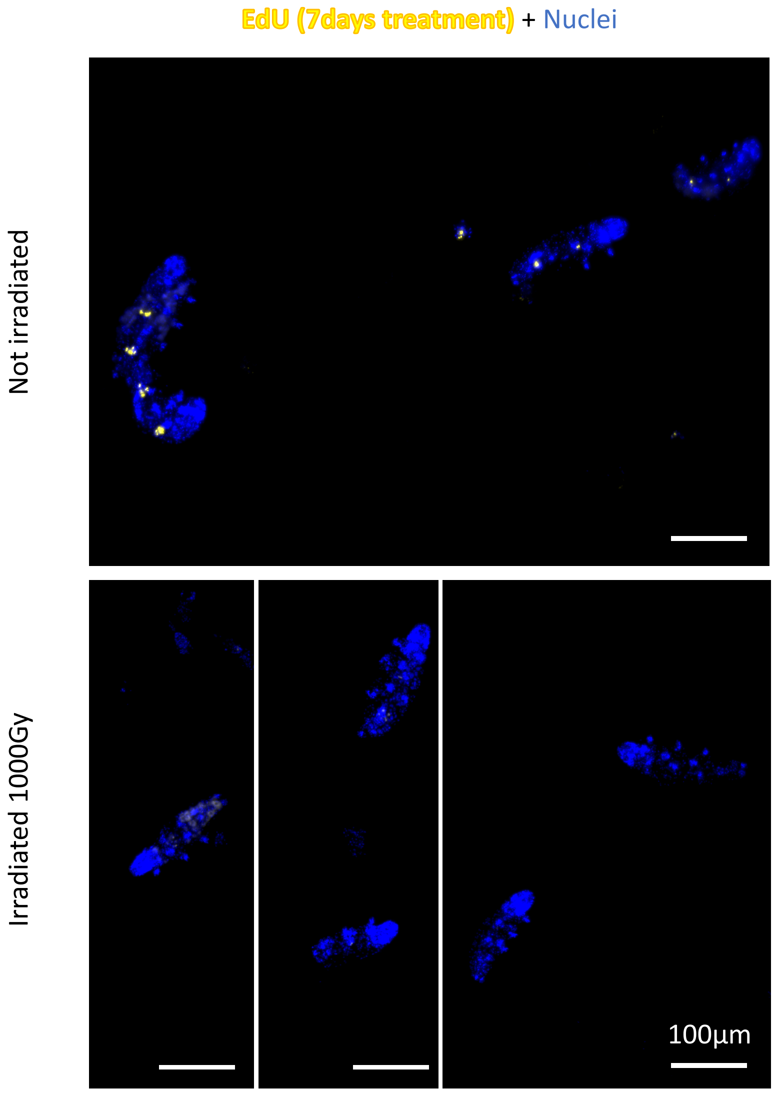


e

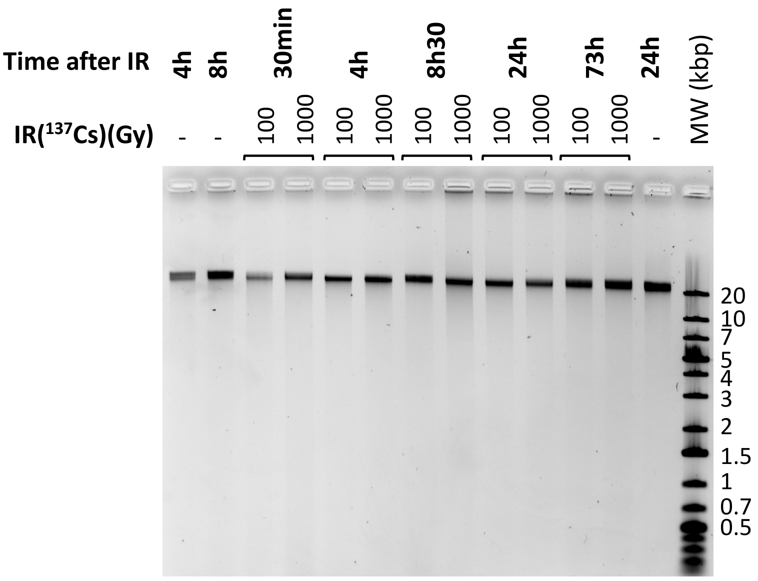


**(Supp Figure 2** Analysis of DNA damage after γ-ray irradiation)

**Supp Figure 2** Analysis of DNA damage after γ-ray irradiation.

**(a)** Quantification of phospho-H2AX after $^{137}$Cs gamma irradiation or accelerated electronic beam irradiation in the conventional mode or FLASH mode as indicated. Independent experiments labeled n1 and n2 are represented (as indicated in caption labeling). Phospho-H2AX levels from Western blots were normalized by tubulin levels and quantification is provided.
**(b)** Analysis of phospho-H2AX expression in whole mount *H. exemplaris* after exposure to 1000 Gy by confocal imaging.
*H. exemplaris* were exposed to 1000 Gy, fixed with 4% PFA at 4 and 24h post irradiation, immunolabeled with anti-phosphoH2AX antibody and anti-rabbit IgG conjugated to Alexa488 and visualized by confocal microscopy. Images at different time points were taken with identical settings so that signal intensity could be compared. Scale bar corresponds to 20 µM. Arrowhead indicates position of the gonad.
**(c)** Analysis of phospho-H2AX expression in whole mount *H . exemplaris* after exposure to 1000 Gy was also performed by standard fluorescence microscopy (enabling to readily image multiple specimens). Arrowhead indicates position of the gonad.
At 24h, while phosphoH2AX labeling had widely decreased in other tissues, strong, persistent labeling was still detected in the gonad, which likely explains that *H. exemplaris* are no longer able to lay eggs and have become sterile after 1000 Gy IR (Beltran-Pardo et al, 2005).
**(d)** Analysis of DNA synthesis in control, non-irradiated tardigrades and after 1000 Gy.
*H. exemplaris* were exposed to 1000 Gy and incubated with EdU as described by (Gross et al 2018) except for EdU concentration and time, which was 50µM EdU for 7 days, to maximize staining and sensitivity of potential DNA synthesis. After fixation with 4% PFA, EdU labeling was revealed as described (Gross et al. 2018) and imaging was performed by standard fluorescence microscopy. As shown in (Gross et al. 2018) non-irradiated, control animals exhibit intense labeling in intestinal cells whereas in irradiated tardigrades, no labeling can be detected.
**(e)** Analysis of double-strand breaks by native agarose gel electrophoresis of DNA isolated from ~8000 *H. exemplaris* individuals at indicated timepoints post-irradiation (100Gy or 1000Gy γ-rays from $^{137}$Cs source). ( - ) lanes show DNA from control, non-irradiated *H. exemplaris*.

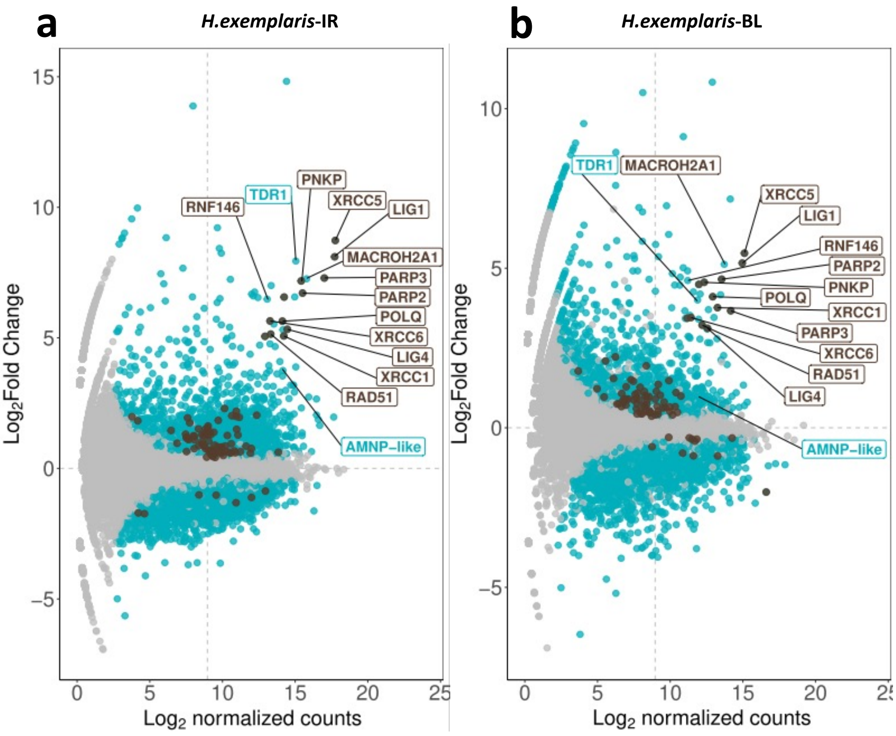


**(Supp Figure 3** Abundance of *H. exemplaris* differentially expressed genes after IR and Bleomycin treatment)

c

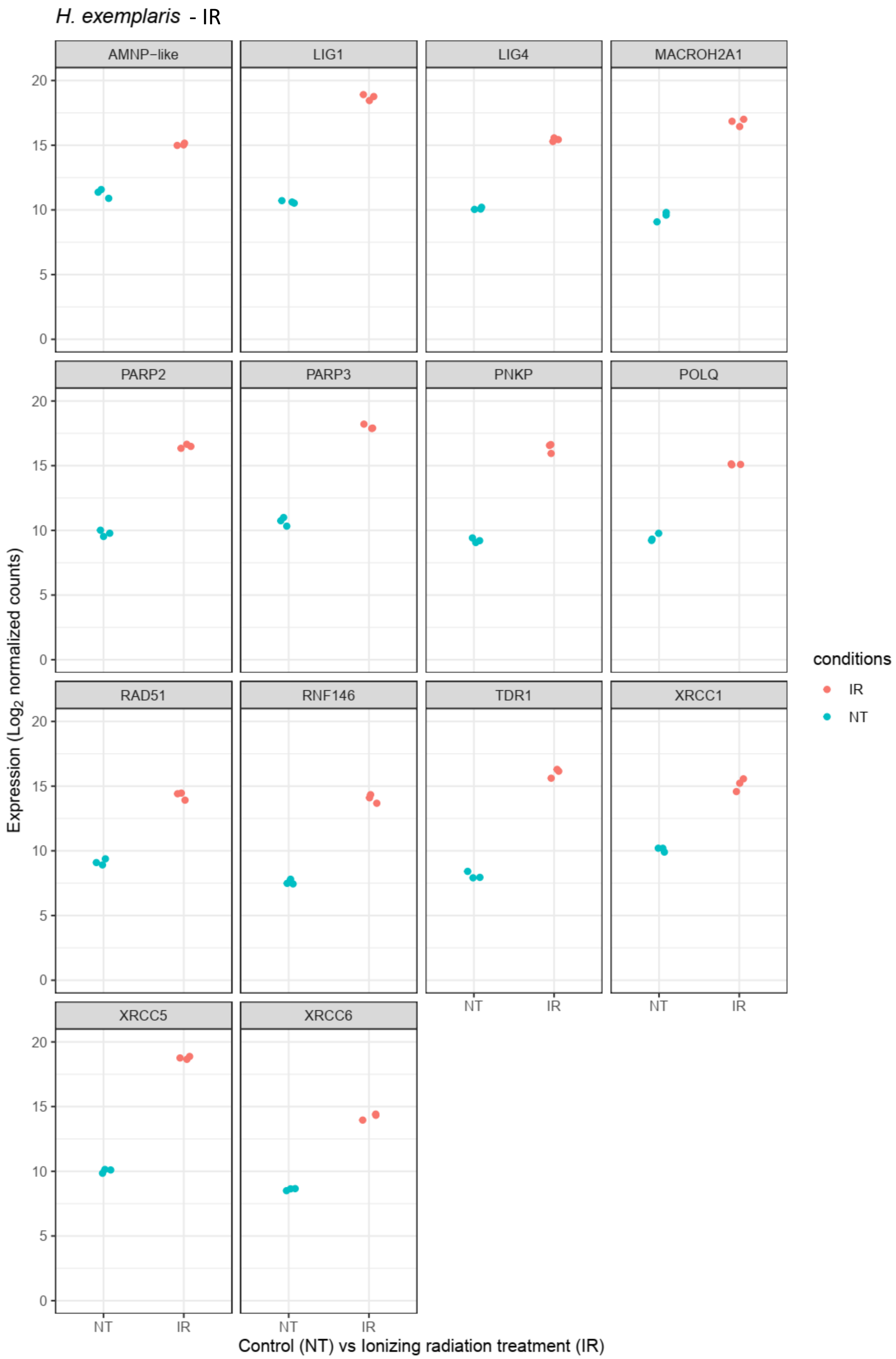


**(Supp Figure 3** Abundance of *H. exemplaris* differentially expressed genes after IR and Bleomycin treatment)

d

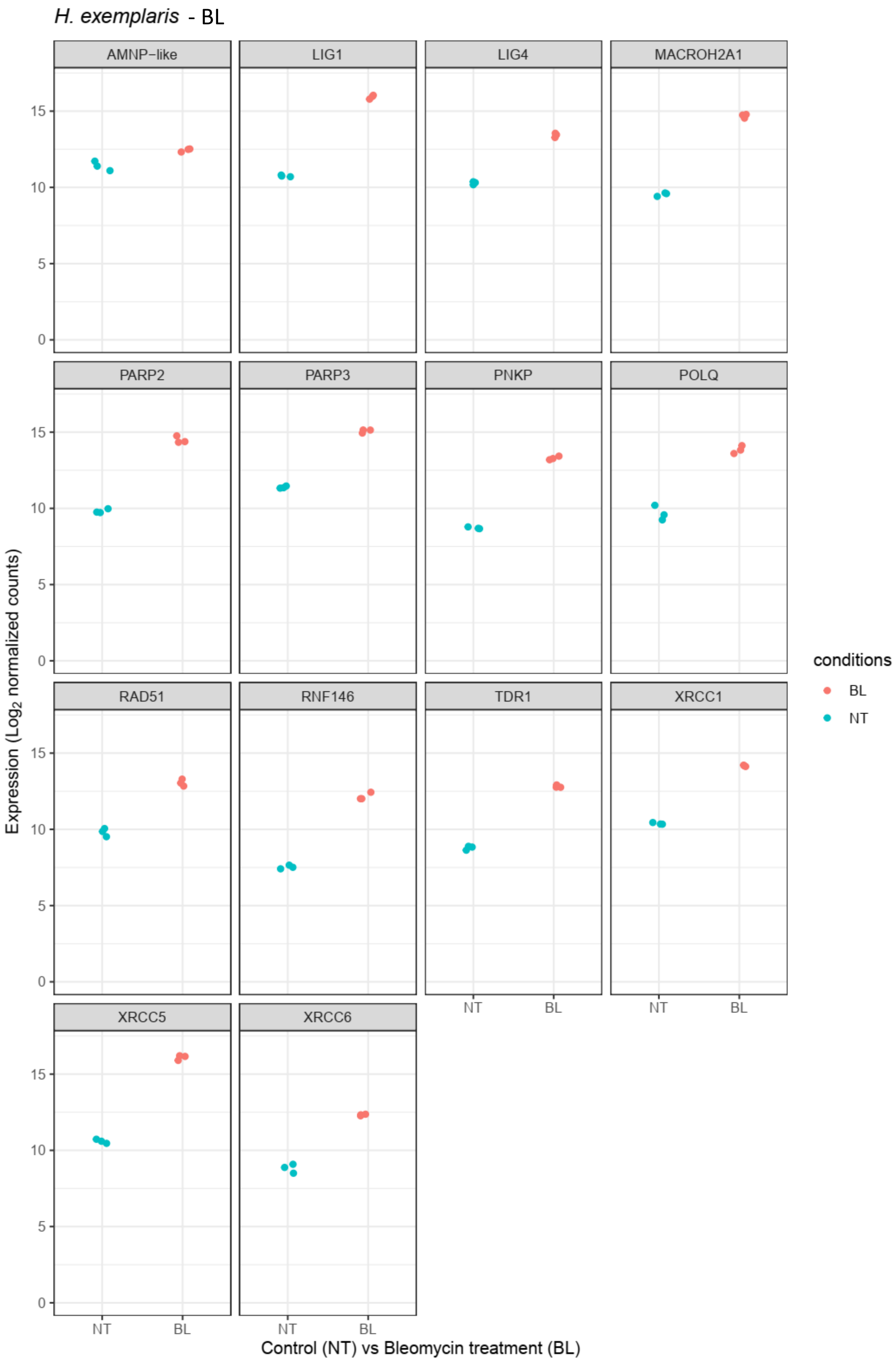


**Supp Figure 3** Abundance of *H. exemplaris* differentially expressed genes after IR and Bleomycin treatment

**(a)** MA plot of *H. exemplaris* differentially expressed genes after IR treatment
**(b)** MA plot of *H. exemplaris* differentially expressed genes after Bleomycin treatment
**(c)** Relative abundance of selected genes represented in figure 2a
**(d)** Relative abundance of selected genes represented in figure 2b

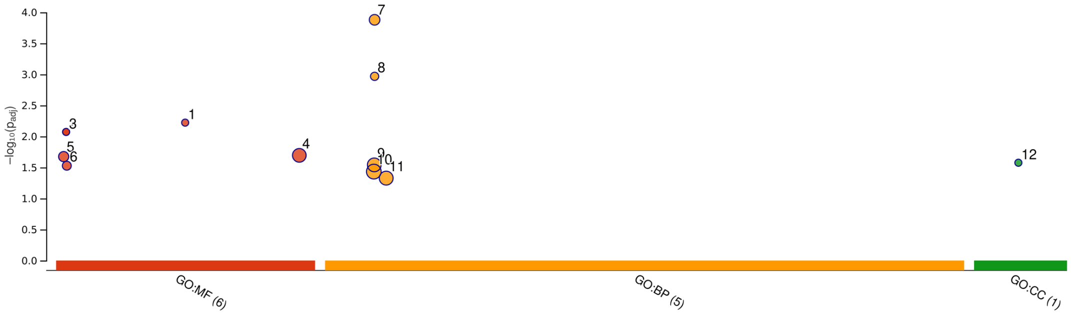


| ID | Source | Term ID | Term Name | $p_{adj}$ (query_1) |
|---|---|---|---|---|
| 1 | GO:MF | GO:0042162 | telomeric DNA binding | $5.996\times10^{-3}$ |
| 2 | GO:MF | GO:0003909 | DNA ligase activity | $8.467\times10^{-3}$ |
| 3 | GO:MF | GO:0003910 | DNA ligase (ATP) activity | $8.467\times10^{-3}$ |
| 4 | GO:MF | GO:0140097 | catalytic activity, acting on DNA | $2.018\times10^{-2}$ |
| 5 | GO:MF | GO:0003684 | damaged DNA binding | $2.114\times10^{-2}$ |
| 6 | GO:MF | GO:0003950 | NAD+ ADP-ribosyltransferase activity | $2.966\times10^{-2}$ |
| 7 | GO:BP | GO:0006302 | double-strand break repair | $1.321\times10^{-4}$ |
| 8 | GO:BP | GO:0006303 | double-strand break repair via nonhomologous … | $1.075\times10^{-3}$ |
| 9 | GO:BP | GO:0006281 | DNA repair | $2.875\times10^{-2}$ |
| 10 | GO:BP | GO:0006259 | DNA metabolic process | $3.690\times10^{-2}$ |
| 11 | GO:BP | GO:0006974 | DNA damage response | $4.700\times10^{-2}$ |
| 12 | GO:CC | GO:0043564 | Ku70:Ku80 complex | $2.655\times10^{-2}$ |

| | |
|---|---|
| **version** | e109_eg56_p17_1d3191d |
| **date** | 6/30/2023, 2:58:09 PM |
| **organism** | hegca002082055v1 |

g:Profiler

**Supp Figure 4** g:Profiler analysis of differentially expressed genes with adjusted p-value<0.05 in *H. exemplaris* after IR.

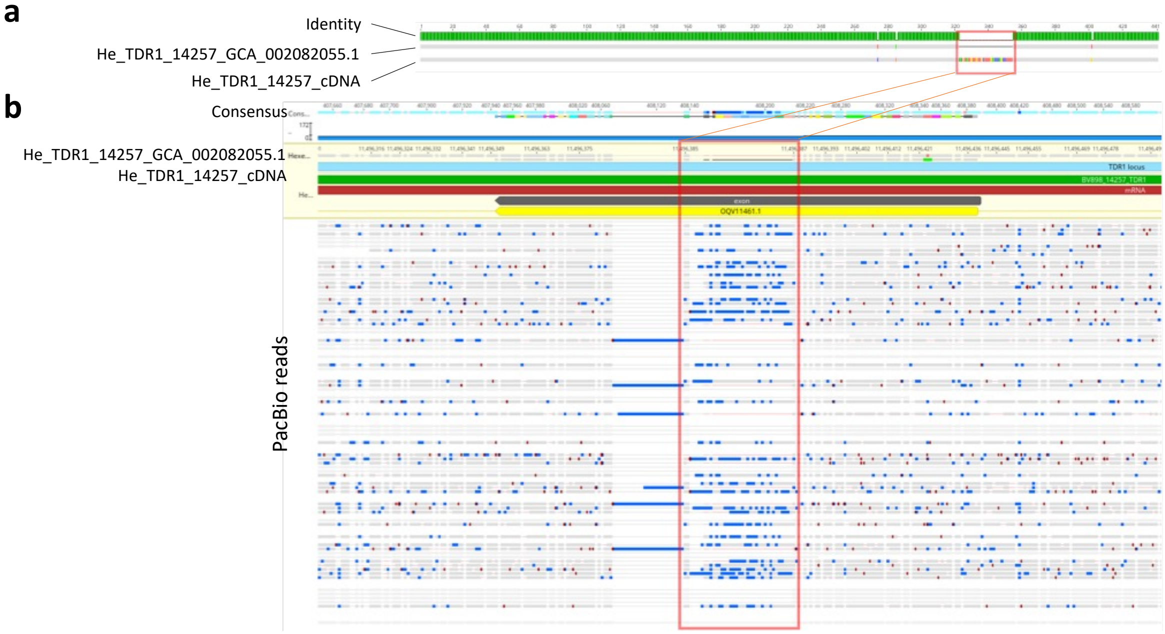


**Supp Figure 5** Manual annotation of TDR1 gene correcting the *H. exemplaris* reference genome annotation.

**(a)** Alignment of *H. exemplaris* genome assembly GCA_002082055.1 with cDNA sequence of *He*-TDR1 obtained from ONT long read sequencing and cDNA cloning showed that a portion of TDR1 sequence is missing in the current assembly
**(b)** Alignment of PacBio reads used for genome assembly with *H. exemplaris* genome assembly GCA_002082055.1 and *He*-TDR1 cDNA.

A zoom on the missing sequence (boxed in orange) shows the poor quality of PacBio reads used for genome assembly at this locus, likely explaining the absence of the missing *He*-TDR1 cDNA sequence in the current genome assembly. PacBio reads (SRX2495681, Yoshida et al. 2017) were downloaded from NCBI, mapped with minimap2 (Li 2018) and alignment visualization was performed with Geneious Prime (v2023.1). Blue and red dots respectively indicate mismatches and indels in the alignment.
cDNA sequence of *He*-TDR1 is provided in Supp Table 8 and encodes for a 146 amino acids long protein.

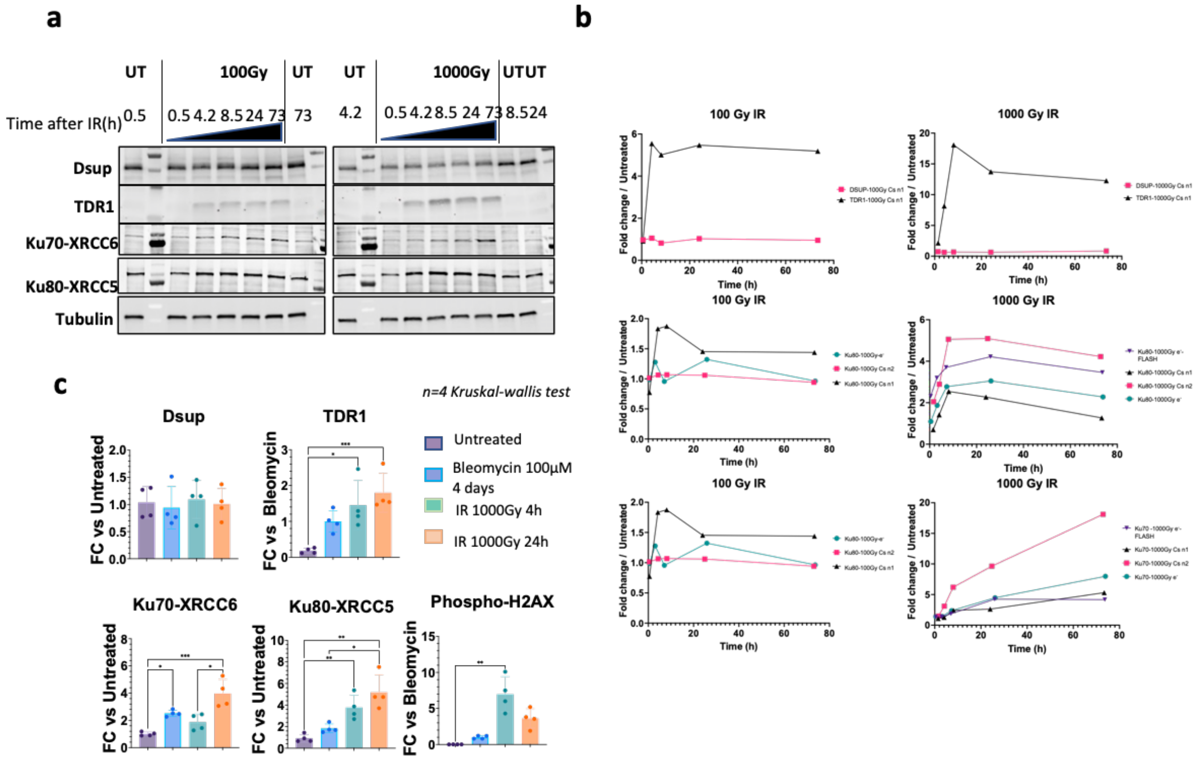


**Supp Figure 6:** Expression of selected proteins by Western blot and quantifications.

**(a)** Kinetics of protein expression after $\gamma$-irradiation (100Gy (left-side panel) and 1000Gy (right-side panel) assessed by Western blots with antibodies against *He*Dsup, *He*TDR1, *He*Ku70-XRCC6, *He*Ku80-XRCC5 and $\alpha$-Tubulin for normalization.
**(b)** Quantification of Western blots from experiments carried out with a Cesium source (two independent experiments, labeled Csn1and Csn2 respectively, were performed) or with accelerated electron beam in conventional electron mode or FLASH electron mode as indicated. Signal is normalized with $\alpha$-Tubulin. Western blots from experiment Csn1 are shown in (a).
**(c)** Quantification of Western blots of samples used for differential proteomic analysis reported in Figure 3 (normalized to $\alpha$-Tubulin signal) n=4 independent experiments. FC stands for Fold Change. Kruskal-wallis statistical test: *, p-value<0.05; **, p-value<0.005; ***, p-value<0.0005.

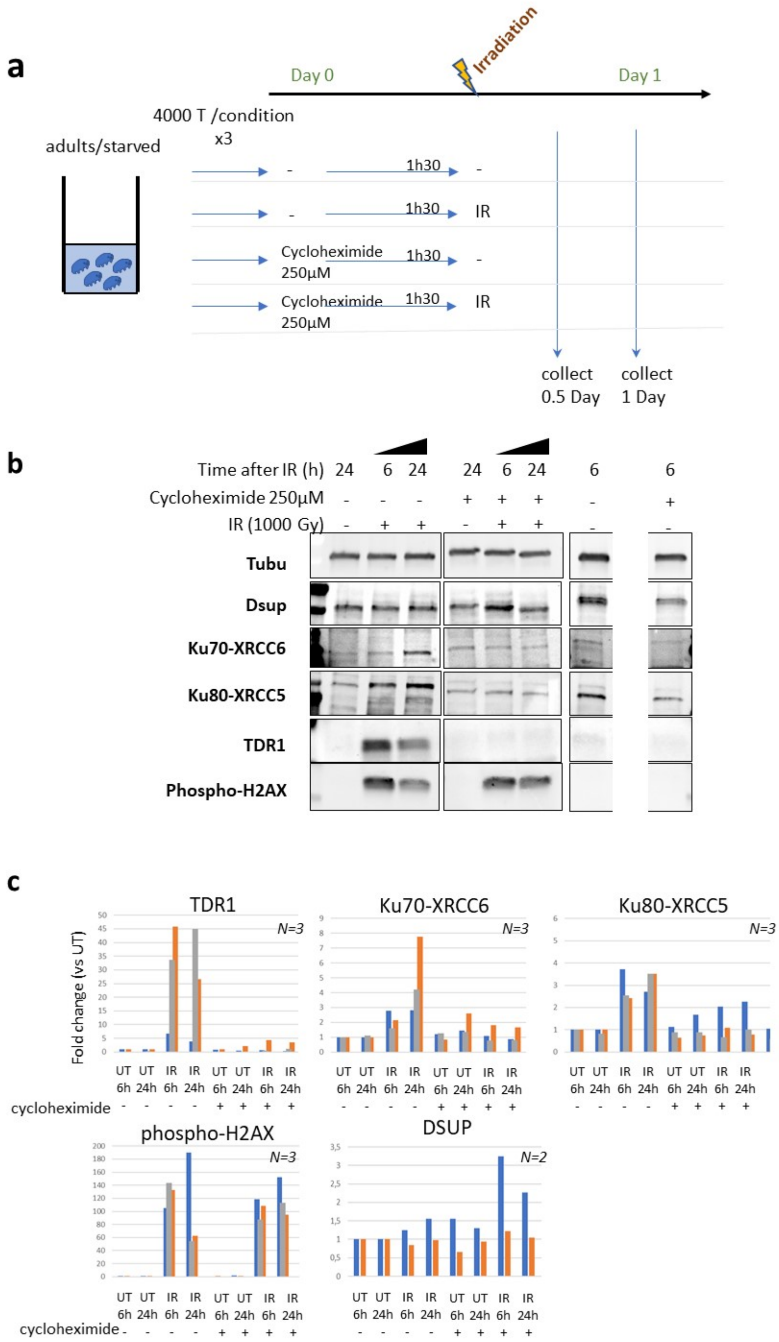


**Supp Figure 7** Impact of cycloheximide on protein levels in *H. exemplaris* after exposure to IR.

**(a)** Schematic showing experimental set up used to determine the impact of cycloheximide on protein levels in *H. exemplaris* after exposure to IR.

**(b)** Western blot analysis of TDR1, XRCC5, XRCC6 (among the most strongly stimulated genes at the RNA level) and Dsup (not stimulated at the RNA level) in irradiated *H. exemplaris* tardigrades untreated ( - ) or treated with 1000Gy γ-rays and extracts prepared at indicated times after irradiation.

**(c)** Quantification of Western blots (normalized to $\alpha$-tubulin signal).

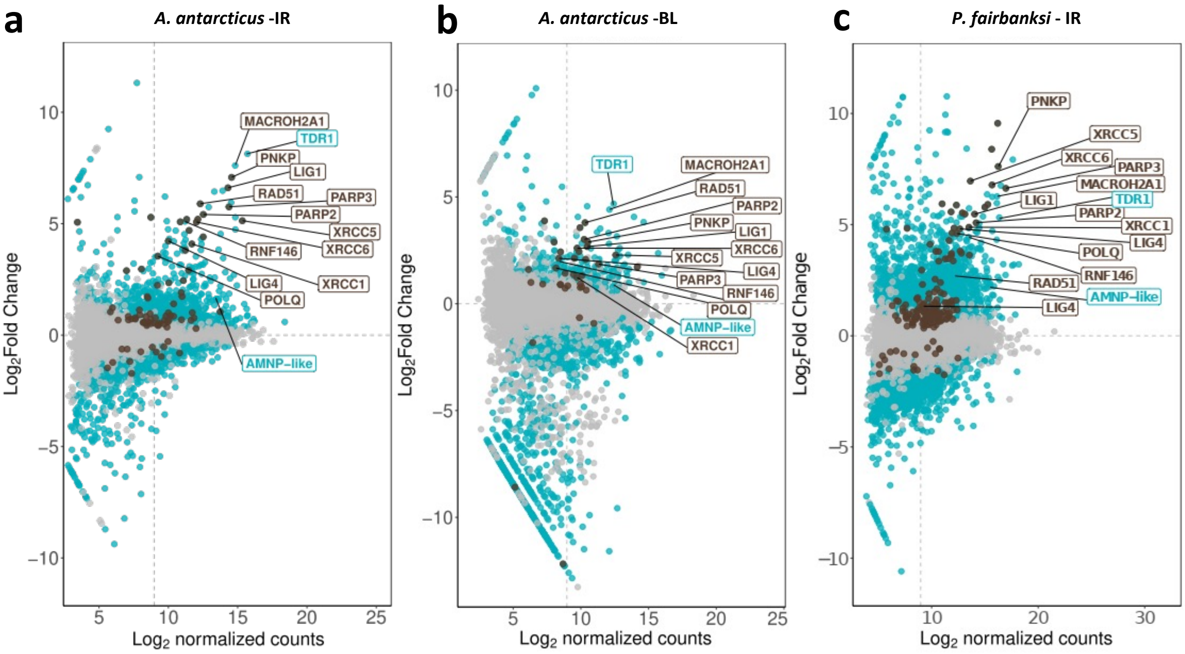


**(Supp Figure 8** Abundance of differentially expressed genes of *A. antarcticus and P.fairbanksi* after IR and of *A. Antarcticus* after Bleomycin treatment*)***.**

**d**

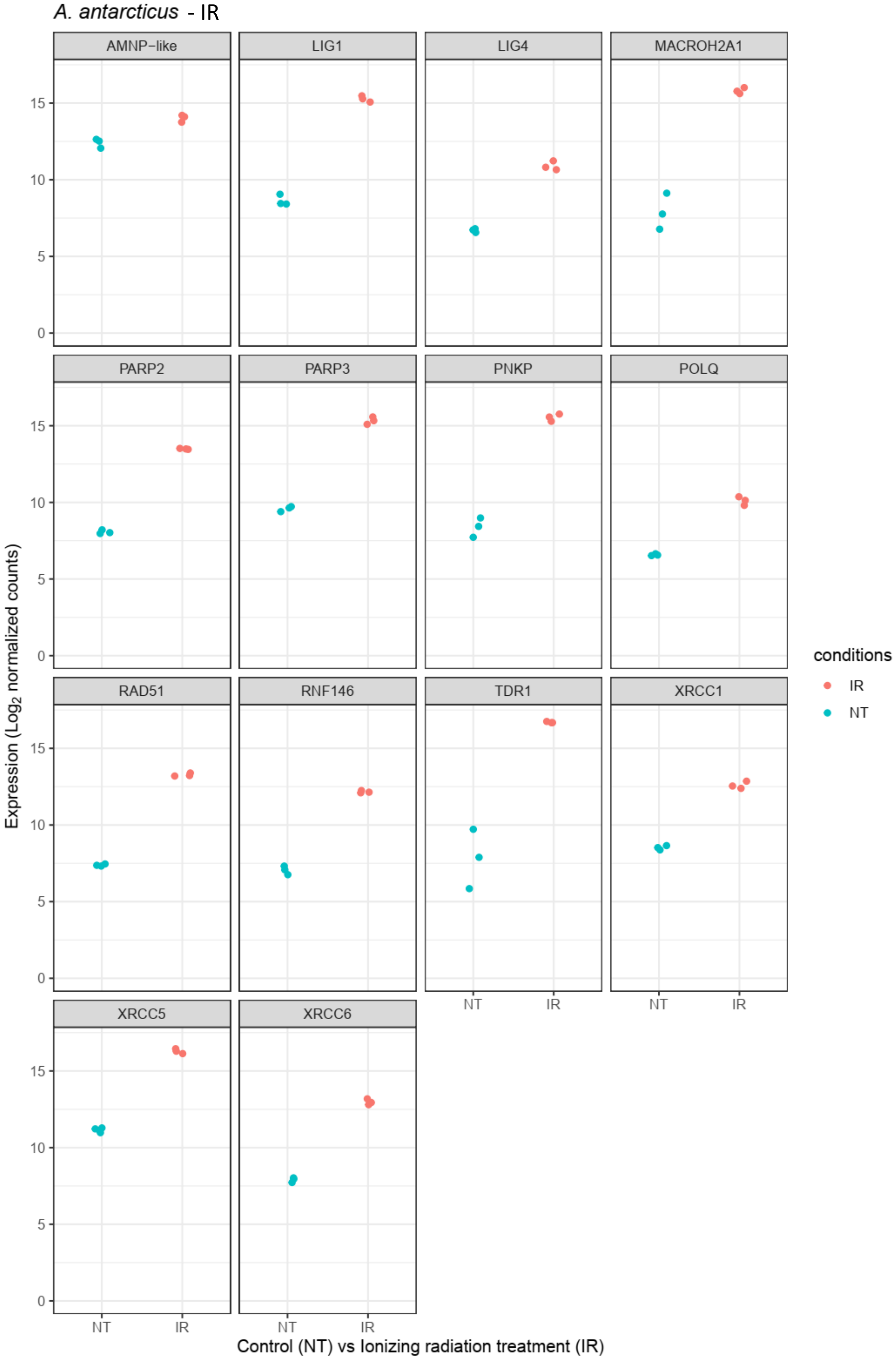


**(Supp Figure 8** Abundance of differentially expressed genes of *A. antarcticus and P.fairbanksi* after IR and of *A. Antarcticus* after Bleomycin treatment*)*.

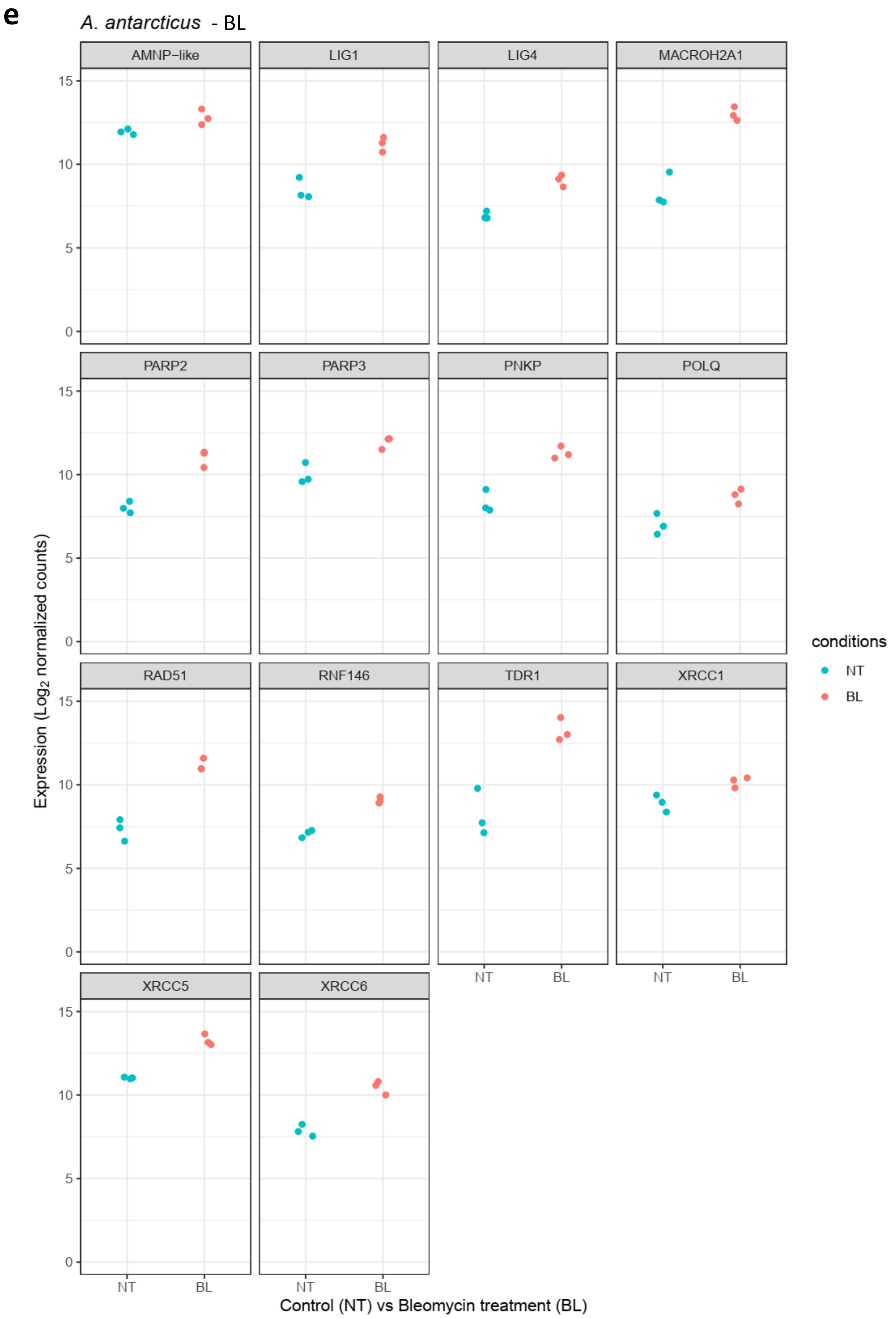


**(Supp Figure 8** Abundance of differentially expressed genes of *A. antarcticus and P.fairbanksi* after IR and of *A. Antarcticus* after Bleomycin treatment*)***.**

f

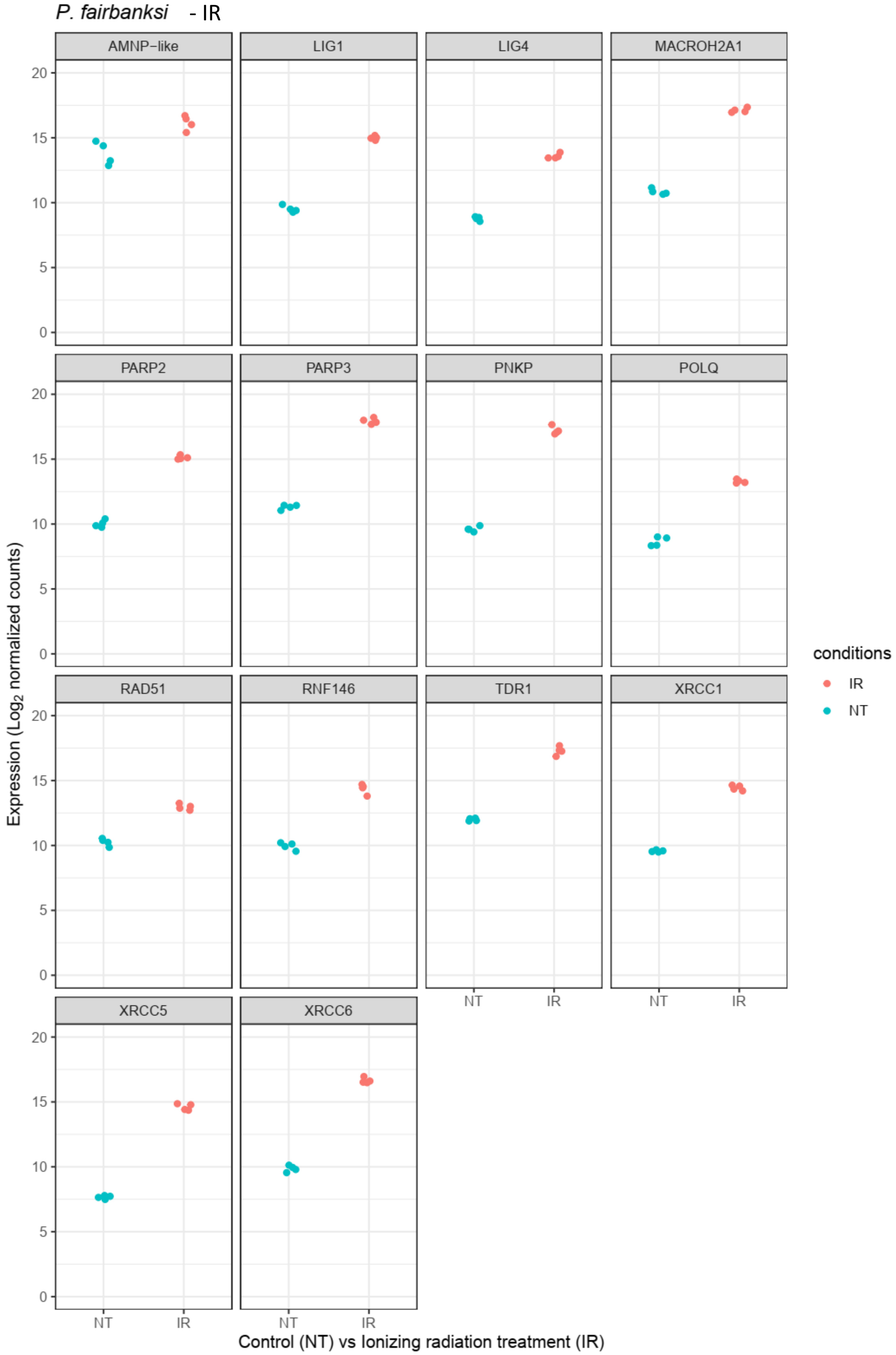


**(Supp Figure 8** Abundance of differentially expressed genes of *A. antarcticus and P.fairbanksi* after IR and of *A. Antarcticus* after Bleomycin treatment*)***.**

**Supp Figure 8** Abundance of differentially expressed genes of *A. antarcticus and P.fairbanksi* after IR and of *A. Antarcticus* after Bleomycin treatmen*t***.**

**(a)** MA plot of *A. antarcticus* differentially expressed genes after IR treatment
**(b)** MA plot of *A. antarcticus* differentially expressed genes after Bleomycin treatment
**(c)** MA plot of *P. fairbanksi* differentially expressed genes after IR treatment
**(d)** Relative abundance of selected genes represented in figure 4a and Supp Figure 8a
**(e)** Relative abundance of selected genes represented in Supp Figure 8b
**(f)** Relative abundance of selected genes represented in figure 4b and Supp Figure 8c

**a**

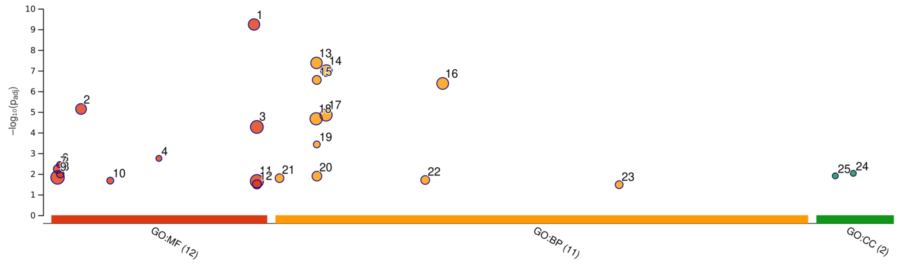


| ID | Source | Term ID | Term Name | $p_{adj}$ (query_1) |
|---|---|---|---|---|
| 1 | GO:MF | GO:0140097 | catalytic activity, acting on DNA | $5.827\times10^{-10}$ |
| 2 | GO:MF | GO:0008094 | ATP-dependent activity, acting on DNA | $7.179\times10^{-6}$ |
| 3 | GO:MF | GO:0140640 | catalytic activity, acting on a nucleic acid | $5.338\times10^{-5}$ |
| 4 | GO:MF | GO:0042162 | telomeric DNA binding | $1.768\times10^{-3}$ |
| 5 | GO:MF | GO:0003909 | DNA ligase activity | $3.542\times10^{-3}$ |
| 6 | GO:MF | GO:0003910 | DNA ligase (ATP) activity | $3.542\times10^{-3}$ |
| 7 | GO:MF | GO:0003684 | damaged DNA binding | $5.753\times10^{-3}$ |
| 8 | GO:MF | GO:0003950 | NAD+ ADP-ribosyltransferase activity | $1.023\times10^{-2}$ |
| 9 | GO:MF | GO:0003677 | DNA binding | $1.487\times10^{-2}$ |
| 10 | GO:MF | GO:0016886 | ligase activity, forming phosphoric ester bonds | $2.113\times10^{-2}$ |
| 11 | GO:MF | GO:0140657 | ATP-dependent activity | $2.237\times10^{-2}$ |
| 12 | GO:MF | GO:0140658 | ATP-dependent chromatin remodeler activity | $3.221\times10^{-2}$ |
| 13 | GO:BP | GO:0006281 | DNA repair | $4.234\times10^{-8}$ |
| 14 | GO:BP | GO:0006974 | DNA damage response | $9.943\times10^{-8}$ |
| 15 | GO:BP | GO:0006302 | double-strand break repair | $2.828\times10^{-7}$ |
| 16 | GO:BP | GO:0033554 | cellular response to stress | $4.177\times10^{-7}$ |
| 17 | GO:BP | GO:0006950 | response to stress | $1.399\times10^{-5}$ |
| 18 | GO:BP | GO:0006259 | DNA metabolic process | $2.138\times10^{-5}$ |
| 19 | GO:BP | GO:0006303 | double-strand break repair via nonhomologous ... | $3.738\times10^{-4}$ |
| 20 | GO:BP | GO:0006310 | DNA recombination | $1.280\times10^{-2}$ |
| 21 | GO:BP | GO:0000723 | telomere maintenance | $1.604\times10^{-2}$ |
| 22 | GO:BP | GO:0032200 | telomere organization | $1.967\times10^{-2}$ |
| 23 | GO:BP | GO:0071897 | DNA biosynthetic process | $3.324\times10^{-2}$ |
| 24 | GO:CC | GO:0043564 | Ku70:Ku80 complex | $9.374\times10^{-3}$ |
| 25 | GO:CC | GO:0030870 | Mre11 complex | $1.242\times10^{-2}$ |

**version** e109_eg56_p17_1d3191d
**date** 6/27/2023, 9:55:36 AM
**organism** hegca002082055v1

g:Profiler

**b**

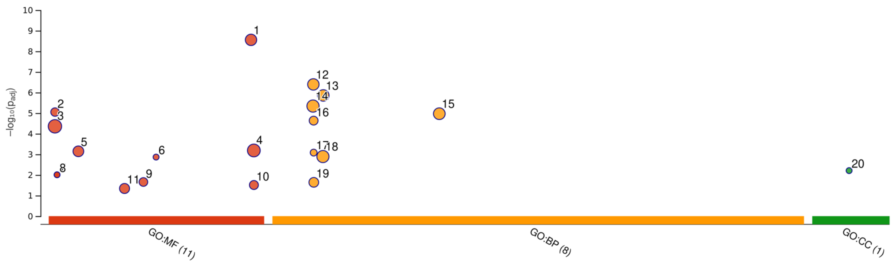


| ID | Source | Term ID | Term Name | $p_{adj}$ (query_1) |
|---|---|---|---|---|
| 1 | GO:MF | GO:0140097 | catalytic activity, acting on DNA | $2.786\times10^{-9}$ |
| 2 | GO:MF | GO:0003684 | damaged DNA binding | $8.895\times10^{-6}$ |
| 3 | GO:MF | GO:0003677 | DNA binding | $4.371\times10^{-5}$ |
| 4 | GO:MF | GO:0140640 | catalytic activity, acting on a nucleic acid | $6.492\times10^{-4}$ |
| 5 | GO:MF | GO:0008094 | ATP-dependent activity, acting on DNA | $7.142\times10^{-4}$ |
| 6 | GO:MF | GO:0042162 | telomeric DNA binding | $1.360\times10^{-3}$ |
| 7 | GO:MF | GO:0003910 | DNA ligase (ATP) activity | $9.840\times10^{-3}$ |
| 8 | GO:MF | GO:0003909 | DNA ligase activity | $9.840\times10^{-3}$ |
| 9 | GO:MF | GO:0034061 | DNA polymerase activity | $2.203\times10^{-2}$ |
| 10 | GO:MF | GO:0140658 | ATP-dependent chromatin remodeler activity | $3.083\times10^{-2}$ |
| 11 | GO:MF | GO:0030527 | structural constituent of chromatin | $4.549\times10^{-2}$ |
| 12 | GO:BP | GO:0006281 | DNA repair | $4.044\times10^{-7}$ |
| 13 | GO:BP | GO:0006974 | DNA damage response | $1.389\times10^{-6}$ |
| 14 | GO:BP | GO:0006259 | DNA metabolic process | $4.551\times10^{-6}$ |
| 15 | GO:BP | GO:0033554 | cellular response to stress | $1.069\times10^{-5}$ |
| 16 | GO:BP | GO:0006302 | double-strand break repair | $2.318\times10^{-5}$ |
| 17 | GO:BP | GO:0006303 | double-strand break repair via nonhomologous ... | $8.244\times10^{-4}$ |
| 18 | GO:BP | GO:0006950 | response to stress | $1.281\times10^{-3}$ |
| 19 | GO:BP | GO:0006310 | DNA recombination | $2.291\times10^{-2}$ |
| 20 | GO:CC | GO:0043564 | Ku70:Ku80 complex | $6.174\times10^{-3}$ |

**version** e109_eg56_p17_1d3191d
**date** 6/27/2023, 10:03:30 AM
**organism** hegca002082055v1

g:Profiler

**c**

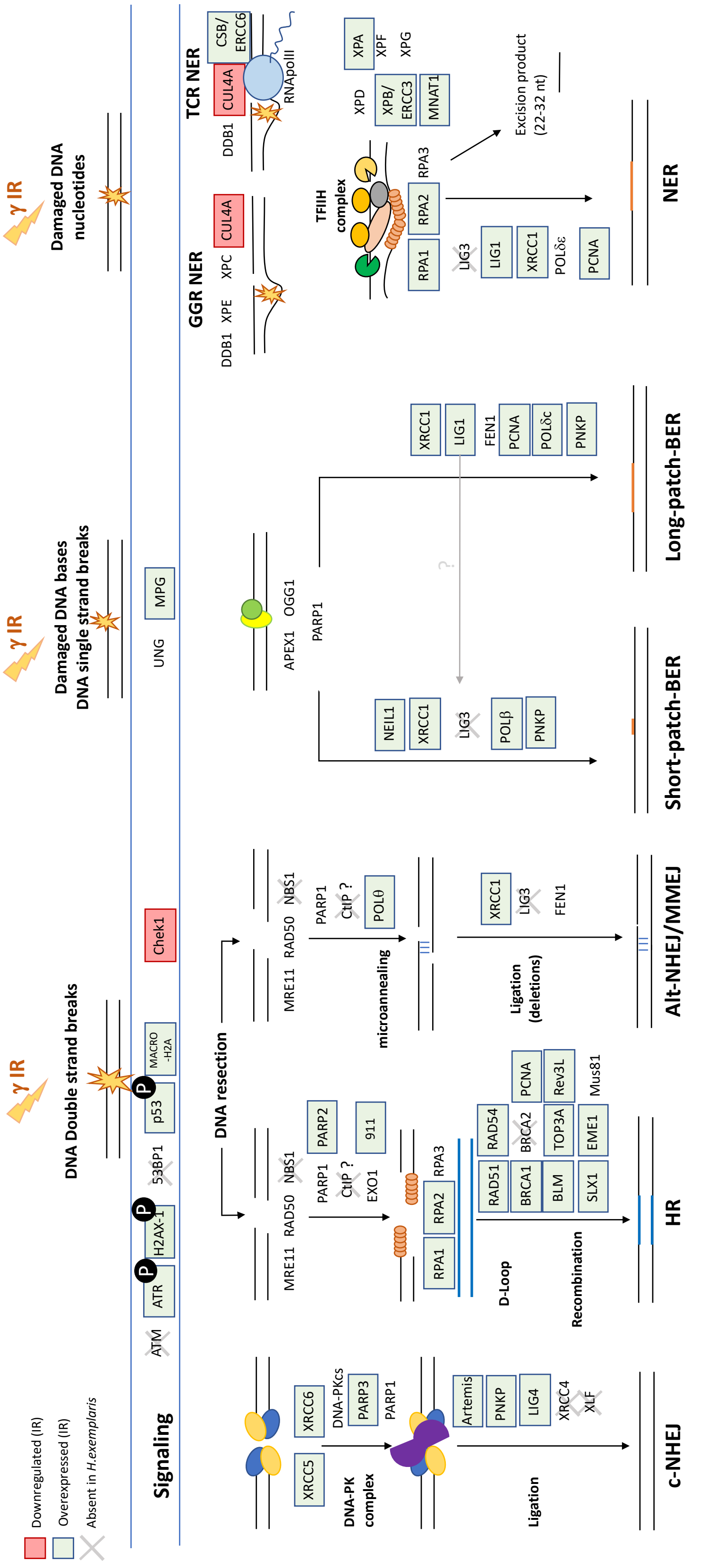

Downregulated (IR)
Overexpressed (IR)
Absent in H.exemplaris
γ IR
DNA Double strand breaks
γ IR
Damaged DNA bases
DNA single strand breaks
γ IR
Damaged DNA nucleotides
Signaling
ATM
ATR
H2AX-1
53BP1
p53
MACRO -H2A
Chek1
UNG
MPG
DNA resection
DNA-PK complex
XRCC5
XRCC6
DNA-PKcs
PARP3
PARP1
Artemis
PNKP
LIG4
XRCC4
XLF
Ligation
c-NHEJ
MRE11
RAD50
NBS1
PARP1
CtIP ?
EXO1
PARP2
911
RPA1
RPA2
RPA3
D-Loop
RAD51
RAD54
BRCA1
BRCA2
PCNA
BLM
TOP3A
Rev3L
SLX1
EME1
Mus81
Recombination
HR
POLθ
microannealing
XRCC1
LIG3
FEN1
Ligation (deletions)
Alt-NHEJ/MMEJ
APEX1
OGG1
PARP1
NEIL1
POLβ
Short-patch-BER
LIG1
PCNA
POLδε
Long-patch-BER
GGR NER
DDB1
XPE
XPC
CUL4A
TCR NER
CSB/ ERCC6
RNApolII
TFIIH complex
XPD
XPB/ ERCC3
MNAT1
XPA
XPF
XPG
Excision product (22-32 nt)
NER

**d**

Differentially expressed genes

| term_name | Base excision repair | | | Nucleotide excision repair | | | Mismatch repair | | | Homologous recombination | | | Non-homologous end-joining | | | Fanconi anemia pathway | | |
|---|---|---|---|---|---|---|---|---|---|---|---|---|---|---|---|---|---|---|
| term_id | KEGG:03410 | | | KEGG:03420 | | | KEGG:03430 | | | KEGG:03440 | | | KEGG:03450 | | | KEGG:03460 | | |
| *Species* | *H.exemplaris* | *A.antarcticus* | *P.fairbanksi* | *H.exemplaris* | *A.antarcticus* | *P.fairbanksi* | *H.exemplaris* | *A.antarcticus* | *P.fairbanksi* | *H.exemplaris* | *A.antarcticus* | *P.fairbanksi* | *H.exemplaris* | *A.antarcticus* | *P.fairbanksi* | *H.exemplaris* | *A.antarcticus* | *P.fairbanksi* |
| APEX1 | | APEX1 | | | | | | | | | | | | | | ATR | | ATR |
| APEX2 | | APEX2 | | | | | | | | | | | | | | | | |
| ATR | | | | | | | | | | | | | | | | | | |
| BLM | | | | | | | | | | BLM | BLM | BLM | | | | BLM | BLM | BLM |
| BRIP1 | | | | | | | | | | BRIP1 | | BRIP1 | | | | BRIP1 | | BRIP1 |
| CCNH | | | | CCNH | | | | | | | | | | | | | | |
| CDK7 | | | | CDK7 | CDK7 | CDK7 | | | | | | | | | | | | |
| CUL4A | | | | | CUL4A | CUL4A | | | | | | | | | | | | |
| DCLRE1C | | | | | | | | | | | | | DCLRE1C | | | | | |
| DDB1 | | | | *DDB1* | | DDB1 | | | | | | | | | | | | |
| ERCC1 | | | | ERCC1 | | | | | | | | | | | | ERCC1 | | |
| ERCC2 | | | | | ERCC2 | | | | | | | | | | | | | |
| ERCC3 | | | | ERCC3 | ERCC3 | ERCC3 | | | | | | | | | | | | |
| ERCC4 | | | | ERCC4 | | ERCC4 | | | | | | | | | | ERCC4 | | ERCC4 |
| ERCC5 | | | | | | ERCC5 | | | | | | | | | | | | |
| ERCC6 | | | | ERCC6 | ERCC6 | ERCC6 | | | | | | | | | | | | |
| FAN1 | | | | | | | | | | | | | | | | FAN1 | | |
| FANCM | | | | | | | | | | | | | | | | | FANCM | FANCM |
| FEN1 | | | FEN1 | | | | | | | | | | | | FEN1 | | | |
| GTF2H1 | | | | GTF2H1 | | | | | | | | | | | | | | |
| GTF2H2 | | | | GTF2H2 | | GTF2H1 | | | | | | | | | | | | |
| GTF2H3 | | | | | GTF2H3 | GTF2H3 | | | | | | | | | | | | |
| GTF2H4 | | | | GTF2H4 | | GTF2H4 | | | | | | | | | | | | |
| GTF2H5 | | | | GTF2H5 | | | | | | | | | | | | | | |
| HES1 | | | | | | | | | | | | | | | | | | HES1 |
| HMGB1 | HMGB1 | HMGB1 | HMGB1 | | | | | | | | | | | | | | | |
| LIG1 | LIG1 | LIG1 | LIG1 | LIG1 | LIG1 | LIG1 | LIG1 | LIG1 | LIG1 | | | | | | | | | |
| LIG4 | | | | | POLD1 | | | | | | | | LIG4 | LIG4 | LIG4 | | | |
| MLH1 | | | | | POLD2 | | | | MLH1 | | | | | | | | | MLH1 |
| MLH3 | | | | | POLD4 | | | | MLH3 | | | | | | | | | |
| MNAT1 | | | | MNAT1 | | MNAT1 | | | | | | | | | | | | |
| MPG | MPG | | | | | | | | | | | | | | | | | |
| MRE11A | | | | | | | | | | | MRE11A | MRE11A | | MRE11A | MRE11A | | | |
| MSH2 | | | | | | | | MSH2 | MSH2 | | | | | | | | | |
| MUS81 | | | | | | | | | | | MUS81 | MUS81 | | | | | MUS81 | MUS81 |
| NEIL1 | NEIL1 | NEIL1 | NEIL1 | | | | | | | | | | | | | | | |
| OGG1 | | OGG1 | | | | | | | | | | | | | | | | |
| PARP1 | | | PARP1 | | | | | | | | | | | | | | | |
| PARP2 | PARP2 | PARP2 | PARP2 | | | | | | | | | | | | | | | |
| PARP3 | PARP3 | PARP3 | PARP3 | | | | | | | | | | | | | | | |
| PARP4 | PARP4 | | | | | | | | | | | | | | | | | |
| PCNA | PCNA | | PCNA | PCNA | | PCNA | PCNA | | PCNA | | | | | | | | | |
| POLB | | | POLB | | | | POLD1 | POLD1 | POLD1 | | | | | | | | | |
| POLD1 | POLD1 | POLD1 | POLD1 | POLD1 | | POLD1 | POLD2 | POLD2 | POLD2 | POLD1 | POLD1 | POLD1 | | | | | | |
| POLD2 | POLD2 | POLD2 | POLD2 | POLD2 | | POLD2 | | | POLD3 | POLD2 | POLD2 | POLD2 | | | | | | |
| POLD3 | | | POLD3 | | | POLD3 | POLD4 | POLD4 | | | | POLD3 | | | | | | |
| POLD4 | POLD4 | POLD4 | | POLD4 | | | | | | POLD4 | POLD4 | | | | | | | |
| POLE | | | POLE | | | POLE | | | | | | | | | | | | |
| POLE2 | | | POLE2 | | | POLE2 | | | | | | | | | | | | |
| POLE4 | POLE4 | | | POLE4 | | | | | | | | | | | | | | |
| POLH | | | | | | | | | | | | | | | | | POLH | POLH |
| POLK | | | | | | | | | | | | | | | | POLK | POLK | POLK |
| POLL | POLL | POLL | POLL | | | | | | | | | | POLL | POLL | POLL | | | |
| PRKDC | | | | | | | | | | | | | | | PRKDC | | | |
| RAD23A | | | | | RAD23A | RAD23A | | | | | | | | | | | | |
| RAD23B | | | | RAD23B | | RAD23B | | | | | | | | RAD50 | RAD50 | | | |
| RAD50 | | | | | | | | | | | RAD50 | RAD50 | | | | | | |
| RAD51 | | | | | | | | | | RAD51 | RAD51 | *RAD51* | | | | RAD51 | RAD51 | *RAD51* |
| RAD54B | | | | | | | | | | | RAD54B | | | | | | | |
| RAD54L | | | | | | | | | | RAD54L | RAD54L | RAD54L | | | | | | |
| REV1 | | | | | | | | | | | | | | | | REV1 | REV1 | REV1 |
| REV3L | | | | | | | | | | | | | | | | REV3L | REV3L | REV3L |
| RFC1 | | | | RFC1 | RFC1 | RFC1 | RFC1 | RFC1 | RFC1 | | | | | | | | | |
| RFC4 | | | | | | RFC4 | | | RFC4 | | | | | | | | | |
| RFC5 | | | | RFC5 | | RFC5 | RFC5 | | RFC5 | | | | | | | | | |
| RMI1 | | | | | | | | | | | | | | | | RMI1 | | RMI1 |
| RPA1 | | | | RPA1 | | | RPA1 | | | RPA1 | | | | | | RPA1 | | |
| RPA2 | | | | *RPA2* | | RPA2 | *RPA2* | | RPA2 | *RPA2* | | RPA2 | | | | *RPA2* | | RPA2 |
| RPA3 | | | | | | RPA3 | | | RPA3 | | | RPA3 | | | | | | RPA3 |
| SHFM1 | | | | | | | | | | | | SHFM1 | | | | | | |
| SLX4 | | | | | | | | | | | | | | | | *SLX4* | | SLX4 |
| SSBP1 | | | | | | | | SSBP1 | | | SSBP1 | | | | | | | |
| TOP3A | | | | | | | | | | TOP3A | TOP3A | TOP3A | | | | TOP3A | TOP3A | TOP3A |
| TOP3B | | | | | | | | | | | TOP3B | TOP3B | | | | | TOP3B | TOP3B |
| TOPBP1 | | | | | | | | | | | | TOPBP1 | | | | | | |
| UNG | | | UNG | | | | | | | | | | | | | | | |
| XPA | | | | XPA | XPA | XPA | | | | | | | | | | | | |
| XPC | | | | XPC | | XPC | | | | | | | | | | | | |
| XRCC1 | *XRCC1* | XRCC1 | XRCC1 | | | | | | | | | | | | | | | |
| XRCC5 | | | | | | | | | | | | | XRCC5 | XRCC5 | XRCC5 | | | |
| XRCC6 | | | | | | | | | | | | | XRCC6 | XRCC6 | XRCC6 | | | |

**Supp Figure 9** Genes of major DNA repair pathways of DNA damages caused by IR are upregulated in all three species studies.

**(a)** g:Profiler analysis of differentially expressed genes in *A. antarcticus* after IR.
**(b)** g:Profiler analysis of differentially expressed genes in *P. fairbanksi* after IR.
**(c)** Schematic representation of DNA repair genes up- or down-regulated in *H. exemplaris* after IR. Genes in green colored boxes are upregulated with adjusted p-value < 0.05. Genes in red colored boxes are downregulated with adjusted p-value < 0.05. Genes with no homolog identified in *H. exemplaris* genome are checked with a black cross.
**(d)** Table of DNA repair genes up- or down-regulated in *H. exemplaris, A. antarcticus,* or *P. fairbanksi* after IR, classified based on the KEGG database. Note that the alternative end joining pathway, also called the Micro-homology-Mediated End-Joining (MMEJ) pathway is not currently included in the KEGG database. In the KEGG database, the POLQ gene is included in the BER pathway only. Only genes showing differential gene expression with adjusted p value < 0.05 are shown.

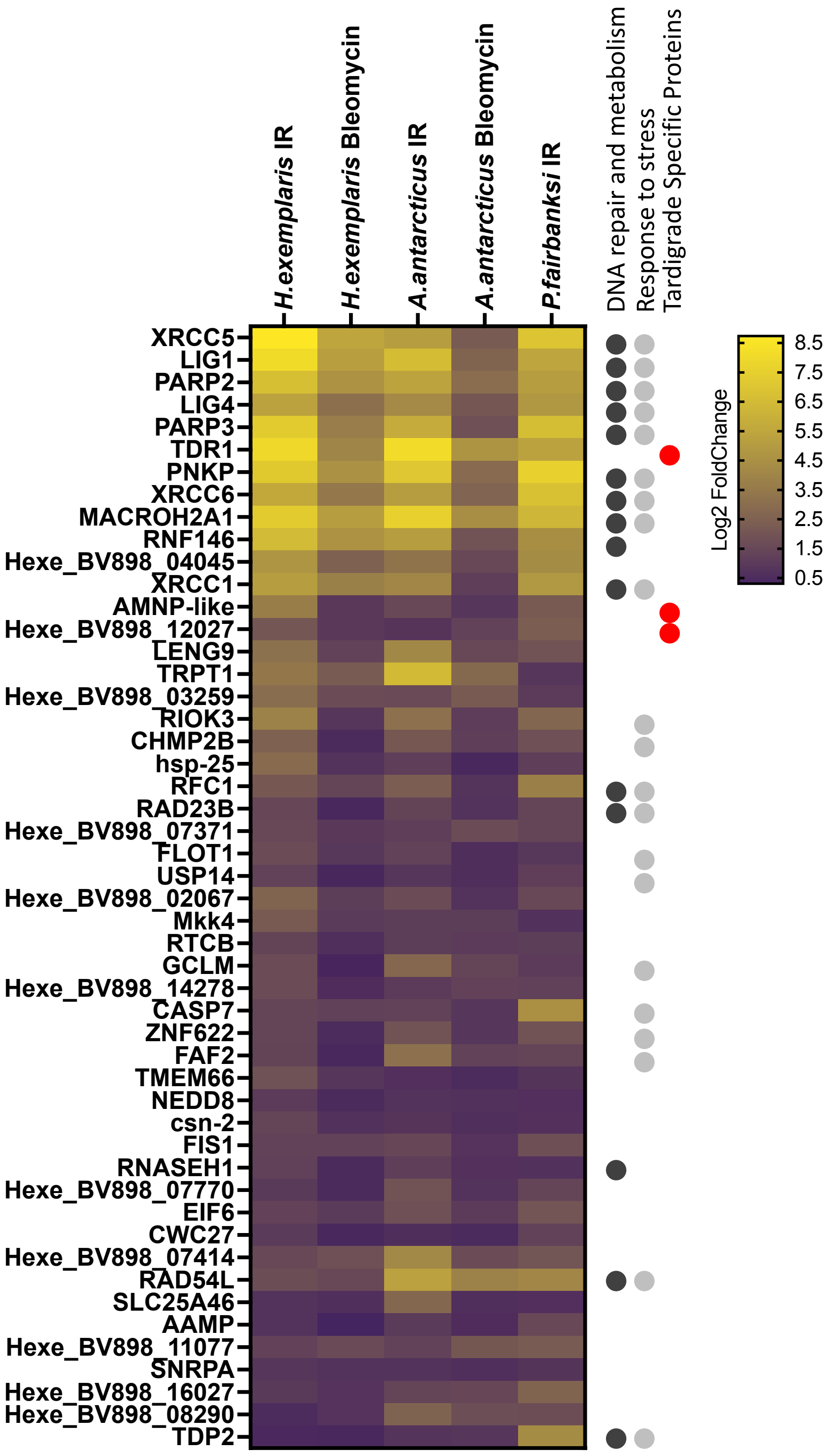


**Supp Figure 10** Heatmap of 50 putative orthologous genes upregulated in response to IR in all the 3 species analyzed, *H. exemplaris*, *A. antarcticus* and *P. fairbanksi,* and in response to Bleomycin in both *H. exemplaris* and *A. antarcticus.*

Heat map of Log2FoldChange (ordered by adjusted p-value) is represented. On the right, information on gene annotation is given. KEGG annotation groups are indicated as well as Tardigrade specific proteins as found in this work (see Material and Methods section for the criterion used to define tardigrade specific genes/proteins). 14/50 genes are DNA repair genes and the other commonality is that 21/50 genes are stress response genes.

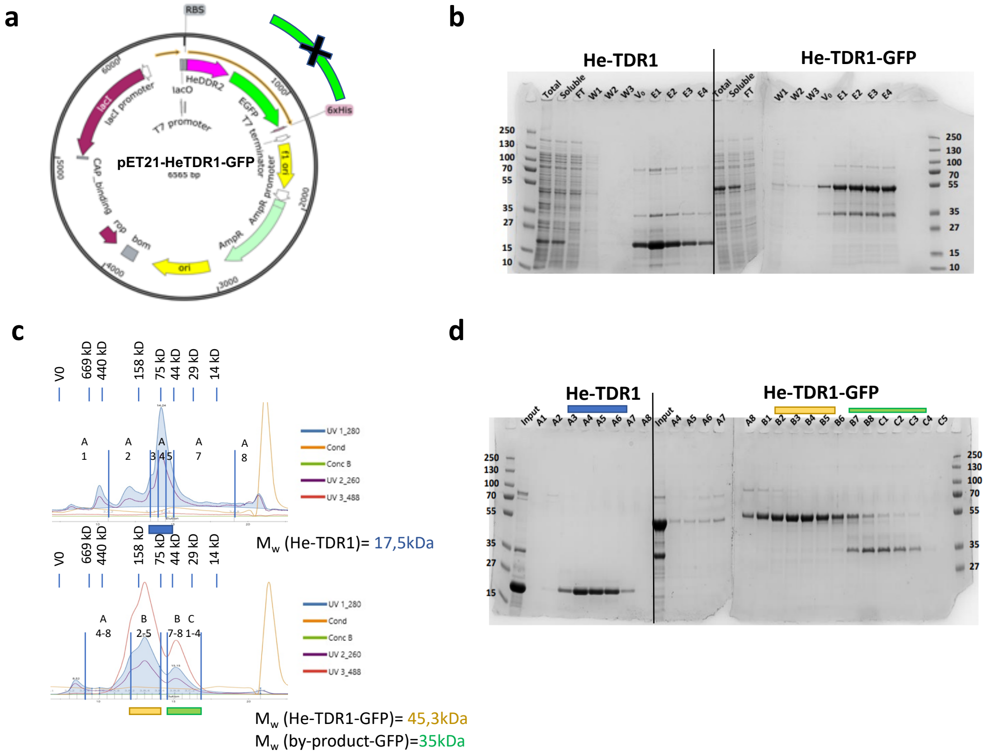


**Supp Figure 11** Production of recombinant *He*-TDR1 and *He*-TDR1-GFP.

**(a)** Map of vector for expression in *E.coli* Rosetta2(DE3): *He*-TDR1 (Mw= 17,5kDa) and its GFP fusion (Mw= 45,3KDa) are His-tagged at their C-terminal end for affinity purification on Nickel Sepharose beads.

**(b)** SDS-PAGE after His-TRAP Sepharose beads chromatography (Instant blue stained). Both proteins are soluble. Fraction V0 to E4 were collected and concentrated on 10kDa MWCO Amicon for gel filtration.

**(c)** Chromatogram (UV absorbance at 260nm, 280nm and 488nm for GFP).

**(d)** SDS-PAGE of indicated fractions obtained after Superdex 200 Increase gel filtration for *He*-TDR1 (C upper panel) and *He*-TDR1-GFP (C bottom panel). V0 and Mw calibration protein volume are reported for comparison to the apparent Mw of *He*-TDR1 (purified protein peak shown in blue) and He-TDR1-GFP (purified protein peak shown in yellow). By-product GFP protein is also purified (shown in green) and is used as internal Mw control. Apparent Mw of *He*-TDR1 and its GFP fusion are higher than expected and suggest a possible multimerization of the *He*-TDR1 protein.

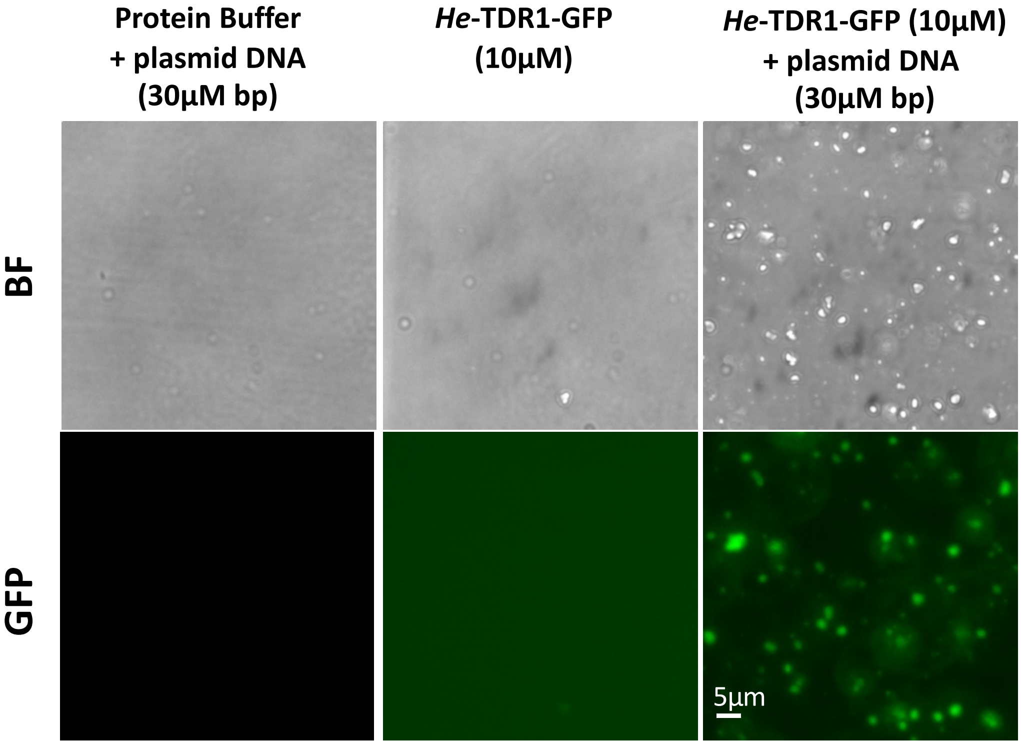


**Supp Figure 12** Formation of aggregates of *He*-TDR1 and DNA.

Bright-field and GFP fluorescence imaging of plasmid DNA 10µM in bp (5900pb) incubated in presence of 10µM of He-TDR1-GFP (10 min at 25°C in 10mM Tris-HCl pH8, NaCl 150mM, Glycerol 10%, 1mM TCEP). No aggregates are observed with the DNA or the protein alone. Large aggregates of DNA with the protein are observed both in bright field and in fluorescence imaging as a result of the binding of He-TDR1-GFP. (similar aggregates were observed in bright-field for He-TDR1 (not shown)).

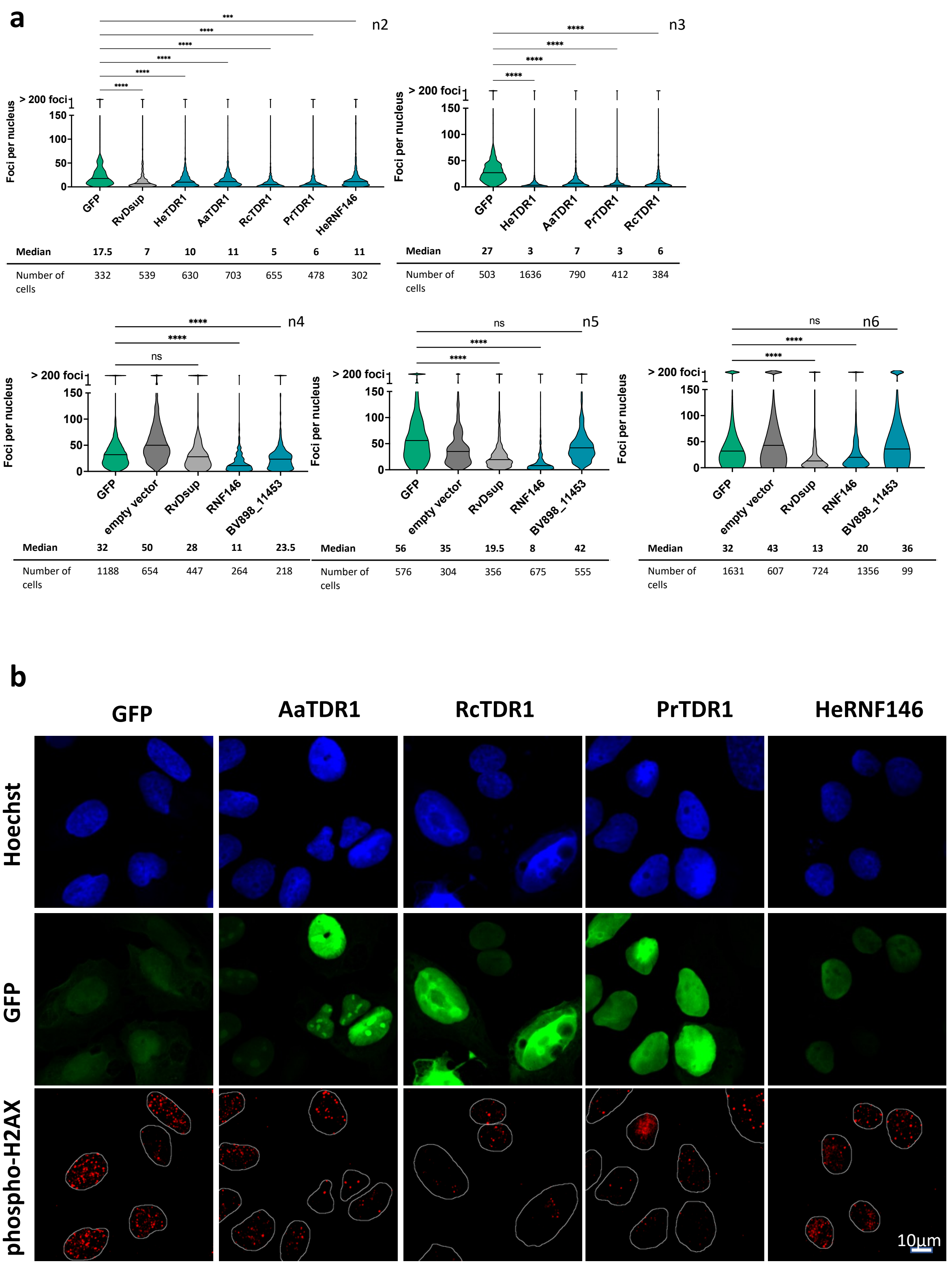


**Supp Figure 13** Independent replicates of experiments of Figure 6.

(a) Number of phospho-H2AX foci in U2OS GFP cells for each indicated Tardigrade gene, empty vector or GFP control plasmid as described in legend to Figure 6. ns, non-significative

(b) Representative images of the experiment presented in Figure 6 (complementing images shown in Figure 6b).

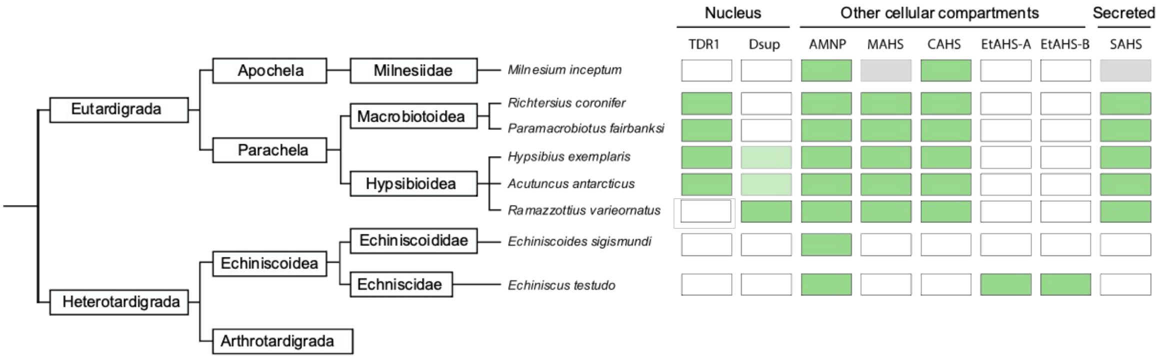


**Supp Figure 14** Phylogenomics of tardigrade specific genes involved in resistance to desiccation and DNA damages (adapted from Arakawa, 2022).
Green and white boxes indicate presence and absence, respectively, of the indicated gene or gene family as found in (Arakawa, 2022) and in this work for TDR1. Light green indicates presence of potential RvDsup ortholog with hypothetical function in radio-resistance (Arakawa, 2022).

A TDR1 homolog could not be identified by Blast analysis of *R. varieornatus* genome and available transcriptomics data. Sequence similarity of a potential TDR1 protein in *R. varieornatus* may be too low and indicate alternative mechanisms of radio-resistance in *R. varieornatus*, for example based on stronger activity of the *Rv*-Dsup compared to *He*- and *Aa*-Dsup. Investigation in additional species may help to clarify the presence/absence of TDR1 in the Ramazzottius genus.

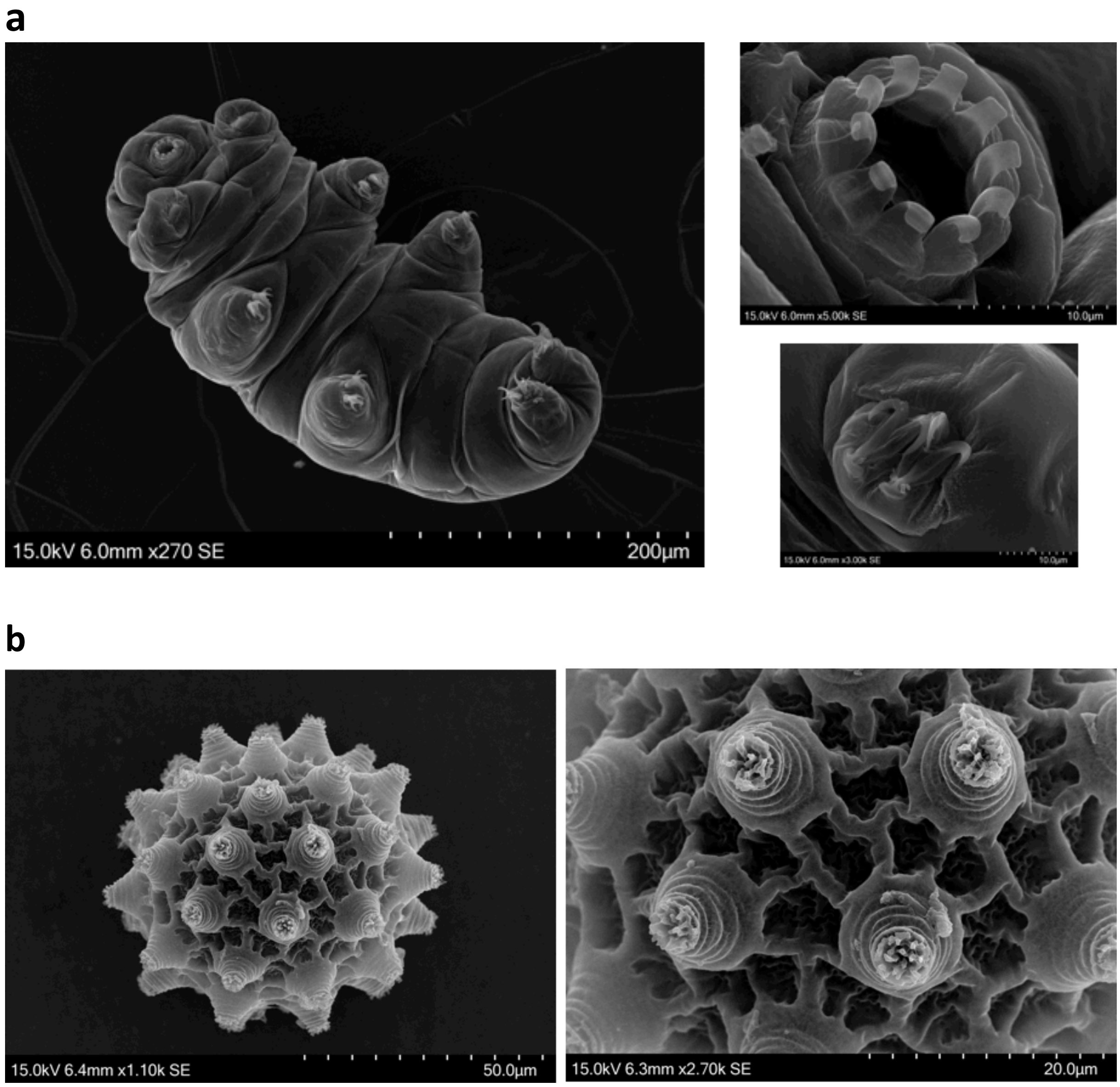


**Supp Figure 15** Identification of *P. fairbanksi* tardigrades isolated and reared from moss garden.

(**a**) SEM of adult specimen with magnification of mouth and claws.
**(b)** SEM of egg with magnification of characteristic spikes decorating the egg surface.
Bright-field morphological analysis performed in parallel by one of the co-authors (R. Guidetti) confirmed *P. fairbanksi* identification.
Species identification was further confirmed by 28S, 18S, COX1, ITS2 sequencing (see next page). For further information on *P. fairbanksi*, see https://doi.org/10.21203/rs.3.rs-2736709/v2
.

```
>Pdec_18S
ATCTGATCGGAAACAGAACTGAACGCAGTTCAAGAGCGCTCAGGATTACTTGTAGCGTGCAAGTATATATCCTGACTCGA
TACGTGCTCGTTGGTATGGTATGCATTCTCTCGATCGCTTGACGGGGTATGAATTGCGAGCCATCTGTGCCGCTGCGCCA
GAAGGTAAGCACAAACCGCATATCGCTGTAAACCTGGCCAGCGCAGTGAAGCTGGTGTCTAGGATACTGTCGATGGTTTG
TTCTAACACTCCTACTGCGTCGAAGTTGCAGCAGCGCTTACGCGCTTACCGACCGATGTAGTACGTGGACAGGTACGCGT
TTCTCGCTTGTTATGCGTTGTGTAGCGTGCACAGCGGACAACGATGGAACAGTATCCGATAAAATGTCCAATGATTGCTA
```

CCTGGTTGATCCTGCCAGTAGTCATATGCTTGTCTCAAAGATTAAGCCATGCATGTCTCAGTACTTGCTATCACAAGGCG
AAACCGCGAATGGCTCATTAAATCAGTTATGGTTCACTAGATCGTACCGTTTACACGGATAACTGTGGTAATTCTAGAGC
TAATACGTGCAACCAGCTCGTTTCCTCGTGAAGCGAGCGCAGTTATTAGAACAAGACCAATCCGGCCTTCGGGTCGGTAA
CTTGGTGACTCTGAATAACCGAAGCAGAGCGCATGGTCTCGTACCGGCGCCAGATCTTTCAAGTGTCTGACTTATCAGCT
TGTTGTTAGGTTATGTTCCTAACAAGGCTATTACGGGTGACGGGGTATCAGGGTCCGATACCGGAGAGGGAGCCTGAGAA
ACGGCTACCACATCCAAGGAAGGCAGCAGGCGCGCAAATTACCCACTCCCAGCACGGGGAGGTAGTGACGAAAAATAACG
ATGCGAGGGCTTTATGCCTCTCGCAATCGGAATGGGTACACTTTAAATCCTTTAACGAGGATCTATTGGAGGGCAAGTCT
GGTGCCAGCAGCCGCGGTAATTCCAGCTCCAATAGCGTATATTAAAGTTGCTGCGGTTAAAAAGCTCGTAGTTGAATCTG
GGCAGTTGGACGGATGGTGCGCTTCACAGCGCTACTGTCTGCTCGGCGCCACAAGCCGGCCATGTCTTGCATGCCCTTCA
CTGGGTGTGCTTGGCGACCGGAACGTTTACTTTGAAAAAATTAGAGTGCTCAAAGCAGGCGTACGGCCTTGCATAATGGT
GCATGGAATAATGGAATAGGACCTCGGTTCTATTTTGTTGGTTTTCGGAACTCGAGGTAATGATTAATAGGAACAGACGG
GGGCATTCGTATTGCGGCGTTAGAGGTGAAATTCTTGGATCGTCGCAAGACGAACTACTGCGAAGGCATTTGCCAAGAAT
GTTTTCATTAATCAAGAACGAAAGTTAGAGGTTCGAAGGCGATCAGATACCGCCCTAGTTCTAACCATAAACGATGCCAA
CCAGCGATCCGTCGGTGTTTATTTGATGACTCGACGGGCAGCTTCCGGGAAACCAAAGTGCTTAGGTTCCGGGGGAAGTA
TGGTTGCAAAGCTGAAACTTAAAGGAATTGACGGAAGGGCACCACCAGGAGTGGAGCCTGCGGCTTAATTTGACTCAACA
CGGGAAAACTTACCCGGCCCGGACACTGTAAGGATTGACAGATTGAGAGCTCTTTCTTGATTCGGTGGGTGGTGGTGCAT
GGCCGTTCTTAGTTGGTGGAGCGATTTGTCTGGTTAATTCCGATAACGAACGAGACTCTAGCCTGCTAAATAGCCAACTG
ATCCGCAGCGTCAGTTGCTACAAAAGCTTCTTAGAGGGACAGGCGGCGTTTAGTCGCACGAGATTGAGCAATAACAGGTC
TGTGATGCCCTTAGATGTCCGGGGCCGCACGCGCGCTACACTGAAGGGAGCAACGTGCTTAATCACCTTGGCCGGAAGGC
CTGGGGAATCCGATTAAACCCCTTCGTGATTGGGATTGAGCTTTGTAATTATCGCTCATGAACGAGGAATTCCCAGTAAG
CGCGAGTCATAAGCTCGCGTTGATTACGTCCCTGCCCTTTGTACACACCGCCCGTCGCTACTACCGATTGAATGATTTAG
TGAGGTCTTCGGACTGGCCATCGAGGCTGCCGCAAGGCGGCTTCGTCTGGTTGGGAAGACGACCAAACTGGCTCATTTAG
AGGAAGTAAAAGTCGTAACAAGGTTTCCGTAGGTGAACCTGCGGAAGGATCATTAACGTGTAATCGGGCGGCGTGTGCCT
GTGCGCTAGGTCTTCGGGCCTGGCCGGGTGCCGTGCTCGTACGCGATAAGCGCTTCTAGCGCTTAACCTGAAAGTCGAGT
CTACACGAACGGTGACTCGCCTATCGTTACTATCCGGAGCGTTTGGTCAGCAATCGCAGCGTGCTTTGCCGACTGCCGGG
GTACGGTAGTGAACTGGTAACTTCGGTACACCGTCGCATGTTGATTTACCTGTCGGAGCGTGAACGGGTGGTCGACACTG
TAATGGCTGTGCCACGGGATACCTACCGTTCTACAAGACGATGCTCTGCGGTCGCTTGAATAACTCACGCGAGACCGTGG
AGAGGCATTCGCCGTAGAATGAGGGCCAGCAGCGCCCAAGGGTATAAAGTAGCTGCAACGAGCGTCGTTGCCGGTTATAT
CCACGATAACGCATCTGTCGTGGGGTATATACCGAATTGGCCGTCCCGCTGCATTTC

>Pdec_ITS2
TCAAAGTGCTTTTCAATTTTCCCTCACGGTACTTGTTCGCTATCGGACTCGTGGTCATATTTAGCCTTCGATGGAGTTTA
CCACCGACTTCACCTTGTACTCACAAACAAGGCGACTCTCCGAAGATGTACACCGGCATCAACTGCTCACGATACGGGCC
TTGCACCCTCCATGGGATGGAGCCTCACTCAGGAGAACTTACGCAAGCAAGCCACACCGGTAACCACCTTCTTCACGCCA
CAGTTCCTGTCACCGTTACCAGCAACAGGATTCGGCGCTGGGCTGTTCCCCTTTCACTCGCAGTTACTATGGGAATCCCC
GTTGGTTTCTTTTCCTCCGCTTAGTAATATGCTTAAGTTCAGCGGGTAATCTCGTCTGAGCTGAGGTCAAAAGATGAGTG
TAGCGAAGCGCAAGTGATTCTTCACGGCCCTGTTCTACAGTTTACTTGCTCCGTAGATTGGGACCTGTTCACCGCTCAAC
GCATAGTTGGCACAGATACGCCACGAACGTGTTCTGTTTTGCTGAACCACTGAAGTGCCGTTGGCAGCTAATCGATACAC
GTGCAACTGCGGCGCACGGCTTCGACAGCCTTCTGCTCCAACTAGCAACACCAGCATTTACCGGTCAGCTAACGAGAACG
CGTATCTGGTCTAGATTTCATCTGATAGCGCCTTATGCAAGCGCTAGCCAGACAATCCGTAGCTTTCACGCAGCTACGAT
TAGTTTTAATCAACTGACCCTCAACCAGGCGTGGCTTCAGTTAACCCGAAGCCGCAATGTGCGTTCGAAGATCCGACGTT
CACAAAGTCCTGCAATTCACGTCGCGTCTCGCATTTTGCTGCGTTCTTCATCGACCCACGAGCCGAATGATCCACCGCAC
AGAGTGATTCATCGGTTTGCATTTTTCGGCTGCTGCTGGGCTGCTGTCTTGCTGACGAGCGCGCCACAACATCAAGCCTC
AAATAGAATACCTTGATACAAATAACGCATTTCCGAAGAAATGCAGCGGGACGGCCAATTCGGTATATACCCCACGACAG
ATGCGTTATCGTGGATATAACCGGCAACGACGCTCGTTGCAGCTACTTTATACCCTTGGGG

>Pdec_COI
CTGCGATGATTTTTCTCTACAAACCACAAAGATATTGGACTCTCTATTTTATTTTTGGGCTTTGGGCAGCCACCATTGG
GACCTCTTTGAGATTTATTATCCGATCTGAATTAAGCCAACCTGGACAATTGTTTGCAGACGAGCAATTATTTAATGTTA
CAGTAACAAGACATGCCTTTGTTATAATTTTCTTTTTTGTGATACCTATTCTTATTGGGGGGTTTGGTAACTGATTGGTC
CCCCTCATAATTGGGGCTCCAGATATGGCTTTTCCTCGAATAAACAACTTAAGATTTTGACTCCTGCCTCCTTCTTTTCT
TCTTATCCTTATGGGAACAATGGCAGAACAAGGGGCGGGTACTGGATGAACTGTTTACCCGCCACTCTCCCACTATTTTG
CTCATAGCGGCCCTAGGGTTGACCTAACAATTTTTTCTCTTCATATCGCCGGAGCATCTTCTATTTTAGGAGCTATTAAT
TTTATTACTACAATTCTTAATATACGATCTTATTCTATAAGAATAGAGCAAATACCTTTATTTGTATGGTCAGTGCTTAT
CACCGCTATTTTACTCCTTTTAGCTCTACCCGTTTTAGCTGGGGCTATTACTATACTACTTCTAGACCGAAATTTTAATA
CTTCTTTTTTTGACCCAGCAGGAGGGGGGGACCCTATTTTATACCAGCATCTGTTTTGATTCTTTGGCCACCCAGAGGTC
TACATTCTAATTCTTCCGGGATTTGGTATTATTTCTCAAGTTATTATCCACTTTAGAGGAAAGTCACTAACATTTGGACA
TTTGGGTATAATTTATGCAATAAGAACAATCGGCCTATTGGGATTTATTGTGTGAGCACACCATATGTTCACAGTAGGTA
TAGACTTAGATACCCGTGCATACTTTACAGCCGCCACTATAATTATTGCCATTCCTACAGGTGTAAAAGTTTTTAGATGA
CTAAGAACAATTTACGGAAGAAAAATTACATTTAGGGCCCCGATATGATGAGCCCTGGGATTTATTTTCCTTTTTACCCT
GGGAGGACTGACAGGGATTGTGTTATCAAATTCAAGAATTGATATTGCTCTCCATGATACTTACTACGTGGTCGCCCACT
TTCACTACGTCCTGTCTATAGGAGCAGTTTTTGCAATTATTTGCGGGGTAGCTCACTGATTCCCTCTTTTGATAGGGGTT
CAAATGAACAATAAATGACTCCAATCCCAGTTTTTGATTATATTTATTGGGGTGAATATAACCTTTTTCCCTCAACATTT
TCTAGGGTTGGCCGGCATACCACGACGATATGTAGATTACCCAGACACCTTTTTTTCGTGGAACATGGCCTCTTCTTTTG
GGTCCTTATTATCAGCACTCTCTGTTATTTTTCTTTTTTTTATTCTATGAGAAGCAATTGTTTCACAACGTTCAACGTAC
CCGGTGTA

>Pdec_28S
GGATAACTGGTTAATACCAGAACGACGATCAGTACATCTGATCGGAAACAGAACTGAACGCAGTTCAAGAGCGCTCAGGA
TTACTTGTAGCGTGCAAGTATATATCCTGACTCGATACGTGCTCGTTGGTATGGTATGCATTCTCTCGATCGCTTGACGG

GGTATGAATTGCGAGCCATCTGTGCCGCTGCGCCAGAAGGTAAGCACAAACCGCATATCGCTGTAAACCTGGCCAGCGCA
GTGAAGCTGGTGTCTAGGATACTGTCGATGGTTTGTTCTAACACTCCTACTGCGTCGAAGTTGCAGCAGCGCTTACGCGC
TTACCGACCGATGTAGTACGTGGACAGGTACGCGTTTCTCGCTTGTTATGCGTTGTGTAGCGTGCACAGCGGACAACGAT
GGAACAGTATCCGATAAAATGTCCAATGATTGCTACCTGGTTGATCCTGCCAGTAGTCATATGCTTGTCTCAAAGATTAA
GCCATGCATGTCTCAGTACTTGCTATCACAAGGCGAAACCGCGAATGGCTCATTAAATCAGTTATGGTTCACTAGATCGT
ACCGTTTACACGGATAACTGTGGTAATTCTAGAGCTAATACGTGCAACCAGCTCGTTTCCTCGTGAAGCGAGCGCAGTTA
TTAGAACAAGACCAATCCGGCCTTCGGGTCGGTAACTTGGTGACTCTGAATAACCGAAGCAGAGCGCATGGTCTCGTACC
GGCGCCAGATCTTTCAAGTGTCTGACTTATCAGCTTGTTGTTAGGTTATGTTCCTAACAAGGCTATTACGGGTGACGGGG
TATCAGGGTCCGATACCGGAGAGGGAGCCTGAGAAACGGCTACCACATCCAAGGAAGGCAGCAGGCGCGCAAATTACCCA
CTCCCAGCACGGGGAGGTAGTGACGAAAAATAACGATGCGAGGGCTTTATGCCTCTCGCAATCGGAATGGGTACACTTTA
AATCCTTTAACGAGGATCTATTGGAGGGCAAGTCTGGTGCCAGCAGCCGCGGTAATTCCAGCTCCAATAGCGTATATTAA
AGTTGCTGCGGTTAAAAAGCTCGTAGTTGAATCTGGGCAGTTGGACGGATGGTGCGCTTCACAGCGCTACTGTCTGCTCG
GCGCCACAAGCCGGCCATGTCTTGCATGCCCTTCACTGGGTGTGCTTGGCGACCGGAACGTTTACTTTGAAAAAATTAGA
GTGCTCAAAGCAGGCGTACGGCCTTGCATAATGGTGCATGGAATAATGGAATAGGACCTCGGTTCTATTTTGTTGGTTTT
CGGAACTCGAGGTAATGATTAATAGGAACAGACGGGGGCATTCGTATTGCGGCGTTAGAGGTGAAATTCTTGGATCGTCG
CAAGACGAACTACTGCGAAGGCATTTGCCAAGAATGTTTTCATTAATCAAGAACGAAAGTTAGAGGTTCGAAGGCGATCA
GATACCGCCCTAGTTCTAACCATAAACGATGCCAACCAGCGATCCGTCGGTGTTTATTTGATGACTCGACGGGCAGCTTC
CGGGAAACCAAAGTGCTTAGGTTCCGGGGGAAGTATGGTTGCAAAGCTGAAACTTAAAGGAATTGACGGAAGGGCACCAC
CAGGAGTGGAGCCTGCGGCTTAATTTGACTCAACACGGGAAAACTTACCCGGCCCGGACACTGTAAGGATTGACAGATTG
AGAGCTCTTTCTTGATTCGGTGGGTGGTGGTGCATGGCCGTTCTTAGTTGGTGGAGCGATTTGTCTGGTTAATTCCGATA
ACGAACGAGACTCTAGCCTGCTAAATAGCCAACTGATCCGCAGCGTCAGTTGCTACAAAAGCTTCTTAGAGGGACAGGCG
GCGTTTAGTCGCACGAGATTGAGCAATAACAGGTCTGTGATGCCCTTAGATGTCCGGGGCCGCACGCGCGCTACACTGAA
GGGAGCAACGTGCTTAATCACCTTGGCCGGAAGGCCTGGGGAATCCGATTAAACCCCTTCGTGATTGGGATTGAGCTTTG
TAATTATCGCTCATGAACGAGGAATTCCCAGTAAGCGCGAGTCATAAGCTCGCGTTGATTACGTCCCTGCCCTTTGTACA
CACCGCCCGTCGCTACTACCGATTGAATGATTTAGTGAGGTCTTCGGACTGGCCATCGAGGCTGCCGCAAGGCGGCTTCG
TCTGGTTGGGAAGACGACCAAACTGGCTCATTTAGAGGAAGTAAAAGTCGTAACAAGGTTTCCGTAGGTGAACCTGCGGA
AGGATCATTAACGTGTAATCGGGCGGCGTGTGCCTGTGCGCTAGGTCTTCGGGCCTGGCCGGGTGCCGTGCTCGTACGCG
ATAAGCGCTTCTAGCGCTTAACCTGAAAGTCGAGTCTACACGAACGGTGACTCGCCTATCGTTACTATCCGGAGCGTTTG
GTCAGCAATCGCAGCGTGCTTTGCCGACTGCCGGGGTACGGTAGTGAACTGGTAACTTCGGTACACCGTCGCATGTTGAT
TTACCTGTCGGAGCGTGAACGGGTGGTCGACACTGTAATGGCTGTGCCACGGGATACCTACCGTTCTACAAGACGATGCT
CTGCGGTCGCTTGAATAACTCACGCGAGACCGTGGAGAGGCATTCGCCGTAGAATGAGGGCCAGCAGCGCCCAAGGGTAT
AAAGTAGCTGCAACGAGCGTCGTTGCCGGTTATATCCACGATAACGCATCTGTCGTGGGGTATATACCGAATTGGCCGTC
CCGCTGCATTTCTTCGGAAATGCGTTATTTGTATCAAGGTATTCTATTTGAGGCTTGATGTTGTGGCGCGCTCGTCAGCA
AGACAGCAGCCCAGCAGCAGCCGAAAAATGCAAACCGATGAATCACTCTGTGCGGTGGATCATTCGGCTCGTGGGTCGAT
GAAGAACGCAGCAAAATGCGAGACGCGACGTGAATTGCAGGACTTTGTGAACGTCGGATCTTCGAACGCACATTGCGGCT
TCGGGTTAACTGAAGCCACGCCTGGTTGAGGGTCAGTTGATTAAAACTAATCGTAGCTGCGTGAAAGCTACGGATTGTCT
GGCTAGCGCTTGCATAAGGCGCTATCAGATGAAATCTAGACCAGATACGCGTTCTCGTTAGCTGACCGGTAAATGCTGGT
GTTGCTAGTTGGAGCAGAAGGCTGTCGAAGCCGTGCGCCGCAGTTGCACGTGTATCGATTAGCTGCCAACGGCACTTCAG
TGGTTCAGCAAAACAGAACACGTTCGTGGCGTATCTGTGCCAACTATGCGTTGAGCGGTGAACAGGTCCCAATCTACGGA
GCAAGTAAACTGTAGAACAGGGCCGTGAAGAATCACTTGCGCTTCGCTACACTCATCTTTTGACCTCAGCTCAGACGAGA
TTACCCGCTGAACTTAAGCATATTACTAAGCGGAGGAAAAGAAACCAACGGGGATTCCCATAGTAACTGCGAGTGAAAGG
GGAACAGCCCAGCGCCGAATCCTGTTGCTGGTAACGGTGACAGGAACTGTGGCGTGAAGAAGGTGGTTACCGGTGTGGCT
TGCTTGCGTAAGTTCTCCTGAGTGAGGCTCCATCCCATGGAGGGTGCAAGGCCCGTATCGTGAGCAGTTGATGCCGGTGT
ACATCTTCGGAGAGTCGCCTTGTTTGTGAGTACAAGGTGAAGTCGGTGGTAAACTCCATCGAAGGCTAAATATGACCACG
AGTCCGATAGCGAACAAGTACCGTGAGGGAAAATTGAAAAGCACTTTGAAGAGAGAGCGAAACAGTGCGTGAAACCGCTC
AGAGGCAAGCAGATGGGGCCTCGAAGGCAAAGCAGTGAATTCAGCCGGTGTGGTGCGTGGCTGGTTGGTGTTGGGATCGC
AAGACCCTAGCTGATTATGCTCGCGTGCTGGCCGGTGCATTTTCGCTGTTTGTACGTCACCGCCGTTGAGCGAGCATCCG
TCGGGTATGCGTGTGAAGCCTTATTCCTTCGGGCGTAGGTGCTTACTGCTAACTTGTACGCGTTTGCGCCTCAACTGGTC
ATGTCAGCGTGTGCCAGCGTTAGCGTTGGGCCGGTATGCTCTGCGGTGTGTTGTGGGATGACGAGCTTGCTCGGCTCCTC
GATACGCAGTGGACTCGTTGCCGGTTTTCAACGTAGGCACATTGTAGATTCGGTGGCGAGTAGACGGCTGCCCATCTAAC
CCGTCTTGWAACACGGACCAAGGAGTTCAACATGCGCGCTAGTTGTTGGGACTTGAAGCCCGCTAGCAAAGTGAAAGCAA
GACACAGTGTACGCTGTGTATTGGCGAGATCCCGTCACCTGGTTTACCAGGCCGGGCGCACCGCCGGCCCGTCAAAAGCT
CATGTGGCTTTGGCGGAGCTTGAGCGTGCACGTTGAGACCCGAAAGATGGTGAACTATGCCTGGGCAGGATGAAGCCAGG
GGAAACTCTGGTGGAGGTCCGTAGCGATTCTGACGTGCAAATCGATCGTCTGACCTGGGTATAGGGGCGAAAGACTAATC
GAACCATCTAGTAGCTGGTTCCCTCCGAAGTTTCCCTCAGGATAGCTGGCACTCGAGAGAACGTAGTCTCTCCCGGTAAA
GCGAATGATTAGTGGCCTTGGGGTCGAAACGACCTTAACCAATTCTCAAACTTTAAATGGGTGAGAAGTCCGGCTTGCTT
AAATGCATAGCTGAAGTCCGGACGTTGGATACGAGCGCCTAGTGGGCCACTTTTGGTAAGCAGAACTGGCGCTGTGGGAT
GAACCGAACGCTGAGTTATGGCGCCCGACGAGACGCTCATCAGATCCCAGAAAAGGTGTTGGTTGCTATAGACAGCAGGA
CGGTGGCCATGGAAGTCGGAACCCGCTAAGGAGTGTGTAACAACTCACCTGCCGAAGCAACTAGCCCTGAAAATGGATGG
CGCTAGAGCGTCTGGCCTATACTCGGCCGTTGCAGCAGCAGCAACGTTAAGTCAAGCTGCAACGAGTAG

**Supp Figure 16** Uncropped images of Western blots from Figures 1 and 3

(a) Uncropped images of Western blots from Figure 1
(b) Uncropped images of Western blots from Figure 1

**a**

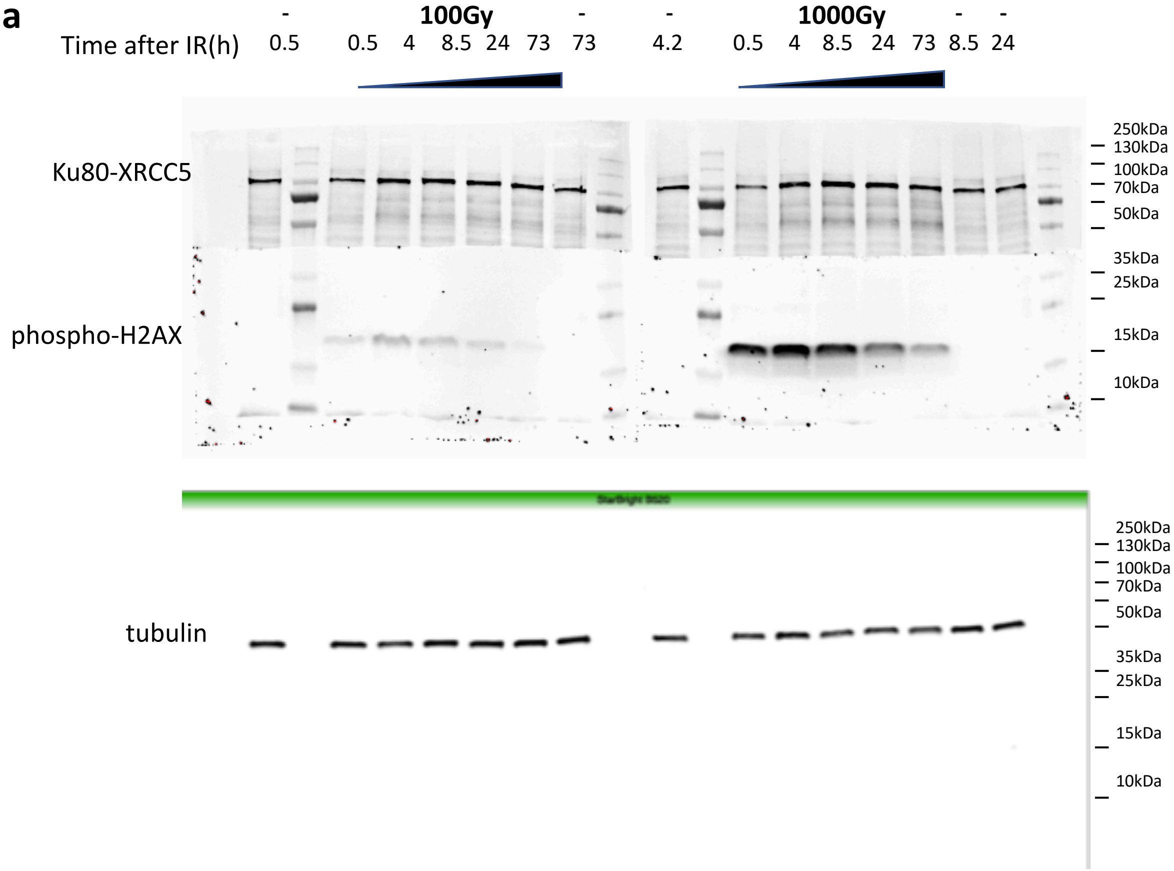

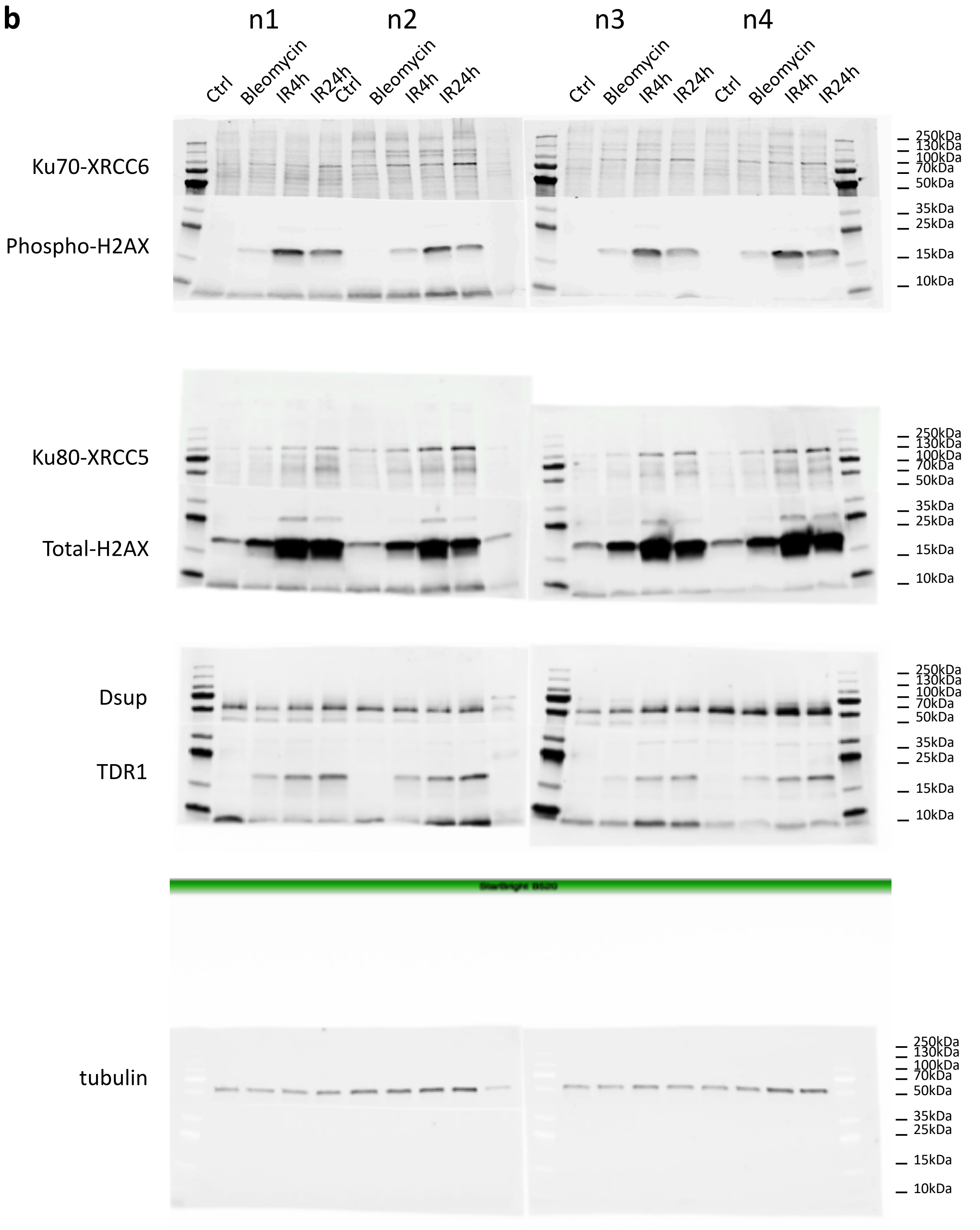
b
n1
n2
n3
n4
Ctrl
Bleomycin
IR4h
IR24h
Ctrl
Bleomycin
IR4h
IR24h
Ctrl
Bleomycin
IR4h
IR24h
Ctrl
Bleomycin
IR4h
IR24h
Ku70-XRCC6
Phospho-H2AX
Ku80-XRCC5
Total-H2AX
Dsup
TDR1
tubulin
250kDa
130kDa
100kDa
70kDa
50kDa
35kDa
25kDa
15kDa
10kDa

**Supp Table 10**: List of Tardigrade specific proteins differentially expressed in all 3 conditions analyzed.

Tardigrade specific proteins are ranked according to the Log2 Fold Change (from highest to lowest) at 4h post-irradiation.

| *Up(+)/Down(-)* | *List of common Tardigrade specific DE protein locus ID (IR4h /IR24h/Bleomycin)* | *NCBI protein ID* |
|---|---|---|
| *+* | BV898_14257 =>TDR1 | OQV11461.1 |
| + | BV898_12300 =>AMNP-like | OQV13448.1 |
| + | BV898_14013 | OQV11669.1 |
| + | BV898_10262 =>AMNP-like | OQV15540.1 |
| + | BV898_06508 | OQV19521.1 |
| *+* | *BV898_10264=>AMNP*-like | *OQV15542.1* |
| + | BV898_06926 | OQV19074.1 |
| + | BV898_01539 | OQV24477.1;OQV24476.1 |
| - | BV898_17845 | OWA53415.1 |
